# Flexible Multi-Step Hypothesis Testing of Human ECoG Data using Cluster-based Permutation Tests with GLMEs

**DOI:** 10.1101/2023.03.31.535153

**Authors:** Seth D König, Sandra Safo, Kai Miller, Alexander B. Herman, David P. Darrow

## Abstract

**Background:** Time series analysis is critical for understanding brain signals and their relationship to behavior and cognition. Cluster-based permutation tests (CBPT) are commonly used to analyze a variety of electrophysiological signals including EEG, MEG, ECoG, and sEEG data without *a priori* assumptions about specific temporal effects. However, two major limitations of CBPT include the inability to directly analyze experiments with multiple fixed effects and the inability to account for random effects (e.g. variability across subjects). Here, we propose a flexible multi-step hypothesis testing strategy using CBPT with Linear Mixed Effects Models (LMEs) and Generalized Linear Mixed Effects Models (GLMEs) that can be applied to a wide range of experimental designs and data types.

**Methods:** We first evaluate the statistical robustness of LMEs and GLMEs using simulated data distributions. Second, we apply a multi-step hypothesis testing strategy to analyze ERPs and broadband power signals extracted from human ECoG recordings collected during a simple image viewing experiment with image category and novelty as fixed effects. Third, we assess the statistical power differences between analyzing signals with CBPT using LMEs compared to CBPT using separate t-tests run on each fixed effect through simulations that emulate broadband power signals. Finally, we apply CBPT using GLMEs to high-gamma burst data to demonstrate the extension of the proposed method to the analysis of nonlinear data.

**Results:** First, we found that LMEs and GLMEs are robust statistical models. In simple simulations LMEs produced highly congruent results with other appropriately applied linear statistical models, but LMEs outperformed many linear statistical models in the analysis of “suboptimal” data and maintained power better than analyzing individual fixed effects with separate t-tests. GLMEs also performed similarly to other nonlinear statistical models. Second, in real world human ECoG data, LMEs performed at least as well as separate t-tests when applied to predefined time windows or when used in conjunction with CBPT. Additionally, fixed effects time courses extracted with CBPT using LMEs from group-level models of pseudo-populations replicated latency effects found in individual category-selective channels. Third, analysis of simulated broadband power signals demonstrated that CBPT using LMEs was superior to CBPT using separate t-tests in identifying time windows with significant fixed effects especially for small effect sizes. Lastly, the analysis of high-gamma burst data using CBPT with GLMEs produced results consistent with CBPT using LMEs applied to broadband power data.

**Conclusions:** We propose a general approach for statistical analysis of electrophysiological data using CBPT in conjunction with LMEs and GLMEs. We demonstrate that this method is robust for experiments with multiple fixed effects and applicable to the analysis of linear *and* nonlinear data. Our methodology maximizes the statistical power available in a dataset across multiple experimental variables while accounting for hierarchical random effects and controlling FWER across fixed effects. This approach substantially improves power and accuracy leading to better reproducibility. Additionally, CBPT using LMEs and GLMEs can be used to analyze individual channels or pseudo-population data for the comparison of functional or anatomical groups of data.

**Highlights:**

● Combining CBPT with GLMEs allows statistical analysis to match experimental design.
● CBPT with GLMEs accounts for subject variability and hierarchical random effects.
● The proposed method maintains control of type I error, type II error, and FWER.
● CBPT with GLMEs can be applied to individual channels and pseudo-population data.

## 1.0 Introduction

Event-related analysis–such as the analysis of ERPs in EEG data–is one of the most common forms of analysis for electrophysiological data in cognitive neuroscience due to its tractability. In essence, time series data representing voltage signals recorded from the brain using EEG, MEG, ECoG, sEEG, or microelectrodes are aligned to task and/or behavioral events in order to determine how brain activity correlates with task features or behavioral responses. By doing so neuroscientists hope to uncover how the brain performs specific behavioral and cognitive processes.

Importantly, appropriate statistical analysis is required to accurately and reliably draw conclusions from the event-related data (ERD). To test their hypotheses, neuroscientists employ a variety of analysis techniques. Some predefine a time window after an event on which to run all their analyses. Others may employ mass-univariate tests using cluster-based statistics to empirically derive an estimate of a time window for subsequent analyses (Maris & Oostenveld, 2007). Each method has its advantages and disadvantages. For example, predefining a time window is simple and straightforward, but selecting a time window *a priori* without knowing exactly when an effect will occur could decrease statistical power (type II error). Conversely, defining a window based on certain aspects of the observed data can also lead to “bogus” findings that never existed (Luck & Gaspelin, 2017). Cluster-based permutation tests (CBPT) circumvent these issues by empirically estimating a window without *a* p*riori* knowledge or human bias. However, a major limitation of CBPT analysis is CBPT has weak control of Family Wise Error Rate (FWER) pertaining to localization of effects in time (Maris & Oostenveld, 2007). FDR methods–such as the Benjamini-Hochberg procedure–can be used instead of CBPT methods when localization is critical, but FDR methods trade statistical power for better type I error control and thus can be less sensitive to broadly distributed effects commonly seen in neural data (Groppe et al., 2011a, 2011b).

The earliest applications of CBPT in EEG/MEG analysis used t-tests to analyze ERPs in a semantic processing paradigm with a *1×2* factorial experimental design where participants were presented with congruent and incongruent sentences (Maris & Oostenveld, 2007). While others have used ANOVAs to analyze experiments with a signal factor with multiple levels (i.e. *1xn*) (Frossard & Renaud, 2021), many experiments in neuroscience have multiple factors (i.e. multiple fixed effects) often with multiple levels, yet no consistent method has been proposed to analyze these data. Importantly, analyzing multiple fixed effects *separately* (e.g. many simple linear regressions) is statistically less powerful than analyzing multiple fixed effects *together* (e.g. multiple linear regression) because analysis of a single fixed effect ignores potential correlations and interactions between predictors whereas analyzing multiple fixed effects with a single model does not (Morrissey & Ruxton, 2018). Furthermore, the use of single-effect models in experiments involving multiple fixed effects implies the *a priori* selection of the “main’’ fixed effect for estimating a time window of interest. However, the neuroscientist is not guaranteed to pick the “correct” fixed effect nor is there any guarantee that other fixed effects could correlate with activity outside this window. Therefore, in order to broaden the utility of CBPT, we propose the use of CBPT with mixed-effect models that account for all fixed effects in a single model in experiments with a multifactor design and account for hierarchical random effects across groups of data (e.g subjects and channels).

We are not the first to propose the use of CBPT with mixed effect models, but to the best of our knowledge, we are the first to propose a general statistical framework for time series analysis using CBPT with mixed effect models integrated with multi-step hypothesis testing strategy. Linear mixed effect models (LMEs)–even with cluster-based statistics–have been used in fMRI analysis for quite some time (G. Chen et al., 2012; Jo et al., 2021; Lindquist et al., 2012; Rodrigue et al., 2018; Winkler et al., 2014, 2016). LMEs have also been used to analyze ERPs in predefined windows (Frömer et al., 2021; Koerner & Zhang, 2017; Tremblay & Newman, 2015). Additionally, CBPT with LMEs (Bianchi et al., 2019) and LMEs applied after CBPT (Frömer et al., 2018) have been used to analyze human EEG data but certainly not to the same extent as the fMRI literature. For example in (Frömer et al., 2018), the authors first used CBPT with t-tests on the largest fixed effect to estimate a good time window for subsequent analysis with an LME, but this separate analysis of fixed effects may have reduced their statistical power in the initial cluster identification process. In (Bianchi et al., 2019) the authors directly used LMEs with CBPT in a method similar to what we propose here; however, the authors did not correct for the number of fixed effects analyzed (two tailed *t*-test, α =0.05, |*t*| > 2), and they analyzed each fixed effect separately which likely increased the overall FWER; this potential increase in FWER likely explains why such a large number of electrodes and time points were considered significant in their study. Generalized linear mixed effect models (GLM[E]s) have also gained popularity in the neuroscience community due to their power to directly analyze nonlinear data. GLM[E]s have been used to analyze reaction times due to the long tails commonly found in such data (Baayen & Milin, n.d.) as well as accuracy and choice behavior (Frömer et al., 2018; Krajbich & Dean, 2015). More complicated linear-nonlinear-Poisson models have also been used to estimate complex firing rate patterns of individual neurons in the presence of large parameter spaces where regularization is necessary (Hardcastle et al., 2017; Yoo et al., 2020).

The use of mixed effects models like LMEs has increased as the neuroscience community has recognized the important benefits of including random effects in statistical models. First, compared to many traditional linear statistical models, LMEs can fit trial-by-trial data, deal with missing data, are more robust to violations of normality (Schielzeth et al., 2020), are less prone to outliers, account for random effects, and are more sensitive to smaller effect sizes. GLMEs offer these same benefits but are designed to fit nonlinear data. Second, we should use statistical models that reflect experimental designs and how the data is collected. In many experiments, the data is collected across channels, sessions, subjects, etc. including in a hierarchical fashion (e.g. channel is nested within-subject); therefore statistical models should reflect the pattern of random effects in the data (Yu et al., 2022). Third, while mixed-effects models often produce similar beta weights estimates compared to fixed effects-only models, the confidence intervals around the beta weights (and the standard error) can change in meaningful ways, especially in the presence of unbalanced random effects (i.e. unequal observations across groups) (Nikolakopoulou et al., 2014; Pol et al., 2009; Yang & Land, 2008). Combining these benefits of LMEs explains why in many cases where comparisons were made, LMEs outperformed simpler linear models such as ANOVAs in the analysis of EEG data (Heise et al., 2022). Moreover, the use of LMEs is associated with trivial downsides including increased computational time and mixed effects need at least 5 groups for accurate estimation of random intercepts which is often easily met in most experiments (e.g. n > 5 subjects), but more data is required to estimate random slopes. Overall, the many upsides of this generalized framework far outweigh the limited downsides.

In this paper we will demonstrate the feasibility, flexibility, and robustness of CBPT with LMEs and GLMEs through simulation and empirical analysis. In simple simulations we will show that LMEs and GLMEs are at least as good as, if not better than, other commonly used statistical models when analyzing simple data distributions. Further, in simulations of broadband power signals, we will show that analyzing individual fixed effects with CBPT using separate t-tests identifies significantly fewer clusters than CBPT using LMEs. We will also show that LME results from analyzing ERPs and broadband power signals extracted from real-world human ECoG data are similar to or better than results from separate t-tests when applied to predefined time windows or used in conjunction with CBPT. Finally, we will show that CBPT using GLMEs applied to high-gamma burst signals–a form of nonlinear data–produces similar results to CBPT with LMEs applied to broadband power signals.

### 2.0 Materials and Methods

All analyses and simulations were carried out in Matlab with custom written code. Simulation methods and results are described elsewhere in the supplementary materials.

Example Matlab code can be found on Github (https://github.com/hermandarrowlab/GLME-CBPT) and example preprocessed data can be found on Zenodo (https://doi.org/10.5281/zenodo.7703148). The code and data are available to replicate the basic methods and results found in this paper.

### 2.1 Processing and Re-Analysis of Human ECoG Data

Human intracranial ECoG data were obtained from an open-source repository (K. J. Miller, 2019; K. J. Miller et al., 2015, 2016). Epilepsy patients undergoing seizure localization were shown a semi-random sequence of images for 400 ms each with a 400 ms inter-stimulus-interval. The experiment had a 2×2 factorial design: image category and image novelty; participants were shown 50 unique face images and 50 unique house images, and images were shown a total of 3 times: novel and then repeated twice. The full data set contained 714 ECoG channels from 14 subjects. However, we found that two subjects had voltage ranges 2 orders of magnitude greater than the others, so we ignored these data. In total we imported 629 ECoG channels from 12 subjects. Data were sampled at 1000 Hz.

We used custom written code and Fieldtrip functions (Oostenveld et al., 2011) to filter data from 1 Hz to 200 Hz as well as remove line noise using spectral interpolation. Next, we removed 17/629 (2.7%) ECoG channels with outlier power spectra using a custom written algorithm to remove “bad” channels corrupted by things like intermittent line noise. We used common average re-referencing to further reduce noise, remove artifacts, and remove other common signals. Broadband power was calculated from 1 to 200 Hz using the code provided in the data repository (K. J. Miller, 2019; K. J. Miller et al., 2015, 2016). Upon examination of broadband signals, we found and removed 2 outlier ECoG channels with greater than 100 a.u. standard deviation across the whole recording. For each channel broadband signals and ERPs were extracted 200 ms before to 600 ms after image onset. We then de-baselined the ERP and broadband signals by subtracting the mean value from 200 ms before image onset to 100 ms after image onset; baseline correction helped remove a consistent difference in mean values for many channels across novel and repeated blocks.

ERP and broadband power signals were confirmed to be normally distributed using visual inspection of distribution shapes and Q-Q plots (data not shown) although broadband power signals appeared more likely to have outliers and skewed tails. However, we determined that we had sufficiently trial counts and that LMEs could appropriately handle the small amount of outliers and skewness found in the broadband power signals.

We initially replicated the previously published results by analyzing the ERP and broadband power signals using a simple time window-based analysis from 100 ms to 400 ms after image onset. We then compared the results from separate t-tests to LMEs for the ERP and broadband power data. The significance threshold (α) was set at 0.05 but divided by 2 for the number of fixed effects. The main LME formula for the predefined time window analysis was *’erd ∼ imageCategory + novelRepeat + (1|imageID)’*, where *erd* generically stands for event-related data and includes both ERPs and broadband power data (**Table 1**).

**Table 1:**
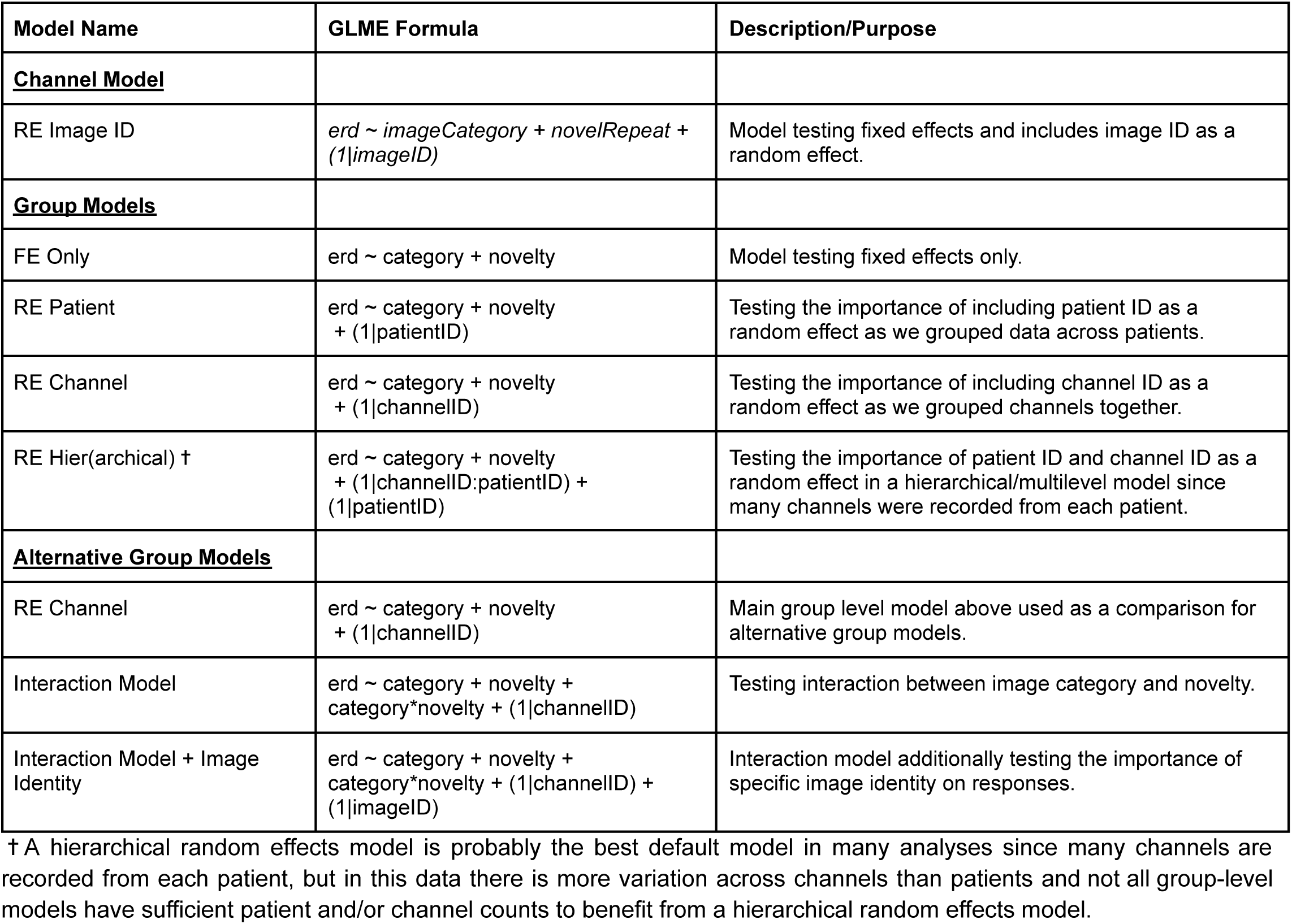
Group-Level Model Description.

| Model Name | GLME Formula | Description/Purpose |
| --- | --- | --- |
| **<u>Channel Model</u>** |  |  |
| RE Image ID | $erd \sim imageCategory + novelRepeat + (1\vert imageID)$ | Model testing fixed effects and includes image ID as a random effect. |
| **<u>Group Models</u>** |  |  |
| FE Only | $erd \sim category + novelty$ | Model testing fixed effects only. |
| RE Patient | $erd \sim category + novelty + (1\vert patientID)$ | Testing the importance of including patient ID as a random effect as we grouped data across patients. |
| RE Channel | $erd \sim category + novelty + (1\vert channelID)$ | Testing the importance of including channel ID as a random effect as we grouped channels together. |
| RE Hier(archical) † | $erd \sim category + novelty + (1\vert channelID:patientID) + (1\vert patientID)$ | Testing the importance of patient ID and channel ID as a random effect in a hierarchical/multilevel model since many channels were recorded from each patient. |
| **<u>Alternative Group Models</u>** |  |  |
| RE Channel | $erd \sim category + novelty + (1\vert channelID)$ | Main group level model above used as a comparison for alternative group models. |
| Interaction Model | $erd \sim category + novelty + category*novelty + (1\vert channelID)$ | Testing interaction between image category and novelty. |
| Interaction Model + Image Identity | $erd \sim category + novelty + category*novelty + (1\vert channelID) + (1\vert imageID)$ | Interaction model additionally testing the importance of specific image identity on responses. |

† A hierarchical random effects model is probably the best default model in many analyses since many channels are recorded from each patient, but in this data there is more variation across channels than patients and not all group-level models have sufficient patient and/or channel counts to benefit from a hierarchical random effects model.

Congruence was defined as the percent of comparisons with the same statistical results (p < α?). Congruence is similar to accuracy calculated from a confusion matrix and is the sum of the diagonal divided by the number of comparisons (100 * (TP + TN)/N), but there is no gold standard statistical model. Instead, for all empirical and simulation analyses, we compared each statistical model results to the LME or GLME results.

Similarly, statistical power was defined based on the relative proportion of significant results of a given statistical model compared to the LME or GLME results. Thus, if there were significantly fewer significant results from a given statistical model compared to the LME or GLME results (i.e. χ^2^ test, p < α), then we considered this to be a significant reduction in statistical power; this is equivalent to a significant increase in the false negative (FN) rate.

### 2.2 Analysis of Broadband Power and ERP Data using Cluster-based Permutation testing with LMEs

#### 2.2.1 Cluster Based Permutation Tests with LMEs

CBPT using LMEs was performed similarly to CBPT using t-tests (Maris & Oostenveld, 2007) with some modifications to adjust for the flexibility and potential complexity of LMEs:

1. For each time sample, fit the data with the desired LME formula.
2. Select all time samples for which at least one fixed effect is significant at some threshold α, Bonferroni corrected for the number of fixed effects (i.e. p < α/#_FE_).

a. Each interaction term is also considered in the Bonferroni correction.
3. Cluster all significant time samples in the previous step based on temporal adjacency.
4. Remove clusters smaller than a certain length in time (*t*_min_).
5. For each cluster calculate the mean response across samples for each trial, and then fit the mean response across trials with the desired LME formula above. From this cluster-level LME, calculate the sum(*t-statistic*^2^) for the beta weights across all fixed effects ignoring the intercept. This is the cluster-level statistic for that cluster.
6. Identify the cluster with the largest cluster-level statistic. Permute the data (from step 5) for the largest cluster *N* times and calculate a shuffled sum(*t-statistic*^2^) distribution.
7. A cluster is considered significant if the cluster-level statistic for that cluster exceeds the 100*(1-α)-percentile of the shuffled sum(*t-statistic*^2^) distribution from the largest cluster.

In Matlab, we used the function *fitlme()* to fit the LMEs as *fitlme()* is more efficient than *fitglme()* for linear data

We briefly discuss why we made changes to the steps above. Regarding step #2, we use a Bonferroni correction because we are asking if *ANY* fixed effect is significant. This is equivalent to asking independently if fixed effect #1 significant, is fixed effect #2 significant, etc.? We include interaction terms in the Bonferroni correction because the addition of interaction terms is equivalent to testing an additional hypothesis that an interaction is significant. Regarding step #5, we fit the mean response of the data with an LME then calculate the sum(*t-statistic*^2^) for the beta weights instead of calculating a cluster-mass statistic by summing the sum(*t-statistic*^2^) for the beta weights across all time samples in a cluster. Cluster-mass statistics can be biased against smaller cluster sizes whereas the sum(*t-statistic*^2^) is related to the effect size at the cluster-level and should not be biased towards cluster size.

Simulations were used to validate the use of sum(*t-statistic*^2^) even though it was not found in previous literature. We decided to use sum(*t-statistic*^2^) whole-model test statistics for several reasons. First, *t-statistic*s for the fixed effects are invariant to predictor scaling and centering. Similarly, we ignore the intercept in the calculation of the sum(*t-statistic*^2^) because intercepts are sensitive to predictor centering and traditionally dummy variables for categorical predictors are not centered. Second, the *t-statistic* from a linear model (LM) with 1 fixed effect and 2 levels (i.e. 1×2) is equivalent to the *t-statistic*s from a two sample t-test, and a two sample t-test is the most common test statistic used in CBPT. Third, we square the *t-statistic* to account for positive and negative *t-statistics* as well as to create a one-tailed permutation test statistic. Fourth, we use the sum(*t-statistic*^2^) to account for all fixed effects together. Finally, *t-statistics* are available in both LMEs and GLMEs.

### 2.2.2 Multi-step Hypothesis Testing to Control FWER

We used a multi-step hypothesis testing strategy to analyze both individual and group-level, pseudo-population data to help control for FWER across all levels of analysis. First, we identified ECoG channels whose responses significantly varied across image category and/or novelty using CBPT with LMEs. The main formula for the channel-level LME model was *’erd ∼ imageCategory + novelRepeat + (1|imageID)’ (***Table 1**). Second, we grouped channels with significant fixed effects into face-, house-, and novelty-selective pseudo-populations. Category-selective groups (i.e. face-and house-selective groups) included channels that were only category-selective as well as channels that were category-and novelty-selective while the novelty-selective group only contained channels that had novelty-selectivity; channels were not included in more than one group-level model. Additionally, we created a fourth “non-selective” group model that included all other channels that had no significant fixed effects. We analyzed the group-level data with a modified LME formula to include grouping by channel: *‘erd ∼ category + novelty + (1|channelID)’ (***Table 1**).

Typically, the final analysis step would include testing the above group-level models against alternative models. However, in this paper we simultaneously test the above group-level model against several alternative models (**Table 1**). For simplicity, alternative models can be just run on significant clusters found during initial group-level testing. However, in this paper we were interested in comparing the size and timing of significance of clusters across models so we analyzed all models simultaneously. Moreover, we strongly recommend re-running a full group-level analysis on the best model as a final step because the addition of new fixed and/or random effects may change the time course and significance of beta weights.

The overall probability of producing a type I error rate increases with the number of independent statistical tests conducted (i.e. FWER). To reduce the FWER in the multi-step hypothesis testing strategy, we divided the total α_Total_ = 0.05 in a hierarchical fashion similar to how multi-step hypothesis testing is done in clinical trials (Wang & Ting, 2014); however, we did not split α_Total_ evenly across steps. We set α_Channel_ = 0.04 for the testing of significance of individual ECoG channels because we were interested in finding more task-relevant channels rather than being overly selective. Conversely, for group-level analysis we wanted to be more confident in the results across channels so we set the group-level alpha to α_Group_ = 0.01; to account for the number of groups, we divided α_Group_ by the total number of groups (i.e. 3, for face-, house-, and novelty-selective groups). Note, we do not count testing of the “non-selective” grouped-data towards the α_Group_ threshold because we consider the testing of the “non-selective” grouped-data as verification step; namely, we want to verify that there is little to no task-related activity outside the selective channels. Importantly, significance thresholds (α) for all tests–including the significance of individual fixed effects, cluster-level permutation tests, and log-likelihood ratio (LLR) tests against alternative models–were inherited from the appropriate level in the analysis hierarchy. For example at the individual channel-level, the significance threshold in the main LME equation with 2 fixed effects was 0.02 (i.e. 0.04/2). Similarly, for the group-level analysis, the significance threshold for the LMEs with 2 fixed effects was 0.00166 (i.e. 0.01 divided by 3 groups divided by 2 fixed effects).

### 2.2.3 Testing Category-Selective Beta Weight Latency Differences

Peak latencies were detected from beta weight time courses using *findpeaks* in Matlab. Channels without detectable peaks were ignored and not included in the group-level models. Significance of latency effects at the group-level were tested using subsampling combined with permutation tests. Specifically, a new fixed effect only model was created for the group-level data using preferred vs non-preferred category instead of face vs house so that we could permute group-level labels as beta weights for face-and house-selective channels naturally had the opposite sign otherwise. We randomly permuted group-level labels and then sampled 2000 random trials (1000 to each group) to create 2 permuted group-level models. We then calculated the peak lag between the preferred vs. nonpreferred category beta weight times courses of the two permuted group-level models using cross-correlation. We repeated this procedure 1000 times to calculate a shuffled distribution of correlation lags to compare to the observed group-level lag.

## 2.3 Cluster-Based Permutation Tests Applied to Simulated Broadband Power Curves

We found it difficult to directly compare the empirical results from CBPT using LMEs to CBPT using separate t-tests because results could differ across channels or time points within a channel. Therefore, we decided to analyze simulated data generated with known properties. The main goal of this simulation was to determine how reliable separate t-tests, LMs, and LMEs perform with CBPT. Specifically, what are the differences in the probability of detecting a cluster and how does detected cluster size change across statistical models? Additionally, how does novelty effect size influence these results as novelty effect size is smaller than the category effect size? Finally, can we accurately extract the time course of fixed and random effects?

We simulated 1000 broadband power signals using statistics estimated from broadband power signals extracted from the real-world ECoG data (Table 2). The simulated curves were created by adding several gaussian smoothed curves with normally distributed values together with specific trial-conditioned responses:

**Table 2:** Simulation Broadband Power Curve Parameters.

| Variable Name | Simulation Value Used | Observed Value | Source |
| --- | --- | --- | --- |
| meanVisualResponse | 1.5 | 1.44 | Mean peak amplitude across all channels. |
| stdVisualResponse | 2.75 | 2.69 | Maximum std of ERD across all significant channels. |
| baselineTrialNoise | 0.5 | 0.51476 | Minimum std of ERD across all significant channels. |
| randomTrialNoise | 0.05 | N/A | Random white noise added to the simulation at 10% of baseline trial noise. |
| meanCategoryEffectSize | 2.0 | 1.68-2.67 | Max beta weight from house- and face-selective group-level models, respectively. |
| stdCategoryEffectSize | 1.0 | 1.55-1.81 | Std in peak beta weights across house- & face-selective channels, respectively. |
| meanNoveltyEffectSize | 0.5 | 0.27-0.63 | Max novelty beta weight from face- & house-selective group-level models, respectively. |
| stdNoveltyEffectSize | 0.25 | N/A | Hard to measure so just using half the mean. |
| imageRandomEffectSize | 0.25 | 0.25-0.56 | From alternative group-level models for face- & house-selective channels, respectively. |
| meanVisualResponseLatency | 240 | 239 | Mean peak latency across all channels. |
| meanVisualResponseHalfWidth | 100 | 85 | Mean peak half width across all channels. |
| meanCategoryLatency | 280 | 277 | Mean of peak beta weight latencies of individually category selective channels. |
| halfWidthCategory | 50-80 | 84 | Mean half width of broadband face group-level data. House group-level lacked a detectable peak so added some variation. |
| meanNoveltyLatency | 320 | 313 | From broadband house group-level data using the interaction model which had a detectable peak. |
| halfWidthNovelty | 60-100 | 71 | From house group-level data, but was difficult to estimate so added some variability. |
| baselineTransitionWidth | 10 | NA | Small smoothing transition from pre-stimulus period with no baseline effects (e.g. they'd be subtracted out) to post image onset with baseline effects. |

† Note these parameters were calculated from data obtained with an $\alpha = 0.05$ , before considering the multi-step hypothesis testing for controlling FWER. Exact parameters are unlikely to be important and unlikely to change any results or conclusions.

Response curve = visualResponse + categoryEffect + noveltyEffect + imageIdResponse + baselineNoise + randomTrialNoise.

These curves should have a single peak with a single significant cluster. For each channel, we generated simulated responses to 300 images following an ABB task design for image novelty and split the images evenly across face and house categories.

## 2.4 Analysis of High-Gamma Burst with CBPT using GLMEs

GLMEs are a nonlinear extension of LMEs and are a useful tool for analyzing data such as spike counts and other nonlinear neural phenomena. To demonstrate the utility of CBPT with GLMEs, we extracted high-gamma (70-150 Hz) bursts. Like broadband power, high-gamma power is thought to correlate with neural spiking (Manning et al., 2009), and oscillations like high-gamma oscillations are thought to be transient (Lundqvist et al., 2016). High-gamma burst events were extracted ((Gerrity, 2021); Womelsdorf lab: https://github.com/att-circ-contrl/wlBurst_v2) and stored as 0’s and 1’s representing the absence of a burst or its presence, respectively. No other burst-related data were used for this analysis.

Because gamma power is thought to reflect local processing, we used 2^nd^-order laplace to re-reference the ECoG data. At the end of our analysis, we re-analyzed the high gamma-burst data using common average referencing and found that the results were largely similar for both re-referencing types (data not shown); however, there were slightly fewer significant category-selective channels and more novelty-selective channels when we used common average re-referencing.

High-gamma burst data were analyzed in a similar process to the ERP and broadband power data. We used logistical GLMEs to model time point distributions as the presence of a burst (1) or the absence of a burst (0). At the cluster-level we modeled the high-gamma burst count using a Poisson GLME by summing burst counts across time points within each cluster. We used the canonical link functions ‘logit’ and ‘log’ for the Logistical GLMEs and Poisson GLMEs, respectively. We always measured dispersion for every instance in which we used a GLME model. Finally, we used identical model formulas for the LMEs and GLMEs. In Matlab, we used the function *fitglme()* to fit the GLMEs.

## 3.0 Results

### 3.1 Simulation Results

We conducted a large number of simulations to validate the proposed method. A full explanation of the simulation methods and results can be found in the supplementary materials, but we briefly describe the main simulation results below.

We simulated simple distributions of data to emulate the type of data found at individual time points during Step #1 in the proposed method in order to show that LMEs and GLMEs are statistically robust and produce similar results to other commonly used statistical models in ideal conditions. We found that LMEs produce highly congruent results with other linear statistical models in simple 1×2 linear simulations (**Simulation Figure 1**). However, in 2×2 linear simulations, the separate analysis of fixed effects using t-tests was underpowered compared to ANOVAs and LMEs (**Simulation Figures 2 & 3**). Furthermore, in the presence of random effects, LMEs performed better (LLR test) than models that did not include random effects such as ANOVAs and t-tests (**Simulation Figure 4**). Additionally, in 1×2 nonlinear simulations, we found GLMEs performed similarly to other nonlinear statistical models (**Simulation Figure 5**).

**Figure 1:**
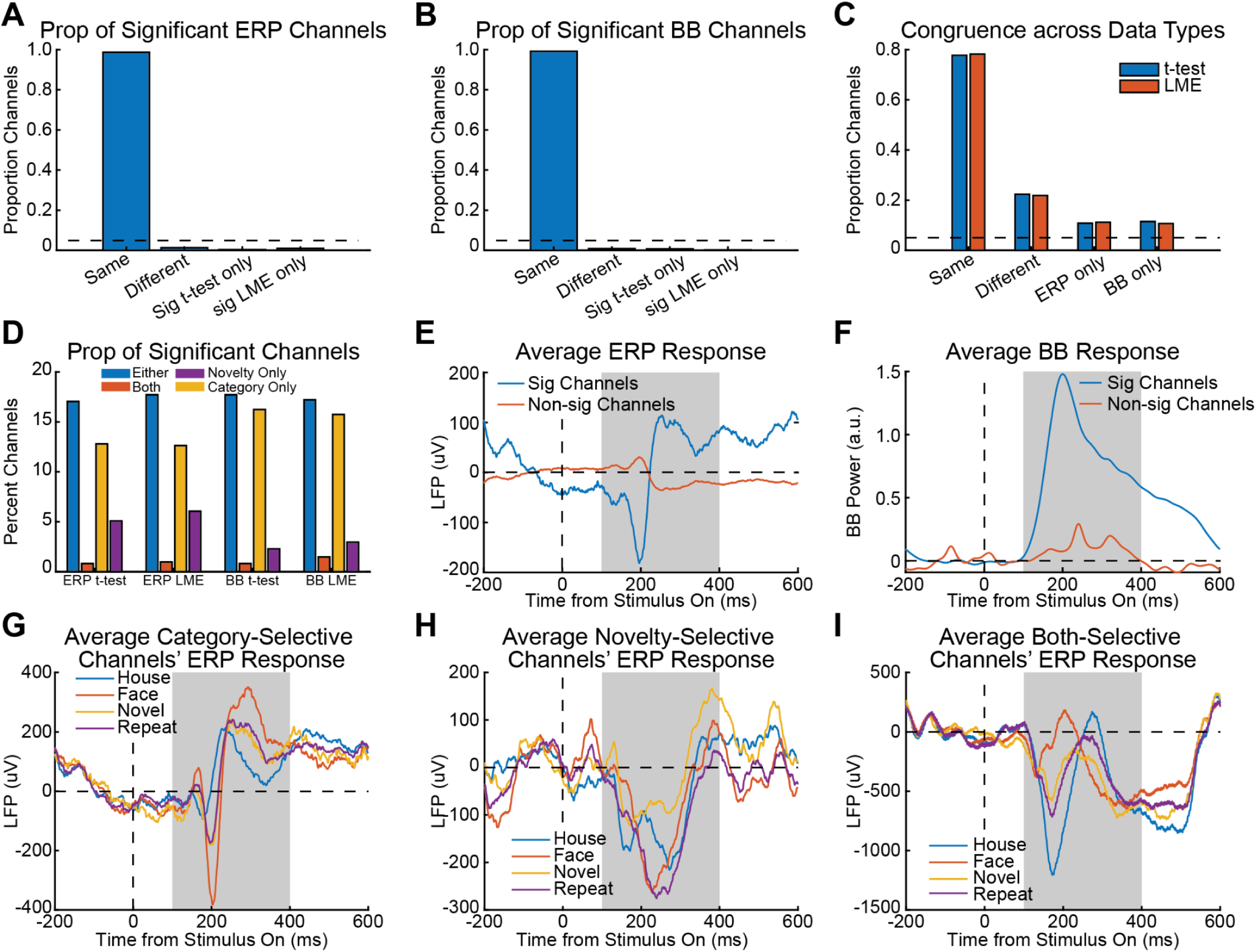
Replication of Previous Results using a Predefined Time Window. A) Proportion of channels whose ERP signals’ significance were labeled the same or different by LMEs and separate t-tests. B) Proportion of channels whose broadband (BB) power signals’ significance was labeled the same or different by LMEs and separate t-tests. C) Congruence of results across data types and statistical models. Congruence was lower across data types. D) Percent of channels labeled as significant by data type and statistical model. E) Averaged ERP responses for significant channels (blue) vs. non-significant channels (orange) showed biphasic responses as well as sustained activity after image onset. F) Averaged broadband power responses for significant channels (blue) vs. non-significant channels (orange) showed good onset timing but missed sustained activity after image offset. G) Averaged ERP responses for category-selective channels (n = 70) showed most category-selectivity occurred within the predefined time window. H) Averaged ERP responses for novelty-selective channels (n = 26) showed sustained novelty-selectivity after image offset. I) Averaged ERP responses for channels (n = 6) selective for both category and novelty showed more complicated responses with sustained activity after image offset. *Shaded regions indicate the predefined time window from 100-400 ms after image onset. Images turned off at 400 ms.

**Figure 2:**
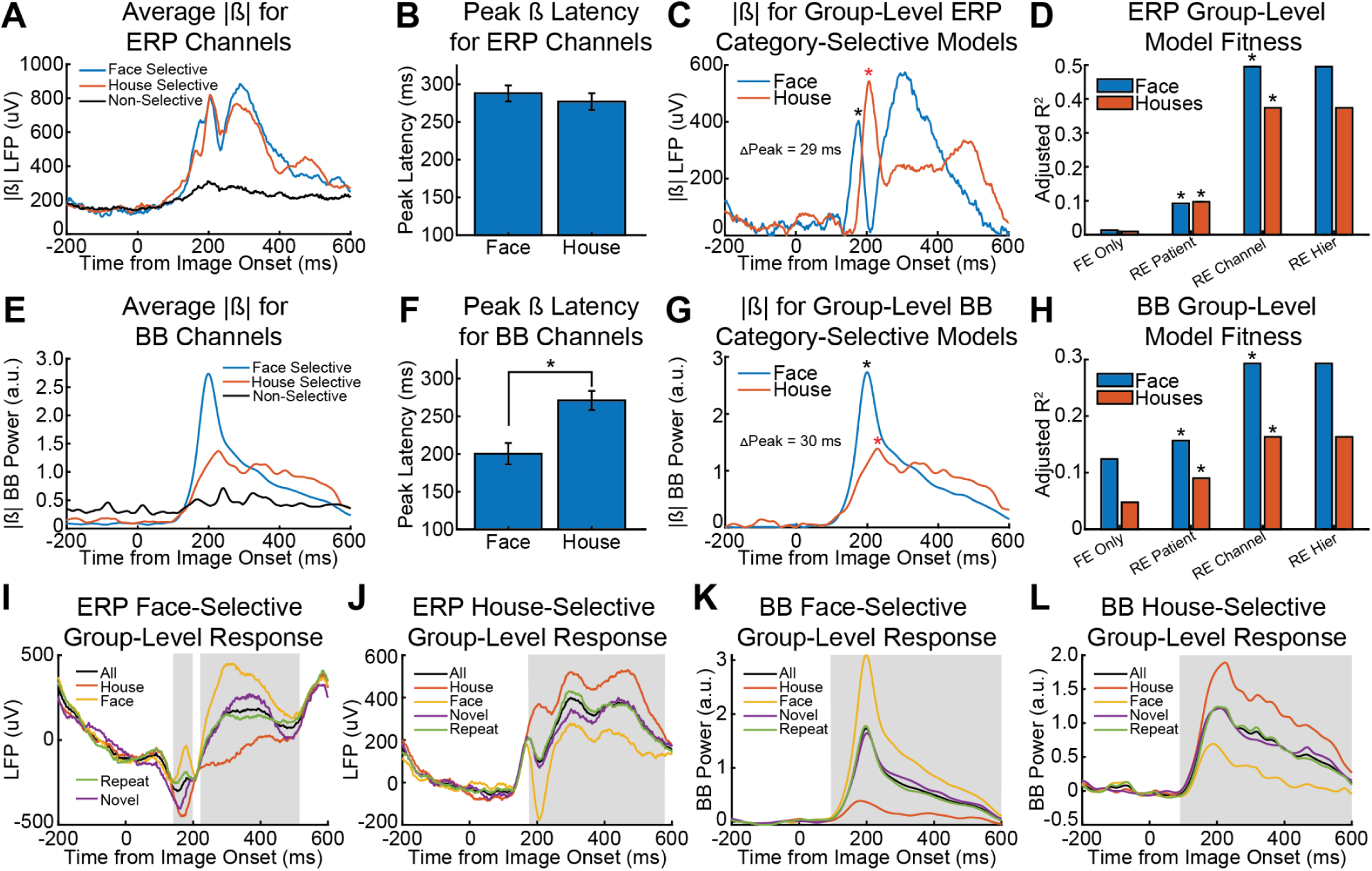
ERP and Broadband Power Analysis using CBPT with LMEs. A) Average magnitude of category beta weights for face-, house-, and non-selective ERP channels. B) Category beta weight peak latencies (mean +/-s.e.m) of face-and house-selective ERP channels (ks-test, p = 0.649). C) Magnitude of the category beta weights for face and house ERP group-level models. D) Adjusted R^2^ values for various group-level models: fixed effects-only (FE only), random effect for patient IDs (RE Patient), random effect for channel IDs (RE Channel), and hierarchical random effects (RE Hier). Asterisks indicate when a more complicated model was a better fit than the less complicated one (LLR test, p < 0.01/3). The best model for the face and house group-level ERP was RE Channel. E-H) Same as A-C except for the broadband (BB) power signals. Category beta weight peak latencies were significantly different across face-and hose-selective channels (*, ks-test, p = 8.08e-6). I-L) Average responses for face-selective ERP group-level data (H), house-selective ERP group-level data (I), face-selective broadband power group-level data (J), and house-selective broadband power group-level data (K). Shaded regions indicate significant cluster times from the RE Channel model.

**Figure 3:**
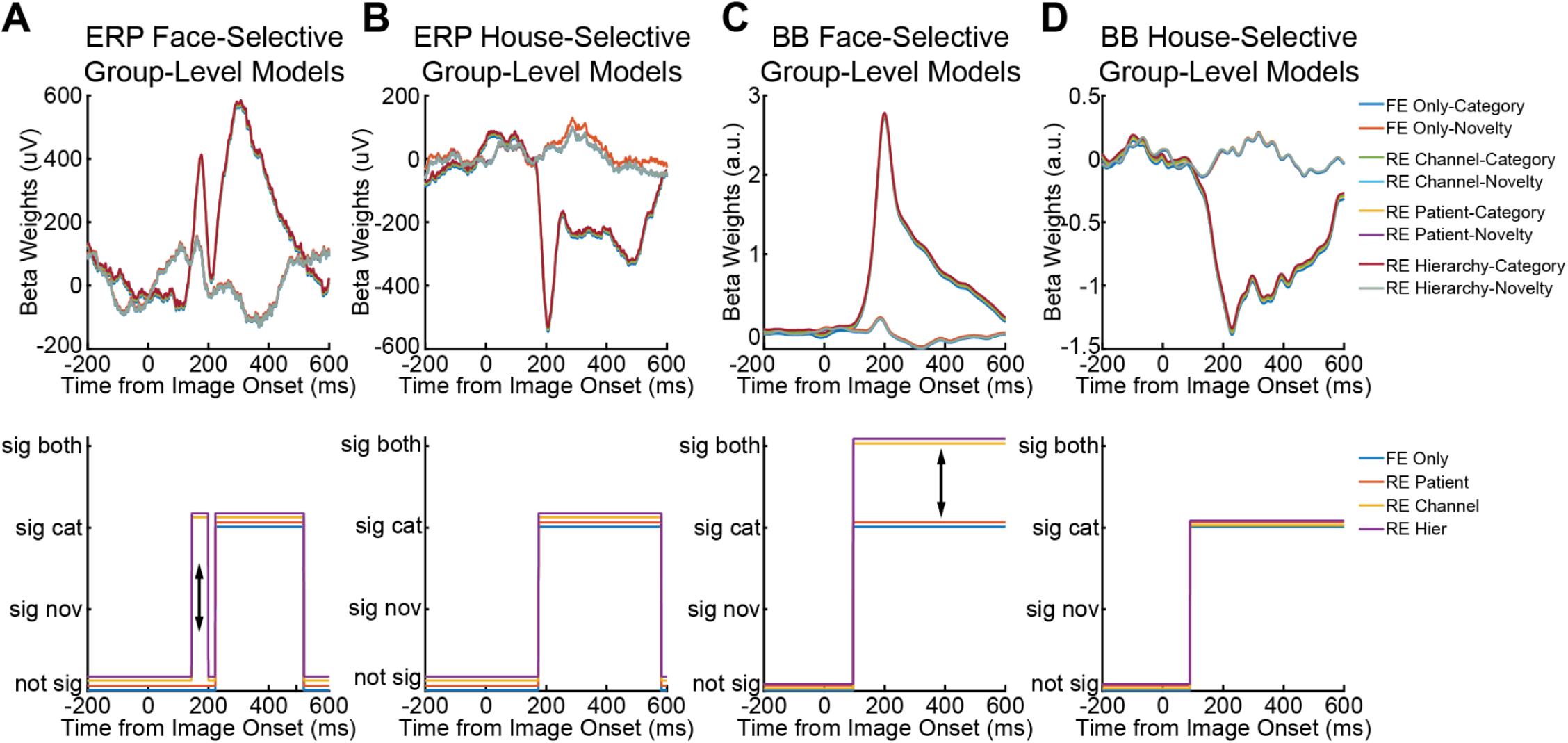
Influence of Random Effects on Beta Weights Estimates and Significance. Time Course of beta weights from various group-level models and their significance; top row is the beta weight time courses and bottom row is the associated significance time course: A) ERP face-selective group-level models, B) ERP house-selective group-level models, C) broadband power (BB) face-selective group-level models, and D) broadband power house-selective group-level models. *Note lines are offset slightly for visualization. Black arrows (↕) indicate different results between models.

**Figure 4:**
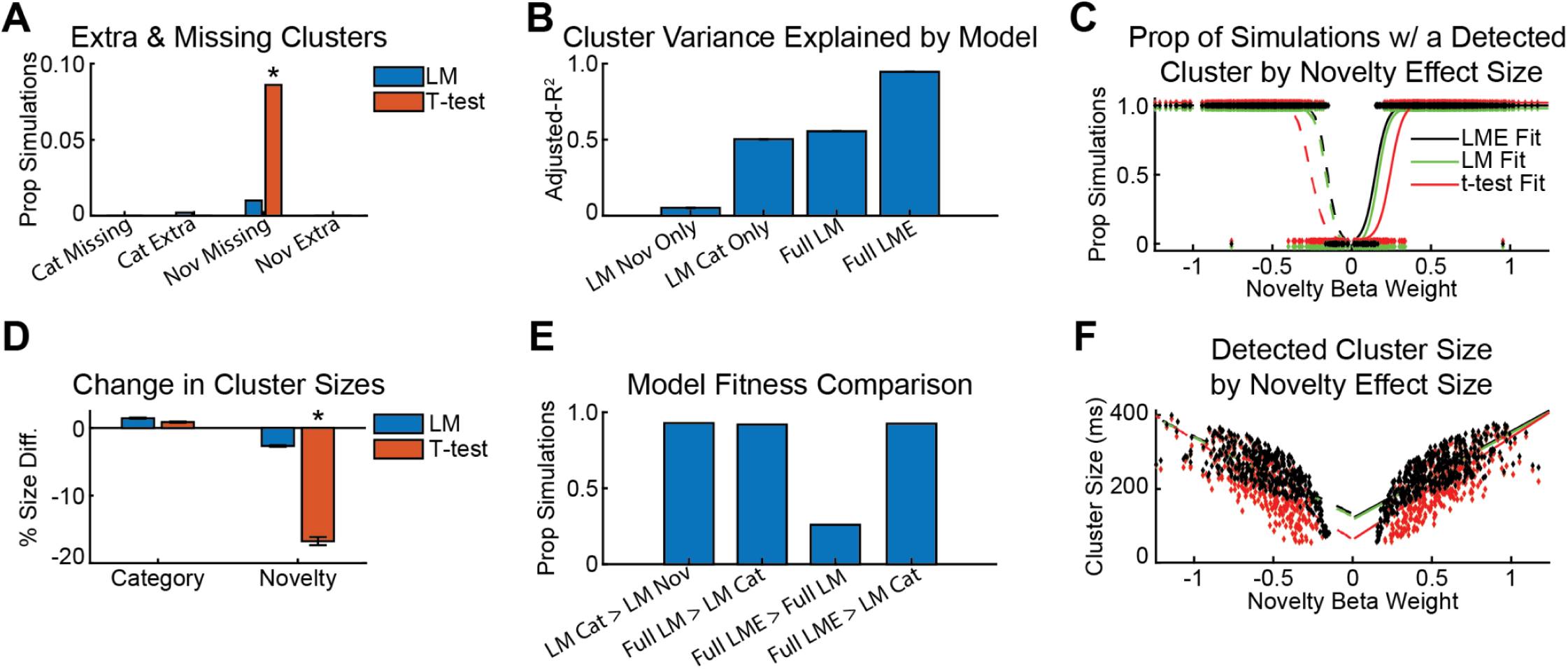
CBPT Analysis of Simulated Broadband Power Signals. A) Proportion of simulations for which LMs and separate t-tests missed or found an extra cluster compared to LMEs (*, χ2 proportion test compared to 5%, p = 0.00139). B) Adjusted-R^2^ for various partial and full LM models as well as LME models. C) The proportion of simulations with a potential cluster identified for the novelty effect as a function of novelty beta weight size. We fit this data with logistic regression models for each statistical model for positive (solid lines) and negative (dashed lines) beta weights separately. D) Percent difference in cluster sizes detected by LMs and separate t-tests compared to those detected by LMEs (*, Wilcoxon rank sum-test, p = 4.34e-16). E) The proportion of simulations that benefited from more complicated models compared to less complicated models (LLR test, p < 0.05). F) Detected novelty cluster sizes as function of novelty beta weight size for each statistical model as well as for positive (solid lines) and negative (dashed lines) beta weights.

**Figure 5:**
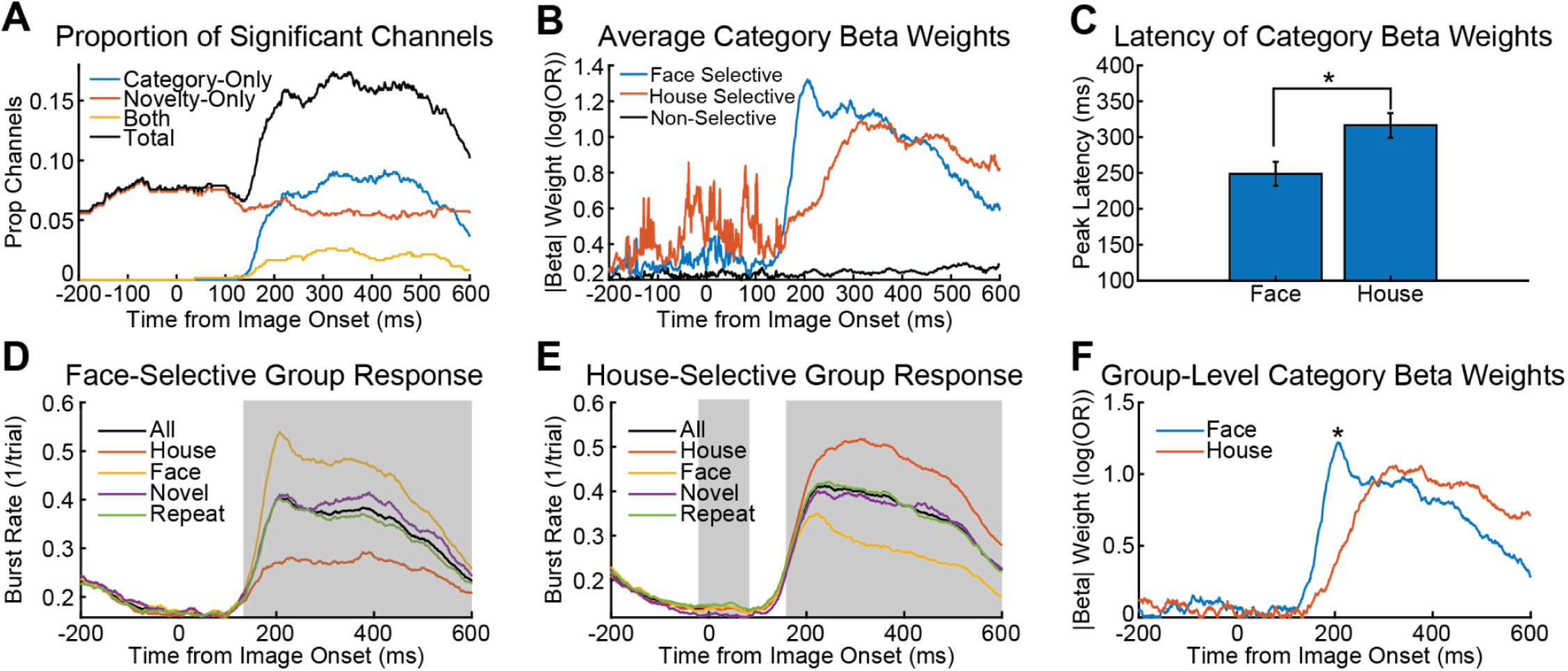
Analysis High-Gamma Burst Data using CBPT with GLMEs. A) Proportion of channels with high-gamma burst rates selective for image category, novelty, or both over time. B) Average category-selective beta weight magnitudes for individual face-, house-selective, and non-selective channels. C) Peak latency (mean +/-s.e.m) of category beta weights for face-and house-selective channels (*, ks-test, p = 0.00852). D) Average burst rate for face-selective group-level data. Shaded regions indicate significant time points after CBPT; the cluster was significant for both category and novelty. E) Average burst rate for house-selective group-level data. Shaded regions indicate significant time points after CBPT; the first cluster was significant for novelty and the second cluster was significant for image category. F) Category beta weight time course for face-and house-selective group-level data. A peak (*) was detected for the face-selective group-level data but not the house-selective group-level data.

Next, we used simulations to validate the other aspects of our proposed method. We found, regardless of the permutation method, in 2×2 linear simulations that sum(*t-statistic*^2^) was a good whole-model test statistic while R^2^ was unreliable because it was prone to the influence of random effects (**Simulation Figure 6**). Furthermore, in 1×2 simulations we found that results from using sum(*t-statistic*^2^) as a cluster-level statistic were consistent with results from using traditional cluster-mass statistics from t-tests when analyzing linear data and cluster-mass statistics from *χ^2^*-test when analyzing nonlinear data (**Simulation Figure 7**). Finally, in simple linear 4x(1×2) simulations of multiple independent fixed effects, we found that not properly controlling for the number of fixed effects increased the FWER (**Simulation Figure 8**). Importantly, we found that using *t-statistic* with a Bonferroni correction for the number of fixed effects was only slightly more conservative than a Bonferroni-Holm correction while also being very easy and straightforward to implement.

**Figure 6:**
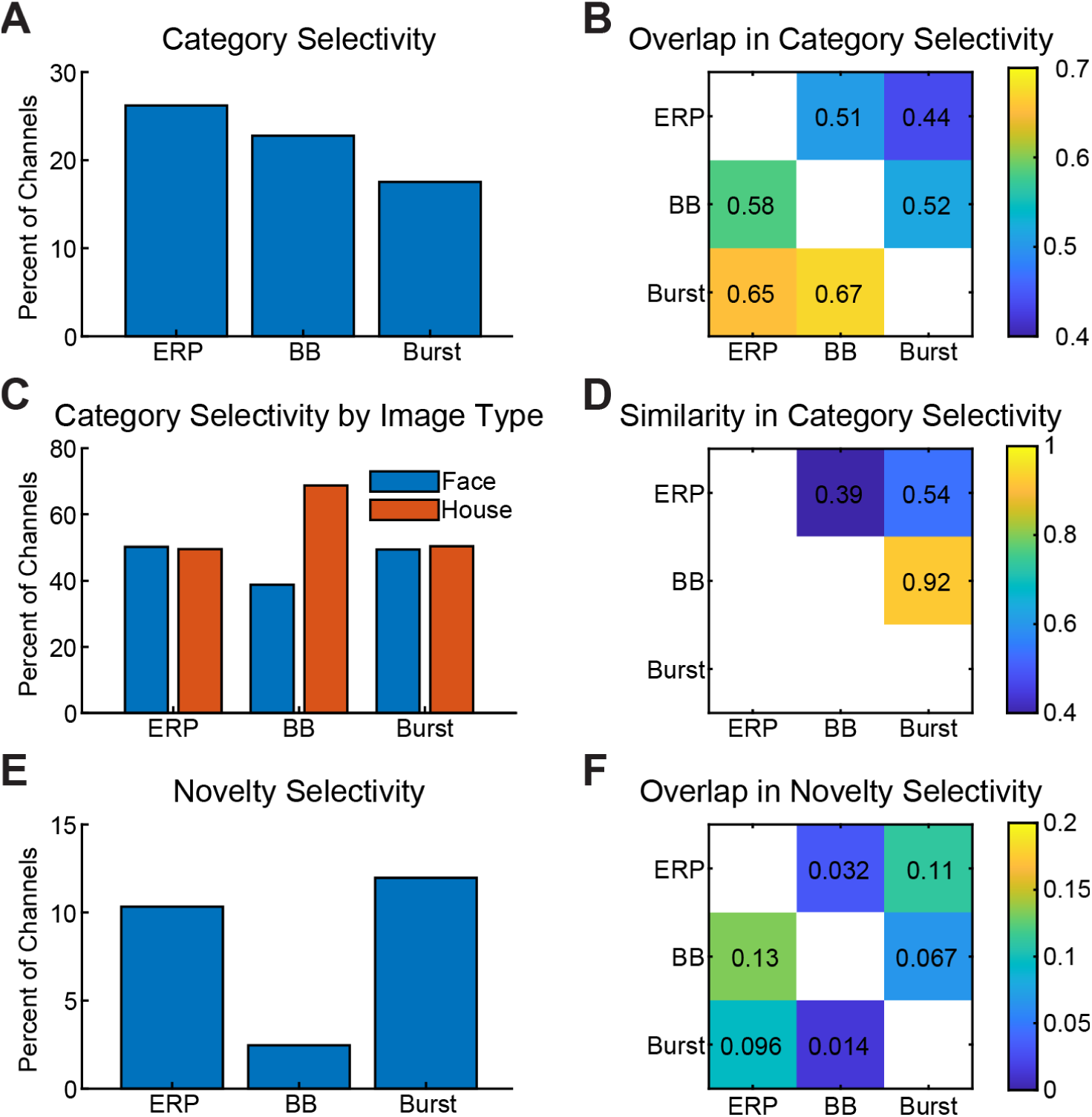
CBPT Summary Results. A) Percent of channels with significant category-selectivity across different neural signals. B) Comparison of channels with category-selectivity across different neural signals. Values indicate the proportion of channels that have category selectivity for both data types; plot is asymmetric. For example, there was only a 0.44 overlap for ERP and [high-gamma] Burst (1,3) signals; this means for all the channels whose ERP signals were category selective, only 44% of those channels had high-gamma burst signals that were also selective for category. Conversely, for Burst vs. ERP (3,1), for all the channels that had high-gamma burst signal selective for category, 65% of channels also had ERP signals selective for category. C) Percent of category-selective channels selective for face or house images by neural signal type. D) Comparison of category-selective channels by selectivity type (i.e. face vs house) across neural signals. For example Burst vs BB (2,3), 92% of category-selective channels had the same selectivity. E) Percent of channels with novelty-selectivity across neural signal types. F) Comparison of channels with novelty-selectivity across neural signals; plot is also asymmetric like B.

Overall, these simulation results show that the proposed method using CBPT with LMEs and GLMEs controls well for type I error, type II error, and FWER.

### 3.2 Replication of Predefined Time Window Analysis Results

To replicate previously published results, we applied separate t-tests and LMEs to ERP and broadband power signals in a predefined time window (K. J. Miller et al., 2015, 2016). Specifically, we analyzed ERP and broadband power signals between 100 ms and 400 ms after image onset extracted from 610 ECoG channels from 12 subjects. Results from separate t-tests were highly congruent with results from LMEs for ERPs (98.7%) and broadband power (99.2%) signals (**Figure 1A and 1B**), but results were less congruent across data types (78%, **Figure 1C**). Further, the percent of channels significant for category, novelty, or both was similar to previous publications (K. J. Miller et al., 2015, 2016) with 12.6% to 16.2% of channels being significant for category only (**Figure 1D**). Specifically, separate t-tests and LMEs found that 12.7% (78/610) and 12.6% (77/610) of ERP channels were category selective, respectively. Separate t-tests and LMEs found that 16.2% (99/610) and 15.7% (96%) of broadband power channels were category selective, respectively. Novelty selectivity was comparatively limited with LMEs finding slightly more novelty selective channels: t-test ERPs 5.1% (31/610), LMEs ERPs 6.1% (37/610), t-tests broadband power 2.3% (14/610), and LMEs broadband power 3.0% (18/610). Selectivity for both category and novelty was extremely rare with only 5 to 9 (0.8% to 1.4%) channels showing selectivity for both across data types and statistical methods.

Visual inspection of the category and novelty selective channels’ signals revealed activity that was outside of the predefined time window (**Figure 1E/F**). Namely activity appeared to be sustained for longer than the 400 ms image display period. Furthermore, averaged activity from ERP-selective channels appeared to have both positive and negative response periods. Individual channels showed biphasic responses as well (data not shown). More detailed visual inspection of averaged ERP activity from category only (**Figure 1G**), novelty only (**Figure 1H**), and both category and novelty (**Figure 1I**) selective channels indicated that category selectivity was largely limited to the first few hundred milliseconds after image onset while novelty selectivity was sustained for longer periods of time. A similar pattern was observed for the broadband power data (data now shown). Overall, these temporal patterns were not well captured by using a predefined time window.

We also tested several alternative LME models on the significantly selective channels. For ERP selective channels, only 27.5% channels benefited from the inclusion of random effects (LLR test, p < 0.05, **Supplementary Figure 1 A**), and 44.2% of broadband power selective channels benefited from the inclusion of random effects (LLR test, p < 0.05, **Supplementary Figure 1 D**). Very few (8.8% ERP, 12.5% broadband) channels benefited from an interaction between image novelty and category (**Supplementary Figure 1 B/E**), but some broadband channels with selectivity for both novelty and category appear to benefit from an interaction term (6/9, 66.6%). We also tested whether the number of times an image was shown was a better fit than a binary novelty-repeat effect. For both ERP and broadband selective-channels, 44.5% of channels benefited (LLR test, p < 0.05) from a model with the number of times an image was shown. However, it was not possible to interpret this result further because the number of times an image was shown was correlated 1:1 with block number.

In summary, we replicated previously published results. Category selectivity for both faces and houses was common in the ERP and broadband power signals (12.6% to 16.2%), while novelty selectivity and selectivity for both category and novelty was comparatively rare (0.8% to 6.1%). Importantly, visual inspection of the time course of selective channels was not always well captured by the predefined time window from 100 to 400 ms after image onset. Specifically, the averaged ERP data showed biphasic response. Additionally, averaged ERP and broadband power data showed sustained responses after image offset especially for image novelty. Together, these data suggest that a time window defined *a priori* may be inferior to empirically defined time windows. Lastly, t-tests and LMEs produced highly congruent results (≥ 98.7%) within data type, while results were less congruent across data types (78%).

### 3.3 ERP & Broadband Power Analysis using CBPT with LMEs

Next, we reanalyzed the ERP and broadband power signals using CBPT with LMEs. We used α_Individual_ = 0.04 and α_Group_ = 0.01/3 for significance thresholds for individual channels and group-level model analyses, respectively. Additionally, we used a minimum cluster size (*t*_min_) of 55 ms determined from the 98%-tile of the baseline cluster sizes from the ERP data using two separate t-tests (Cohen, 2014).

We had a difficult time comparing results across CBPT with t-tests and CBPT with LMEs due to differences in results across channels and time points within a channel (**Supplementary Figure 2**). Nonetheless, at the channel-level, t-tests and LMEs were reasonably congruent: 87.5% for the ERP data and 95.1% for the broadband power data. Modeling results in the next section (3.4) were used to fully explain this reduction in congruence, but for the remainder of this section we solely focus on results from using CBPT with LMEs.

Analysis of the ERP signals using CBPT with LMEs found 126 (20.6%) channels selective for category, 63 (10.3%) channels selective for novelty, and 52 (8.5%) channels selective for both category and novelty. Analysis of the broadband power signals using CBPT with LMEs found 129 (21.1%) channels selective for category, 15 (2.5%) channels selective for novelty, and 14 (2.3%) channels selective for both category and novelty (**Supplementary Figure 3**). Overall, the number of significant channels identified by CBPT with LMEs was higher than what was found by the predefined time window analysis.

One major advantage of using CBPT with LMEs is that we incidentally extract the time course of the beta weights (**Figure 2A/E**). We focused on the category selective channels since there were too few broadband power channels with novelty selectivity. We determined which channels were face-or house-selective based on the sign of the category beta weights within the largest significant cluster. Visual inspection of the beta weights from the ERP signals showed no obvious difference in averaged response for face-(n = 74) and house-selective (n = 73) channels nor was there a significant difference in their peak latency (ks-test, p = 0.649, **Figure 2B)**. However, visual inspection of the category-selective beta weights from the broadband power data suggested that the face-selective channels’ selectivity (n = 40) occurred prior to the house-selective channels’ selectivity (n = 71). Additionally, beta weight peak latencies for face-selective channels occurred earlier than the beta weight peak latencies for the house-selective channels (*ks-test*, p = 8.08e-6, median difference = 70.5 ms, **Figure 2F**).

We next built group-level models of the face-and house-selective channels in order to analyze functionally similar channels as pseudo-populations. This group-level analysis was designed to identify consistent patterns of activity across channels including activity that may not have been clearly present in individual channels’ activity. While there was no latency pattern observed in the individual ERP channels, the group-level face-selective ERP beta weights had an initial peak 29 ms earlier (177 ms vs 206 ms) than the group-level house-selective beta weights (**Figure 2C**). The time course of the face-and house-selective broadband power data appeared similar to the average of the individual channels; the beta weight peak latency of the group-level face-selective data occurred 30 ms before (199 ms vs 229 ms) the group-level, house-selective data (**Figure 2D**). We used cross-correlation with subsampling and permutation tests to determine if these observed group-level latency differences were significant. We found a significant difference (-27 ms, 99.9 percentile) in the latency of the broadband power category-selective group-level data, but there was no significant latency effect (-37 ms, 82.9 percentile) in the group-level ERP data.

Next, we asked how random effects influence the results (**Figure 2D/H**). To this end we fit the ERP and broadband power group-level data with 4 different random effects models (see Table 1). First, all models that included random effects fit the data better than the fixed effects-only models (LLR test, all p’s < 0.01/3). The best models included channel ID as random effect (LLR test, all p’s < 0.01/3); there was no additional benefit of a hierarchical random effects model with channel ID nested within patient ID, but the hierarchical random effects models were not worse than the random effects models with channel ID. Second, we found that while the [mean] beta weight estimates were minimally impacted by the addition of random effects, the significance of fixed effects were impacted by the inclusion of various random effects (**Figure 3**). For example in the face-selective group-level ERP data, the fixed effect-only model did not identify a significant cluster around 200 ms while the random effects model with channel ID did. In another example, the face-selective group-level broadband power data, the fixed effect-only found that the main cluster was selective for category only, but the random effects model with channel ID found that the same cluster was selective for both category and novelty; this significance change was a result of a slight (12.75%) change in the mean beta weight estimate (-0.05965 vs -0.067255) and a slight change (10.0%) in the standard error (0.023588 vs 0.02122) which overall led to a more substantial change (25.3%) in the t-statistic (-2.5288 vs -3.1695) associated with the novelty effect. Overall, these results suggest that subpar models–like the fixed effect-only model–may lead to different interpretations of the data compared to models that better reflect the data structure–like the random effects model with channel ID.

Next, we tested alternative models with interaction terms against the random effects model with channel ID. For the ERP face-selective group model, we only fit the second, larger cluster occurring after 200 ms. All category-selective group-level data were fit better by some sort of interaction model except for the ERP house-selective group-level data (**Supplementary Figure 4**) indicating at the pseudo-population level there is generally some interaction between image category and novelty.

Next, we compared the mean responses taken from the predefined time window to the mean responses in the largest clusters’ time window using the interaction model with the imaged ID. For the ERP group-level data, we found the adjusted-R^2^‘s for the LMEs applied to the cluster-defined windows were higher than the adjusted R^2^‘s for the LMEs applied to the predefined windows (**Supplementary Figure 4**). For the broadband power group-level data, the adjusted-R^2^‘s for the LMEs applied to the cluster-defined windows were slightly lower than they were for the predefined window. However, a LLR test showed that the cluster-defined windows were all a better “fit” for the ERP and broadband power group-level data than the predefined windows (all p’s < 0.01/3), suggesting that CBPT identified a better time window than the *a priori* selected window.

Finally, we briefly analyzed the group-level novelty-selective and non-selective data. Neither the novelty-selective ERP nor the novelty-selective broadband group-level data had any significant clusters (**Supplementary Figure 5**). For the ERP data, there appeared to be a significant difference at some time points, but there were no potential clusters longer than 21 ms. For the broadband data, 15 channels may simply not have been enough to show a significant effect. We then analyzed the non-selective channels to ensure that we did not miss anything important in the data (**Supplementary Figure 6**). For the ERP data (n = 400) there were no significant clusters identified, but again there appeared to be a short lasting category effect. For the broadband power data (n = 460), we found two small clusters, but the data appeared quite messy and the models explained very little variance in the data as expected. Interestingly, the hierarchical random effects model fit these clusters’ data better than any other model (LLR test, p’s < 0.01/3).

### 3.4 CBPT with LMEs Applied to Simulated Broadband Power Signals

We had difficulty comparing results from CBPT using separate t-tests to results from CBPT using LMEs in the previous section because there were differences at the channel-level (i.e. whether a channel had a significant cluster) and at individual time points (i.e. different cluster sizes and/or locations). Because our simple 2×2 simulations showed separate t-tests were less powerful than models that consider all fixed effects together (**Simulation Figures 2 & 3**), we hypothesized that CBPT with separate t-tests would identify fewer significant clusters and these clusters would be smaller especially for the smaller fixed effect sizes here associated with image novelty. We also wanted to determine how well the extracted fixed and random effects time courses reflected the true fixed effect and random effect time courses. To this end we simulated 1000 broadband power signals (see Table 2) and extracted potential cluster times as well as fixed and random effect time courses; potential clusters were assumed to be significant but we did not run permutation tests to verify.

LMs and LMEs were able to accurately extract the expected beta weight time courses with an average error rate < 0.1%. However, it was possible to generate signals where the statistical properties–especially with unbalanced random effects–were such that the observable statistical differences were substantially different from the true underlying distributions from which the data was generated (data not shown). In the results below, the beta weight time courses were reasonably well extracted (< 10% error) for all simulated channels (**Supplementary Figure 7B**). Random effects time courses were also not always accurately extracted either (**Supplementary Figure 7D**); the extracted random effects reflect the observable statistical differences in the data and not necessarily the true underlying distributions from which the data was generated, and the variance we added was sometimes attributed to different random effects. Note, these random effects results could be an artifact of how we generated the data rather than an extraction error per se.

CBPT using LMs performed very similarly to CBPT using LMEs while CBPT using separate t-tests were underpowered especially for novelty clusters. At the channel-level, LMs and LMEs were 99.8% congruent for category clusters and 99.0% congruent for novelty clusters. CBPT using LMEs and CBPT using separate t-tests were 100.0% congruent for the category clusters but only 91.4% congruent for novelty clusters. CBPT using LMs identified 0.2% extra category clusters not identified by CBPT using LMEs; no extra category clusters were identified by the CBPT using separate t-tests (**Figure 4A**). CBPT using LMs missed 1.0% of novelty clusters that CBPT using LMEs found, and the separate t-tests missed 8.6% (χ^2^ proportion test vs 5.0%, p = 0.00139) of novelty clusters that the CBPT using LMEs found. Furthermore, the novelty clusters identified by the CBPT using separate t-tests were significantly smaller than the novelty clusters identified by CBPT using LMEs (Wilcoxon rank sum test, p = 4.34e-16, mean differences = -16.7%), but category clusters were not significantly different in size (Wilcoxon, rank sum test, p = 0.4479, mean difference = 0.86%). There was no statistical difference (Wilcoxon rank sum test, p’s > 0.14) for the cluster sizes for CBPT using LMs compared to CBPT using LMEs for category (mean differences = 1.43%) or novelty (mean difference = -2.67%) clusters (**Figure 4D**).

Next, we analyzed how well LMs and LMEs fit the clusters’ data. As expected, we found adjusted-R^2^ increased with model complexity (**Figure 4B**). We then compared LM and LME models fitness using a LLR test to determine if LMEs were a better fit for the clusters’ data (**Figure 4E**). LMEs fit the clusters’ data better than LMs in only 26.0% of simulated channels for which there was an overlapping cluster. We also ran reduced models of the LM compared to the full LM and the full LME. LMs with category fit the clusters’ data better than LMs with novelty-only in 92.8% of simulations and full LMs fit the clusters’ data better than LMs with category-only in 91.9% of simulations. Furthermore, LMEs fit the clusters’ data better than LMs with category-only in 92.4% of simulations.

Finally, we asked if the probability of detecting a cluster or the cluster size was related to novelty effect size. We fit the probability of finding a cluster with a logistic regression model (**Figure 4C**), and we fit the detected cluster size with a linear regression model (**Figure 4F**). There was not a significant relationship between the probability of finding a cluster and novelty effect size (p’s > 0.51), but there was an apparent shift in the logistic regression curves for the separate t-tests; likely there were not enough simulations (46-90/1000) without a detected clusters to do a good comparison. However, there was a significant relationship between novelty effect size and cluster size for all models (beta weights, p’s < 1.06e-13). LMs and LMEs had statistically similar (equivalence test, overlapping CI) slopes for both positive and negative novelty effects. However, the slopes for separate t-tests and LMEs were statistically different (equivalence test, non-overlapping CI) for both positive and negative novelty effects.

Overall, these results suggest CBPT usig LMEs performs at least as well or better than CBPT using LMs, but CBPT using separate t-tests was significantly underpowered. This decrease in power led to fewer detected clusters and detected clusters that were smaller for the novelty effect. Additionally, in some but not all simulations there was a benefit of adding random effects to a linear model. The random effects in these simulations were perfectly balanced across groups and generated from perfect, normal distributions; the benefit of random effects should increase as conditions become less normal, random effect amplitudes increase, and as random effects become less balanced as seen in previous simulations. Moreover, models that included both fixed effects (i.e. category and novelty) fit the data almost always better than models that only considered the biggest effect only (i.e. category).

### 3.5 Analysis of High-Gamma Burst using CBPT with GLMEs

We analyzed high-gamma burst rates to demonstrate the flexibility of CBPT using GLMEs for nonlinear data. We extracted high-gamma (70-150 Hz) bursts and aligned bursts to image onset similar to how we analyzed the ERP and broadband signals (**Supplementary Figure 8**). The presence of a burst was represented by a 1 and the absence of a burst was represented by a 0. We then modeled the high-gamma burst data using Logistical GLMEs at individual time points and Poisson GLMEs at the cluster-level. Correspondingly, beta weights at the individual time points represent the natural log-Odds Ratio (log(OR)) of burst rate (1/trial) while the beta weights at the cluster-level represent natural-log-Rate Ratios of burst counts.

We found 107 (17.5%) channels had high-gamma burst signals selective for category or category and novelty as well as 73 (12.0%) channels that had high-gamma bursts selective for novelty only. Novelty-selectivity occurred throughout the analysis window while category selectivity was found only after image onset (**Figure 5A/B**). Analysis of high-gamma burst category-selective channels found that the category beta weights for face-selective (n = 51) channels peaked prior to the house-selective (n = 52) channels (ks-test, p = 0.00852, 249 ms vs 316.5 ms, **Figure 5C**).

Next we built group-level models for the face-, house-, novelty-, and non-selective (n = 447) channels. The face-selective group-level data had one significant cluster selective for both category and novelty after image onset (**Figure 5D**). The house-selective group-level data had 2 significant clusters, one around image onset selective for novelty and one after image onset selective for category (**Figure 5E**). The extracted beta weight time course from the group-level category-selective models suggested that the category-selective activity of the face-selective channels preceded that of the house-selective channels, but there was no discernable peak in category beta weights for the house-selective group-level data (**Figure 5F**). Cross-correlation with subsampling and permutation tests indicated that category-selectivity for the face-selective group-level data significantly preceded category-selectivity for the house-selective group-level data (-57 ms, 99.8 percentile).

We again tested the influence of random effects on model fitness. The random effects models generally outperformed the fixed effects-only model except patient ID for the house-selective group, and the best models for the face-and house-selective group-level models included the channel ID as a random effect. Different random effects models did not influence the significance of the clusters or their timing (data not shown).

Analysis of the novelty-selective group-level data found that almost all time points were significant for image novelty and that the post-image onset period was also weakly selective for image category **(Supplementary Figure 9**). There was also evidence that different random effects models influenced the beta weights and their significance; surprisingly the best model included patient ID as a random effect. Two apparently spurious significant clusters were identified in the non-selective group-level data (**Supplementary Figure 10**), but the beta weights were very small.

### 3.6 Summary Results

We briefly compared the results from the analysis of different neural signals. ERP signals had the largest proportion of category-selectivity channels followed by the broadband power signals, and lastly the high-gamma burst signals (**Figure 6A**). On the other hand, high-gamma burst signals had the largest proportion of novelty-selective channels followed by the ERP signals while the broadband power signals had the fewest (**Figure 6E**). Note we removed the baseline activity levels for the ERP and broadband signals which likely affected the proportion of channels with novelty-selectivity especially for the broadband power signals.

We asked if a channel’s selectivity was similar across different neural signals? Category-selective channels were only modestly similar across different neural signals (**Figure 6B**). ERP category-selectivity overlapped the least with other signals (44% to 51%). However, channels with high-gamma burst signals that were category-selective had a higher probability of having ERP signals (65%) and broadband power signals (67%) that were also category-selective. Moreover, the type of category-selectivity (i.e. face-vs. house-selective) was very similar (92%) for broadband power signals and high-gamma burst signals (**Figure 6D**), but this pattern was not true of the ERP signals (39% to 54%). Novelty-selectivity was very dissimilar across different neural signals (**Figure 6E**).

We also looked at the distribution of recording locations for each type of neural signal and their category-selectivity. While areas like the fusiform gyrus and parahippocampal had greater category-selectivity representing their canonical functions, there was widespread category selectivity across many brain areas making it somewhat difficult to interpret brain area specific results (**Supplementary Figure 11)**. Additionally, results varied slightly depending on the analysis method and neural data type analyzed.

## 4.0 Discussion

### 4.1 General Discussion

We demonstrated the feasibility, flexibility, and robustness of using CBPT with mixed effects models to analyze experiments with multiple fixed effects. We applied CBPT with LMEs and GLMEs to three different types of signals–ERPs, broadband power, and high-gamma bursts–representing a wide range of signal types found in neuroscience. We also combined CBPT with a multi-step hypothesis testing strategy to analyze individual channels and group-level, pseudo-population data to control for the total FWER by splitting α_Total_ into α_Channel_ and α_Group_. We also used the group-level models to identify differences in functional groups of data.

Our results provide strong evidence that the use of appropriate statistical models is necessary to conduct both sensitive and reliable time-series data analysis. We showed through simulations that LMEs and GLMEs perform similarly to appropriately applied linear and nonlinear statistical models in ideal scenarios, but the statistical power of many common statistical models was reduced in the presence of random effects. Furthermore, failure to account for all fixed effects in a single statistical model–for example by running separate t-tests–further reduced statistical power. This type of statistical power loss was compounded in the CBPT analysis of simulated broadband power signals: CBPT’s sensitivity for detecting clusters was reduced and the resulting cluster sizes were smaller. Similarly, the inclusion of appropriate random effects (e.g. channel ID) in the group-level models changed the significance of beta weights compared to fixed effects-only models. Additionally, fixed effects-only models sometimes missed clusters entirely.

Combining these empirical and simulation results together, we show that our approach uses appropriate statistical models necessary to maintain control of type I error, type II error, and FWER. Failure to do so can lead to inexact results and inappropriate conclusions, which needlessly adds confusion to a field already troubled with poor reproducibility and repeated translation failures (Open Source Collaboration, 2015). Thus, providing better tools for robust statistical analysis is of the highest priority.

### 4.2 Simulation Results

We ran a large number of simulations to validate our proposed method. For each step in our proposed method, we ran simulations comparing mixed effects models to other commonly used statistical models. Across a large range of simulation parameters, for both linear and nonlinear data, we found that results from mixed effects models were highly congruent with other commonly used statistical models. Importantly mixed effects models never identified significant results where other appropriate statistical models could not. However, in the presence of random effects, even in simple simulations with a single fixed effect, LMEs outperformed fixed effects-only models. The reliability of LME and GLME results over a wide parameter space is extremely important as fixed and random effect sizes can change drastically over time within the analysis window, necessitating a statistical framework that is both flexible and robust. Furthermore, since random effects are ubiquitous in most experimental settings, this result adds to a growing literature on the importance of building statistical models that incorporate random effects (Yu et al., 2022). We extended these results to the analysis of clusters of data, and found that fitting the mean of a cluster’s response with a new LME produced essentially equivalent results to a cluster-mass statistic based on t-tests. Similarly, we found that fitting the sum of a cluster’s response with a new Poisson GLME produced equivalent results to a cluster-mass statistic based on χ^2^ tests. The main advantage of fitting the cluster-level response with a new mixed effects model is that we ultimately want a model of the aggregate response and therefore can combine two steps into one.

Our results have important implications for the analysis of experimental designs with multiple fixed effects. LMEs and ANOVAs–analyzing fixed effects together in a single model–outperformed separately run t-tests, which, dismayingly, remain commonly used despite widespread familiarity with ANOVAs and other linear models. We extended these results to broadband power simulations, showing that separately running t-tests was significantly underpowered compared to LM[E]s in which we could still accurately extract the time course of fixed and random effects.

Given the clear superiority of complete models, we ran additional simulations to determine a good whole-model test statistic. We found that sum(*t-statistic*^2^) was a good whole-model test statistic, while R^2^ generally performed poorly in the presence of random effects producing many false positives. These results are not surprising as R^2^ ***is not*** a measure of model fitness per se but rather measures dispersion of the data around the mean model fit. We also assessed multiple permutation methods for analyzing whole-model results, and found that full-model permutation methods (Manly), full-model residual permutation methods (ter Braak), and reduced model permutation methods (Freedman-Lane-style) all produced similar outcomes.

### 4.3 Empirical Results

Turning to real-world data, we re-analyzed an existing dataset of human ECoG responses to visual stimuli with our statistical framework (K. J. Miller et al., 2015, 2016). Our predefined time window analysis of ERPs and broadband power signals produced similar results to previous findings (K. J. Miller et al., 2015, 2016). However, many of the significantly selective channels showed responses that were outside of the predefined time window suggesting that CBPT would be better for analyzing these data. Additionally, ERPs showed biphasic responses within the predefined window suggesting averaging some signals across the window could cancel effects. Not surprisingly, analysis of ERPs and broadband power signals using CBPT with LMEs found more channels selective for image category and novelty compared to the predefined window analysis. Additionally, results from CBPT using separate t-tests and results from CBPT using LMEs differed more, even though results were similar in the predefined time window analysis. In the LME analysis, we found more novelty-selectivity with the CBPT analysis, underscoring the sensitivity gained by using complete models. Furthermore, if we had only looked for a novelty effect within a predefined time window, then we would have missed that novelty was represented during the majority of the trial period; many experiments have contextual cues, like novelty, that are persistent over the trial period, and thus these important contextual effects could be missed when using predefined time windows.

Whereas group-level inferences are often made indirectly based on single channel results, we leveraged the power of our mixed effects model framework to directly compute group-level effects across pseudo-populations of channels. For instance, we built group-level models for the significant broadband power and ERP channels. Analyzing the group-level beta weights, we observed that broadband power category-selective signals emerged earlier when viewing faces than houses, a pattern consistent with the individual channel responses. We observed a smaller, non-significant effect in the ERP group-level models as well. Furthermore, we found evidence of an interaction effect between category and novelty in ¾ group-level ERP and broadband power models despite rarely finding interaction effects in individual channels. Overall, these data suggest that extrapolating results from the analysis of individual channels may not generalize well to group-level results in all scenarios.

We also explored the impact of various random effects on model performance. Consistent with our simulations, models that included random effects fit the group-level data better than the fixed effects-only models, and models that included channel ID as a random effect fit the broadband power, ERP, and high-gamma group-level category-selective models best. We did not see any benefit in including hierarchical random effects in which channel ID was nested within patient ID. However, there may be a benefit of hierarchical random effects in group models that include many channels and subjects (e.g. here the non-selective group-level data). Nonetheless, a hierarchical random effects model is likely the best starting model in many analyses as it accurately reflects how the data were collected and grouped together across channels and subjects. While there was usually minimal impact of random effects on the [mean] estimates of the beta weights, the inclusion of random effects did alter the significance level of some cluster-level beta weights. For example, a fixed effects-only model found that a broadband power face-selective group-level data cluster was significant only for category, but the mixed effects model with channel ID found this cluster was significant for both category and novelty; the mixed effects model with channel ID also fit the data better. These results highlight the importance of including appropriate random effects and how this can change the significance and interpretation of the results.

Finally, we analyzed high-gamma burst signals using CBPT with GLMEs to demonstrate the flexibility of our approach in analyzing nonlinear data. We expected these results to be similar to the broadband power data since high-gamma signals and broadband power are both thought to correlate with firing rates (Manning et al., 2009), but there are subtle differences beyond the scope of this paper. Broadly speaking, the results from analyzing the high-gamma burst signals using CBPT with GLMEs produced highly similar results to the results from analyzing broadband power signals using CBPT with LMEs, but there were more novelty-selective channels in the high-gamma burst signals which resulted in significant group-level novelty effects as well. Importantly, at the channel-level, there was good overlap in the category selectivity from extracted broadband power signals and high-gamma bursts, but there was less overlap between these extracted signals and ERPs. These results demonstrate how our proposed framework naturally extends to nonlinear models allowing easier access to a diverse array of neural features such as oscillatory bursts, which may reflect different types of neural processes.

### 4.4 Interpreting the Time Course of Fixed Effects and the Localization of Results in CBPT

An important feature of our approach is the ability to extract the time course of beta weights for all fixed effects, which has not been clearly demonstrated in previous applications of CBPT. We specifically focused on category-selective beta weights for individual-channel and group-level models from each of the 3 neural data signals. Analyzing the group-level beta weights, we observed that category selectivity emerged earlier when viewing faces than houses, especially in the broadband power and high-gamma burst signals. We confirmed these results by analyzing peak latencies from beta weights extracted from both individual channels and cross-correlation analysis of group-level beta weights, though the latency differences in the group-level data were smaller than the latency differences in the individual channels. These results are consistent with behavioral and EEG study findings showing that faces evoke higher amplitude and lower latency ERPs than objects in neurotypical people (McPartland et al., 2010; Webb et al., 2006). Nonetheless, it is important to understand the limitations of the proposed method and not to overinterpret the timing differences of beta weights for the several reasons discussed below.

Even assuming that we can reliably test the differences in the beta weight time courses, LMEs and GLMEs are statistical models that explain the observable statistics of the data and not mechanistic models that explain how the observed data was generated. Therefore, we caution overinterpretation of the beta weight time courses because it is possible to extract beta weight time courses that do not accurately reflect the true underlying distributions. Time courses of fixed effects will likely deviate more from the true underlying effects with greater trial bias, more noise, fewer observations, etc.

After accounting for the caveats above, beta weight time courses could be a powerful tool in analyzing latency differences at the channel-level and group-level data, but further simulations are necessary to determine the reliability of using beta weights to test effect latencies compared to previously used methods specifically designed for testing latency effects such as other permutation techniques, bootstrapping, and time-warping (J. Miller et al., 2009; Zoumpoulaki et al., 2015). If the time course of fixed effects is of great importance, we strongly recommend follow up analyses using alternative methods–such as granger causality–that are specifically designed to test latency differences in neural data to confirm the beta weight latency differences.

Somewhat related to the beta weight time course are the location of the significant time windows identified by CBPT. Unfortunately, CBPT has poor FWER control pertaining to the localization of effects in time (Maris & Oostenveld, 2007). In general, CBPT using LMEs and GLMEs should be no different than CBPT using other statistical models. Therefore, significant time windows should be interpreted as an *estimate* of when there is a significant effect. We can only say that with 100%*(1-α) certainty, the significant effect starts by the beginning of the detected cluster and lasts until at least the end of the detected cluster (Groppe et al., 2011b). Any attempt to interpret significant time window results otherwise is not statistically appropriate (Sassenhagen & Draschkow, 2019). There are alternative statistical approaches that can control better for FWER including FDR control methods (Frossard & Renaud, 2022; Groppe et al., 2011a), but many of these methods trade off statistical power (type II error) for better type I error control. Additionally, CBPT with minimum cluster sizes often miss smaller clusters which may still represent meaningful effects. This drawback was potentially observed in our analysis of novelty-selective ERP group-level data. In sum, the timing of a significant cluster is only an estimate of the time course of significant effects and should not be interpreted by itself.

### 4.5 Alternatives to the Proposed CBPT Methodology

There are also additional methods for analyzing data collected from experiments with many fixed effects. In particular, methods using FDR control like the Benjamini-Hochberg and Benjamini-Yekutieli procedures are relatively easy to implement, have superior FDR control, and can easily be used with multiple regression-type models (Benjamini & Hochberg, 1995; Benjamini & Yekutieli, 2001; Groppe et al., 2011b). However, since effects in neurophysiological data are typically broadly distributed–in time, space, and/or frequency–CBPT typically offers greater statistical power. Additionally, FDR methods applied to individual time points can produce intermittent significant effects which then have to be processed further to identify a good time window for subsequent analyses. Ultimately, one has to determine whether to control for FDR at the cost of statistical power or use CBPT at the cost of the ability to localize effects.

We chose to define clusters for all fixed effects together. However, it is likely that in many experiments, fixed effects will have little or no overlap in time. An alternative approach for this situation is to define clusters based on the unique combinations of potentially significant effects at each time point; in the dataset considered here, this would result in each time point being significant for either nothing, image category, image novelty or image category and novelty. Further work is necessary to determine the most robust method(s) for a wide variety of experimental scenarios.

Regardless of the specific methods used in CBPT with LMEs and GLMEs, there are a few important things to always consider. First, models should always include all reasonable fixed and random effects reflecting the experimental design, data collection process, and grouping of data. Second, α should always be corrected by the number of fixed effects being tested, and failure to do so will lead to increased FWER. However, alternatives to Bonferroni corrections like a Bonforroni-Holm correction could be employed especially when analyzing a large number of fixed effects. However, appropriately applying these methods across clusters with different numbers of significant fixed effects could be very difficult. Third, testing of cluster-level significance should be done on all fixed effects simultaneously to control for FWER. In sum, all fixed and random effects should be analyzed together in experiments with multiple fixed effects and when data is grouped together. Failure to do so may lead to increased type I errors, type II errors, and FWER.

### 4.6 Extension of CBPT with LMEs to 2D Data

A natural extension of the proposed method is to apply CBPT using LMEs to 2D time-frequency or 2D time-space data. Many authors have applied LMEs or CBPT to spectral data before (Domenech et al., 2020; Jacobs et al., 2006; Novembre et al., 2019) and in at least one instance have used CBPT with LMEs to analyze human intracranial data in an experiment with one fixed effect (S. Chen et al., 2021). The largest concern with 2D data analysis is that FWER increases with the number of comparisons made. Regardless, the use of CBPT with LMEs to analyze 2D data should be at least as robust as CBPT with other linear models like t-tests.

### 4.7 Designing Better Experiments

Recent years have seen neuroscience move towards more naturalistic, ethologically-valid experimental paradigms. Our CBPT-GLME framework presents a powerful tool for these more complex designs and varying data types (Alday et al., 2017). While well balanced experimental designs will always produce the best results, they are not always practical or compatible with naturalistic or individualized designs. For example, certain kinds of experiments may titrate difficulty to drive accuracy to a desired level. We show in simulations that LMEs and GLMEs can handle trial biases but benefit from higher trial counts like all statistical models. There are however certain design limitations which cannot be circumvented by our statistical framework, such as block-design. In the dataset analyzed here, the experiment had an ABB design for image novelty. Unfortunately, we found that neural responses varied across blocks and image novelty; therefore, we could not determine whether the responses were modulated by the number of times an image was shown and/or the number of blocks participants completed. A better design would be either an ABAB design or a mixed block design where novel images were intermixed with repeated images.

### 5.0 Conclusions

In this paper we combined the flexibility of LMEs and GLMEs with the power of CBPT to analyze various types of neural signals. We demonstrated through simulation and empirical analysis that CBPT with LMEs and GLMEs is a statistically robust method for analyzing time series data collected from experiments with multiple fixed effects and random effects. Further, we combined multi-step hypothesis testing and CBPT to determine if individual channels’ responses were modulated by task-related variables as well as pseudo-populations of functional groups of these channels. Additionally, we showed that CBPT with LMEs and GLME can directly extract the time courses of fixed effects, which can be useful in comparing differences in functional or anatomical groups of data. We believe the methods presented in this paper will be of great use to the neuroscience community in analyzing time series in a wide variety of experimental designs and across various data types, including ECoG, iEEG, EEG, MEG, spikes, and fMRI.

## Abbreviations

BB: broadband power
CBPT: cluster-based permutation tests
EEG: Electroencephalography
ECoG: Electrocorticography
ERD: event-related data

○ A general term that includes ERPs, broadband power data, and high-gamma bursts aligned to events.
ERPs: Event related potentials
iEEG: Intracranial Electroencephalography
FDR: False Discovery Rate
FWER: Familywise Error Rate
GLM: Generalized Linear Model
GLME: Generalized Linear Mixed-Effects Model
LLR: log-likelihood ratio [test]
LFP: local field potentials
LM: Linear Model
LME: Linear Mixed-Effects Model
MEG: Magnetoencephalography
sEEG: invasive stereo/depth EEG

## Acknowledgements & Funding Sources

The research reported here was supported by the National Institute of Drug Addiction under award # 5K23DA050909 (A.B.H) and by the Brain & Behavior Research Foundation under NARSAD Young Investigator Award # 28426 (A.B.H). Research reported in this publication was supported by the University of Minnesota’s MnDRIVE (Minnesota’s Discovery, Research and Innovation Economy) initiative (D.P.D).

## Declaration of interest

The authors have no conflicts of interest to declare.

## Author contributions

S.D.K, A.B.H, and D.P.D designed this study. S.D.K. carried out the simulations and empirical analysis.

S.E.S. provided expertise in biostatics and interpretation of the statistical results.

K.J.M. provided the ECoG data and advice about its interpretation.

All authors helped write the paper.

## Supplementary Figures

**Supplementary Figure 1:**
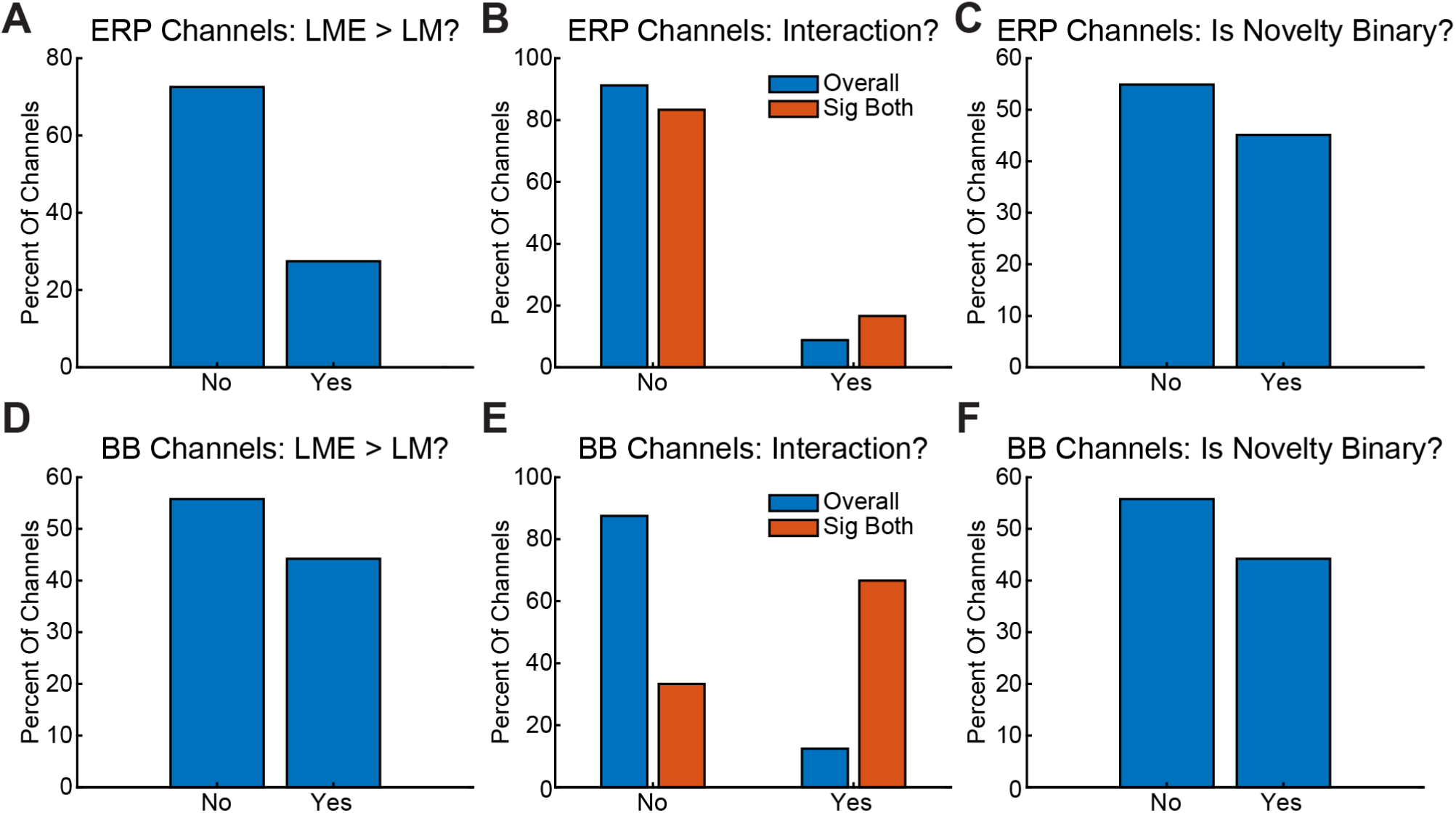
Alternative LME Models for the Predefined Time Window Analysis. We asked whether several different alternative models fit individual significant channels’ data better than a simpler LME using a LLR test (α = 0.05). Top row is for ERP data while the bottom row is for the broadband power (BB) data. A/D) Percent of channels that benefited from a mixed effects model (LME) over a fixed effects-only model (LM). B/E) Percent of channels that benefited from an interaction term between category and novelty. Blue bars are for all channels while orange bars are for channels selective for both category and novelty. C/F) Percent of channels that benefited from a fixed effect that represents the number of image presentations (i.e. 1, 2, and 3) compared to a model that represents binary image novelty (i.e. 0, 1). Note the number of times an image was presented correlated 1:1 with block number so results are difficult to interpret further.

**Supplementary Figure 2:**
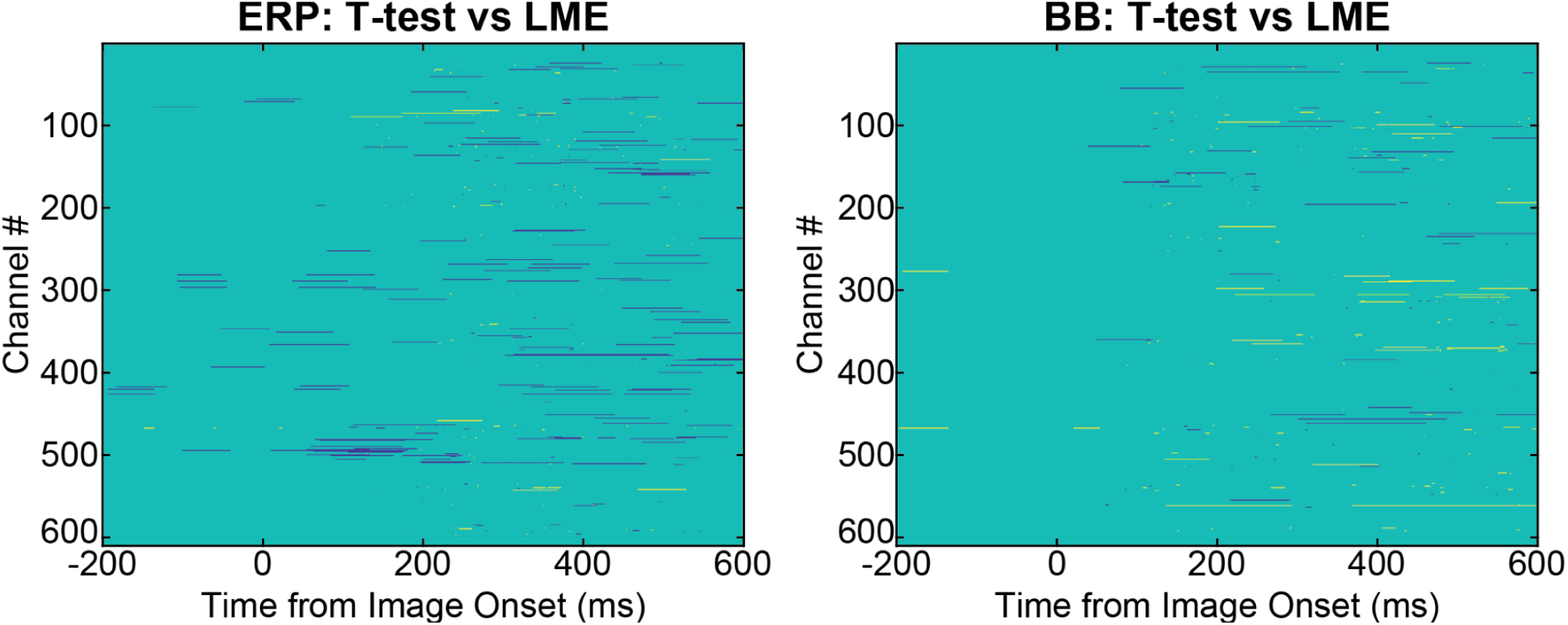
Comparison of Results from CBPT with LMEs and CBPT with Separate t-tests. Difference in results between CBPT with separate t-tests and CBPT with LMEs applied to ERP data and broadband power (BB) data. Gold pixels represent time and channels for which only a CBPT with separate t-tests had significant results after cluster-based correction for either fixed effect, while blue pixels represent time and channels for which only a CBPT with LMEs had significant results after cluster-based correction for either fixed effect. Green pixels indicate where the results were the same. Channel-level congruency was higher for broadband power data (95.1%) than the EPR data (87.5%).

**Supplementary Figure 3:**
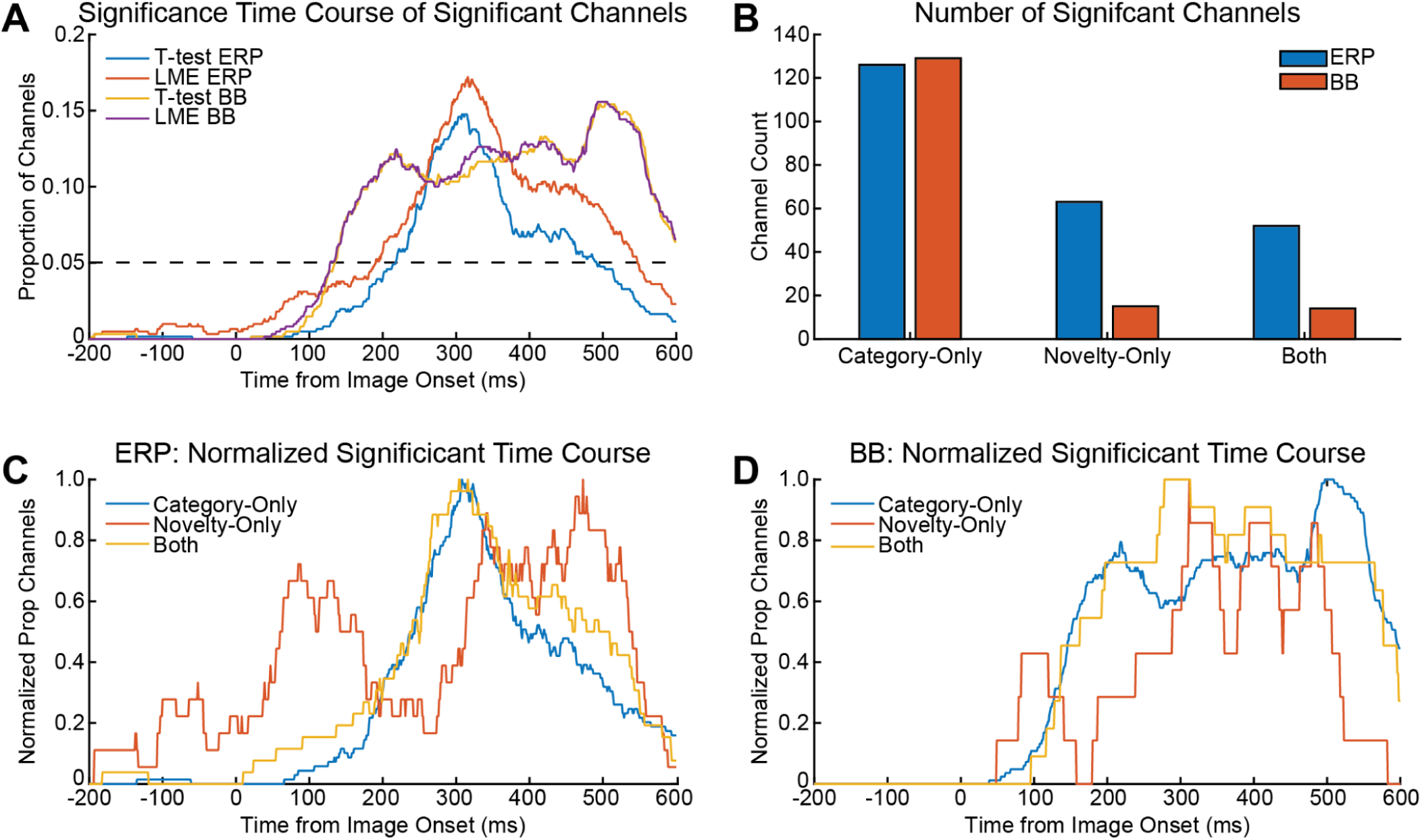
CBPT Time Course of Significance for ERPs and Broadband Power (BB) data. A) Proportion of channels identified as having a significant cluster over time for all data types and statistical models. As expected, most significant time points occurred 100 ms after image onset or later. B) Number of channels that have significant clusters identified with a LME for different data types and different kinds of selectivity. C) Normalized time course plots by selectivity from the ERP data analyzed using a LMEs suggests novelty selectivity occurs before and after category selectivity. D) Normalized time course plots by selectivity from the broadband power data analyzed using a LMEs suggests novelty selectivity co-occurs with category selectivity. *Note, broadband power data was smoothed with an 80 ms gaussian half-width window which likely explains why the time courses of significance for the broadband power data appears wider than the ERP data.

**Supplementary Figure 4:**
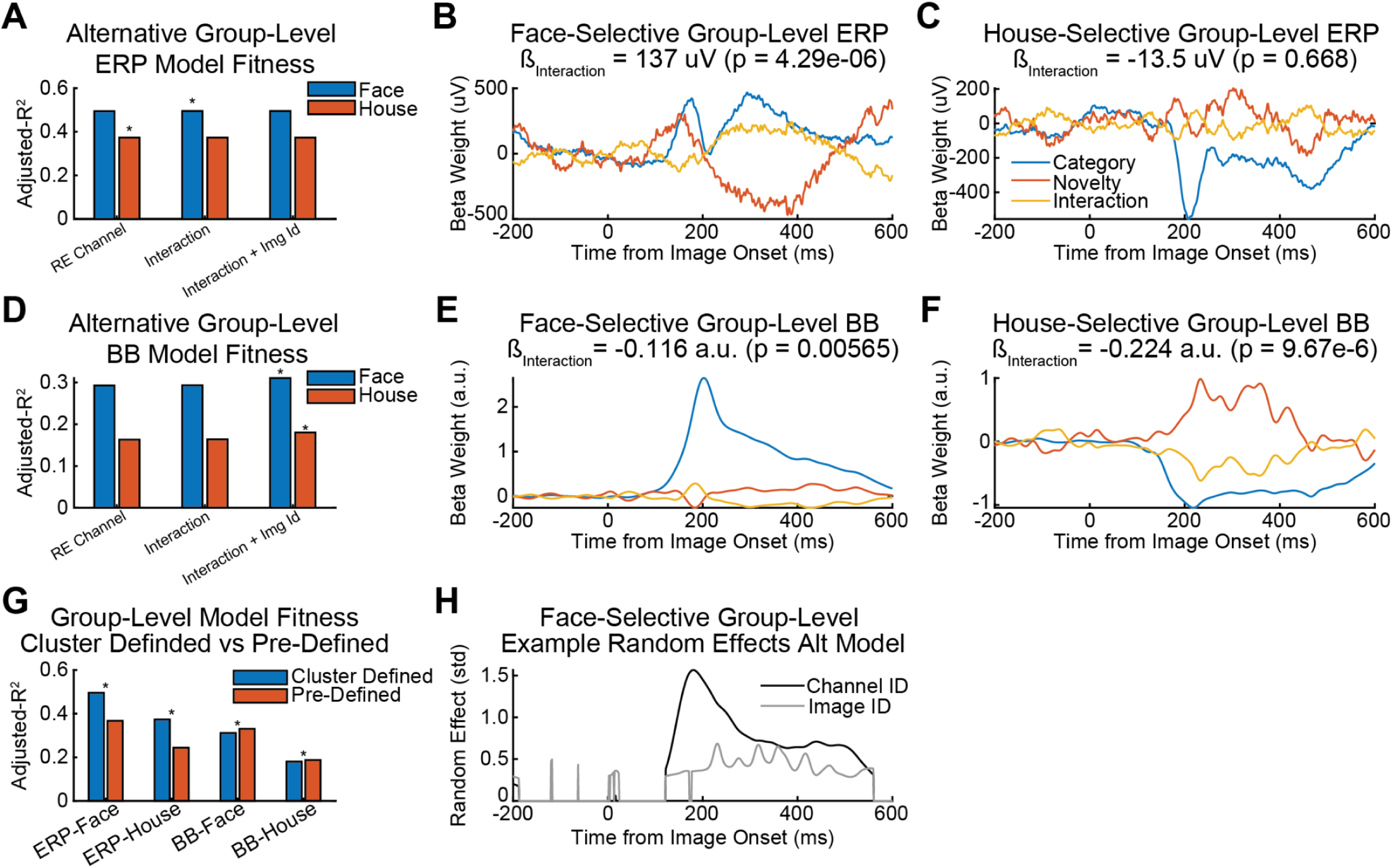
Alternative Group-level Model Testing. A) Adjusted-R^2^ for alternative models for the ERP group-level data with and without interaction terms. Asterisks indicate the best models according to a LLR test (p < 0.01/3). Only the face-selective group-level model benefited from an interaction term. B/C) ERP group-level face-and house-selective beta weight time courses for the category, novelty, interaction terms. Titles contain interaction effect beta weight and its significance estimated from the largest significant clusters (see Figure 7 shaded regions). D-F) Same as A-C except for the broadband power (BB) data. The best models for the face-and house-selective group-level data contained an interaction term between category and novelty as well as a random effects term for image ID. G) Comparison of adjusted-R^2^ for the cluster-defined windows compared to the predefined window from 100 to 400 ms after image onset. LLR tests showed that the cluster-defined windows “fit” the data better for all group-level models compared to the predefined window (*, p < 0.01/3). H) Example time course of random effect intercepts (measured in standard deviation across groups) for the house-selective group-level broadband power model showing larger variation across random effect groups after image onset especially for channel ID; this pattern was similar in the other group-level models.

**Supplementary Figure 5:**
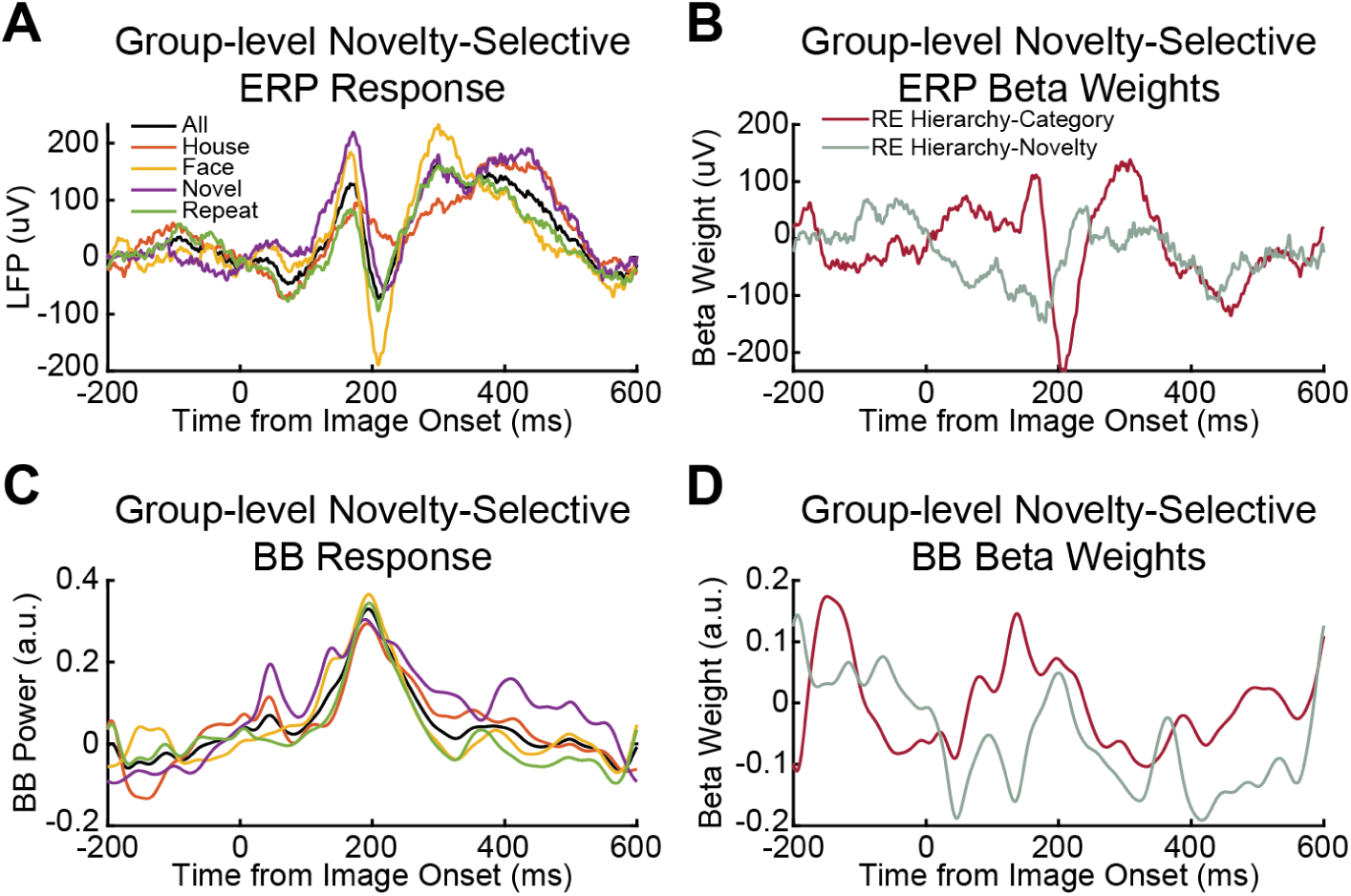
ERP and Broadband Power Novelty-selective Group-level Models. A) Averaged ERP activity for novelty-selective group-level data. B) Beta weights for the ERP novelty-selective group-level model from an LME with hierarchical random effects. C-D) Same as A-C except for the broadband power (BB) data. No time points or clusters were found to be significant for any of these data.

**Supplementary Figure 6:**
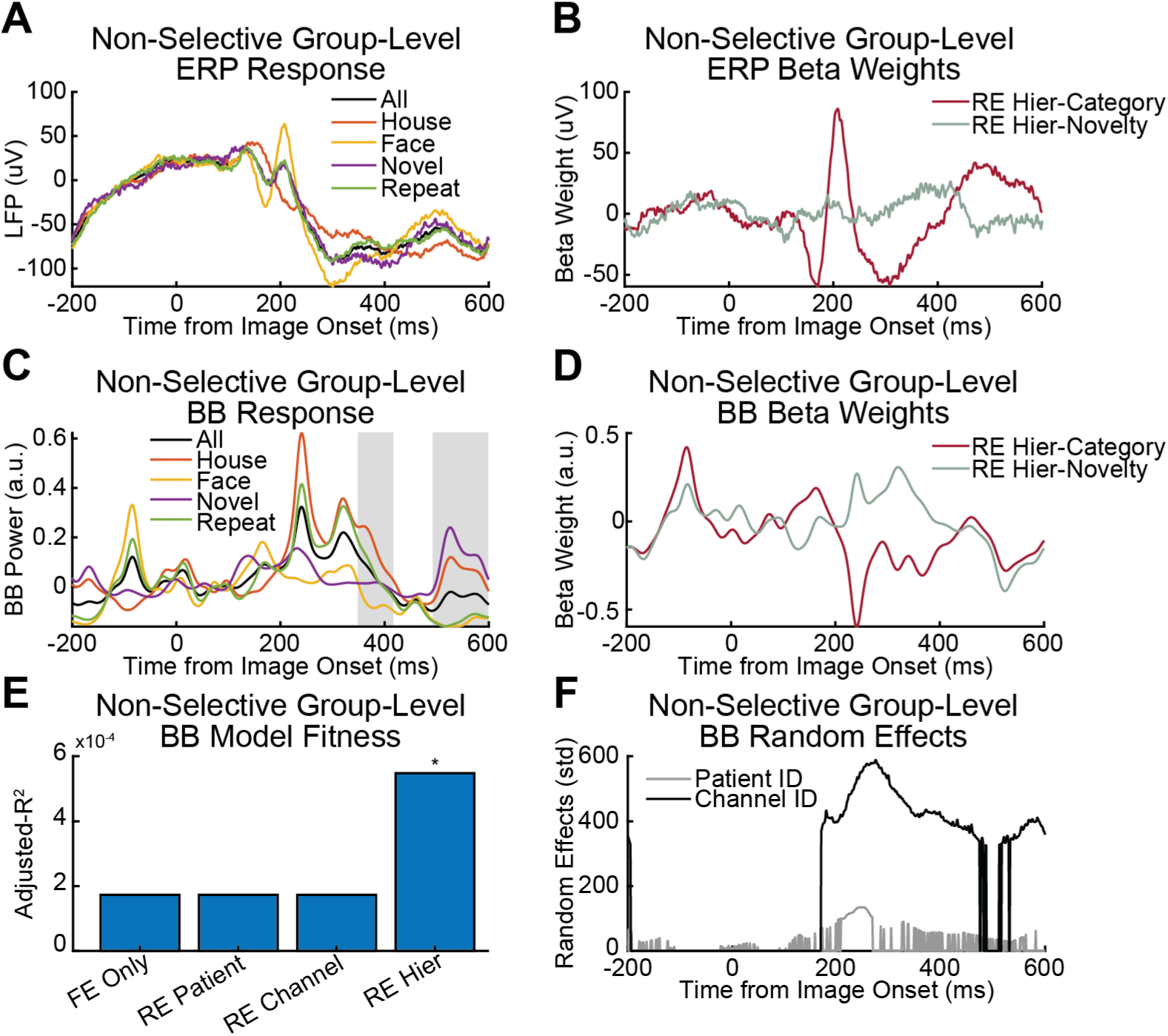
ERP and Broadband Power Non-Selective Group-Level Models. A) Averaged ERP activity for non-selective group-level data. B) Beta weights for the ERP non-selective group-level model from an LME with hierarchical random effects. C-D) Same as A-C except for the broadband power (BB) data. Gray shaded regions indicate significant clusters of time points found by using CBPT with LMEs. E) The best broadband power model for the significant clusters was the hierarchical random effects model (RE Heir) though no model explained a large amount of variance in the data. F) Visualization of time course of random effect intercepts (measured in standard deviation across groups) for the broadband power non-selective group-level model. Notice most of the variation in response is across channels and patients were observed after image onset; this pattern is similar in the other group-level models as well.

**Supplementary Figure 7:**
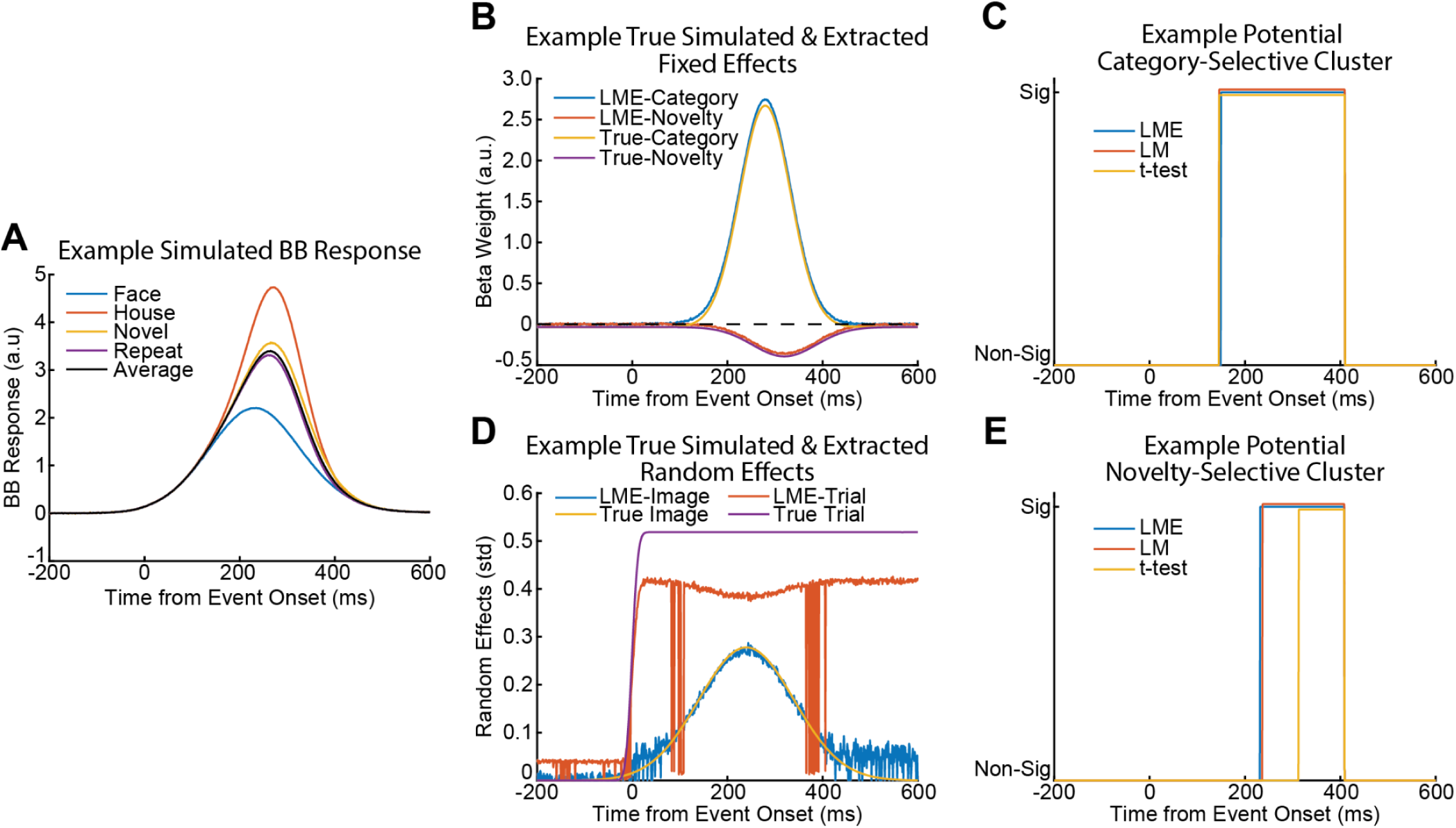
Example Simulated Broadband Power Signals. A) Example averaged response curves for face, house, novel, and repeat images. B) Example estimated LME beta weights and the true beta weights from the underlying distribution. Lines are slightly offset but overall the estimated beta weights from the LME were very similar to the true beta weights. C) Time course of potentially significant clusters for category-selectivity for each statistical model; results are identical across statistical models. D) Random effect time courses estimated by the LME and the true random effects added to the data. The variation in response across images was well estimated by the LME, but the variation in mean responses across trials was not. Note there is also random noise that is added to the model with a standard deviation of 0.05 which may be mixed in with the estimation of the trial-by-trial variation; note, some of the error in the attribution of variance to each random effect may be an artifact of how we generated the data rather than a shortcoming of LMEs. E) Time course of potentially significant clusters for novelty-selectivity for each statistical model; while there was a slight variation in cluster sizes between the potential cluster identified by the LME and LM, the cluster identified by the separate t-tests was substantially smaller.

**Supplementary Figure 8:**
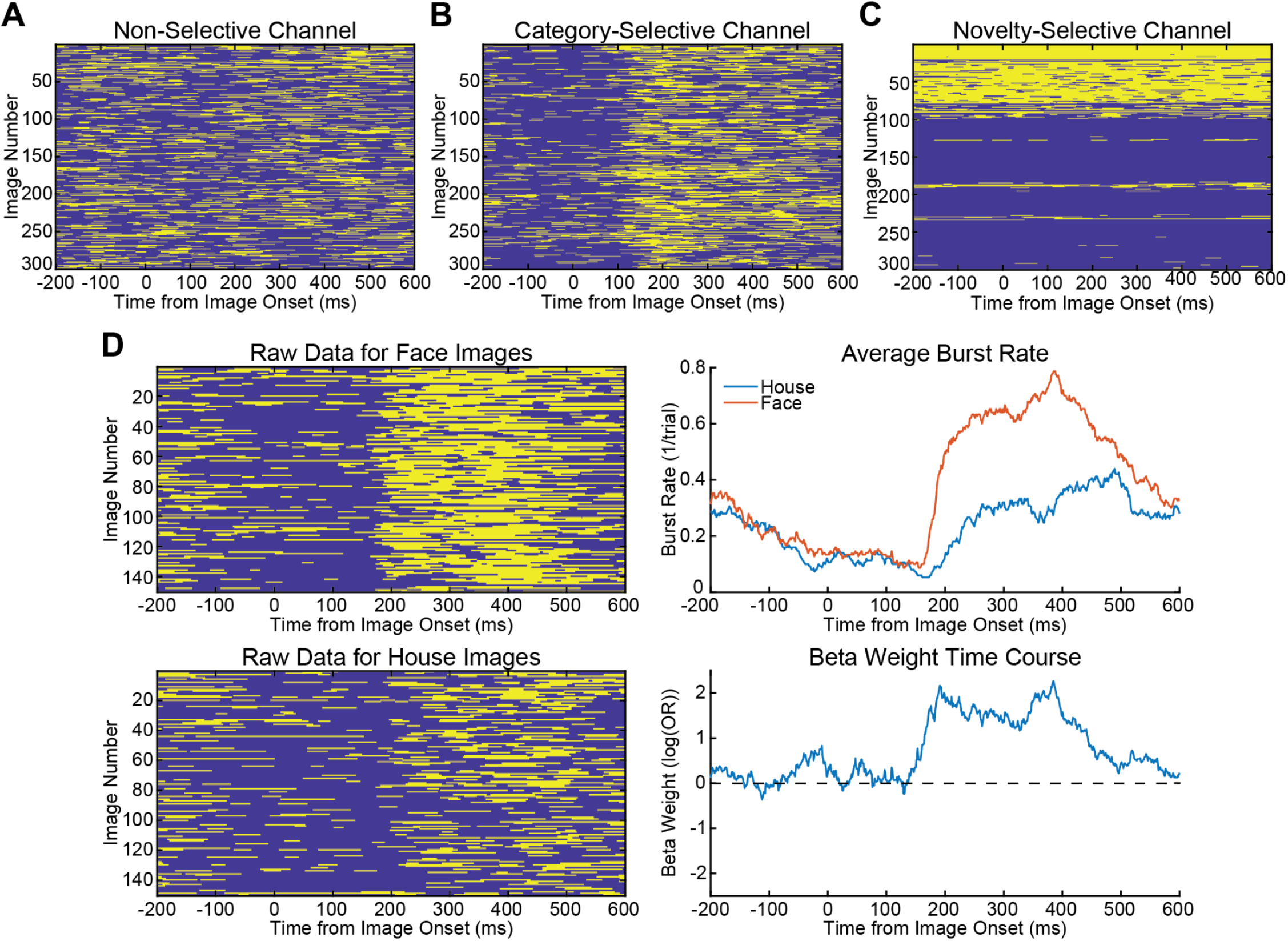
Example of High-Gamma Burst Data and Beta Weight Time Courses. A-C) Example raw high-gamma burst data for a non-selective, category-selective, and novelty-selective channel across all image presentations. Note the 1st 100 images are novel. Gold pixels represent the presence of a burst (1) and blue pixels represent the absence of a burst (0). D) Raw high gamma burst data from a different channel that was category-selective split into face images and house images, the average burst rate for these images, and beta weight time course extracted using a Logistical GLME. The beta weights’ units are in log(OR).

**Supplementary Figure 9:**
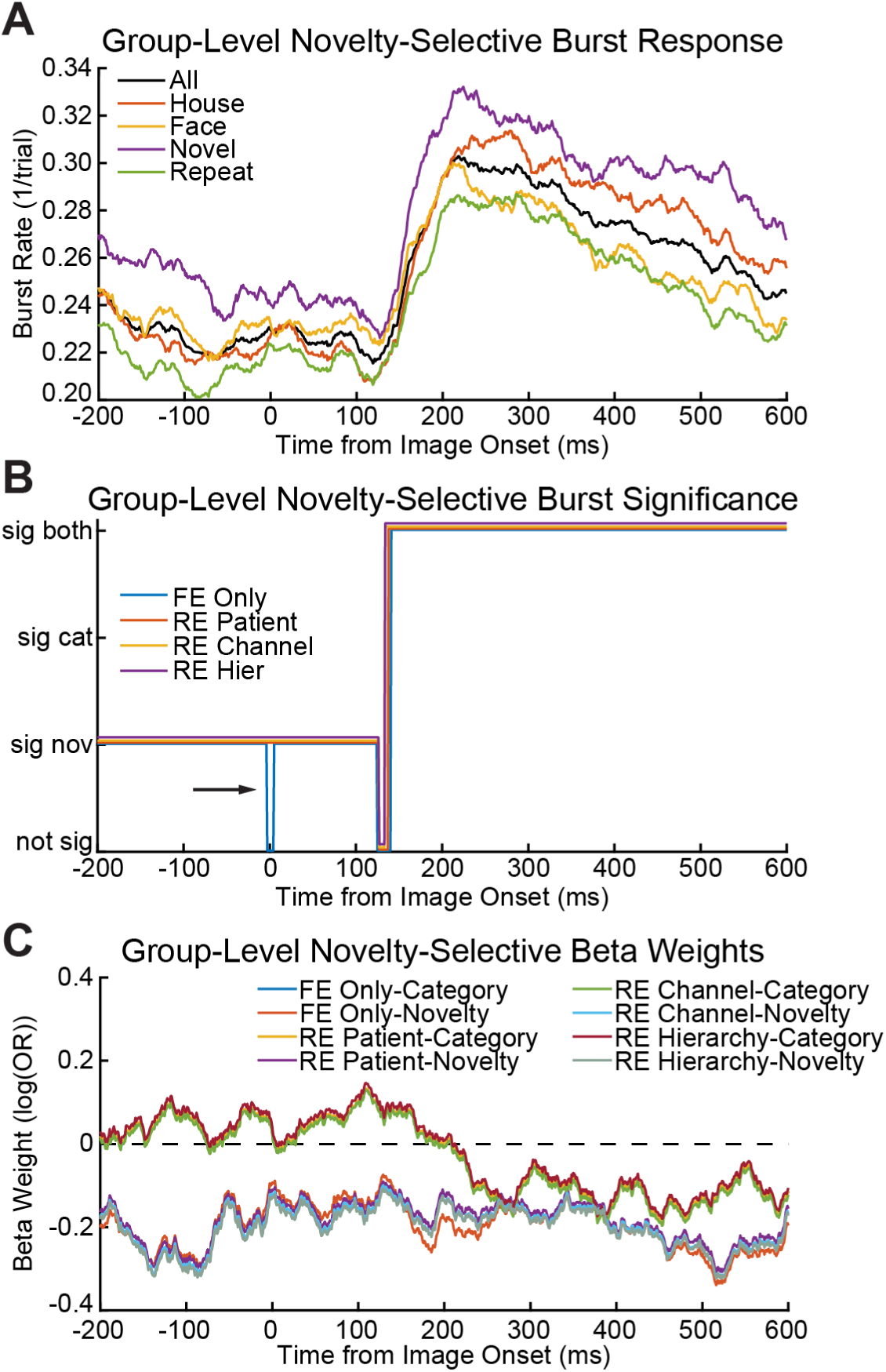
High-Gamma Burst Novelty-Selective Group-level Data Models. A) Average burst rate for novelty-selective group-level data. B) Significance time course for different random effects models. Note there is a small timing difference between the onset/offset of the identified clusters by model. The best model for the post-image onset cluster was the model that included patient ID as a random effect. C) Beta weight time course for different random effects models. Lines are offset slightly for visualization but there is an observable difference in the novelty beta weights over time for the model fixed effect only model (orange) which at least partially explains why the fixed effect only model split the pre-stimulus baseline period cluster into two (B, blue line, →).

**Supplementary Figure 10:**
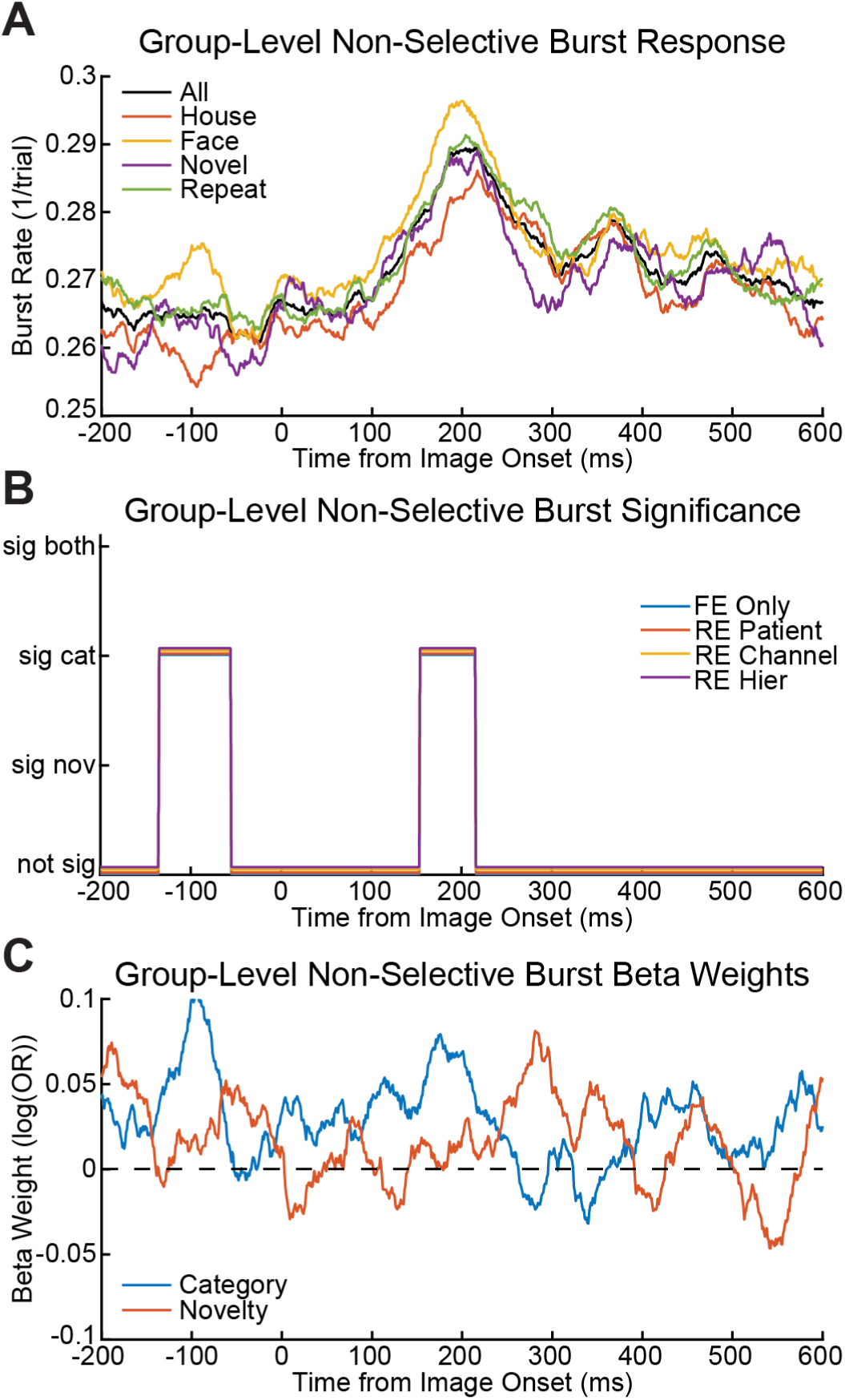
High-Gamma Burst Non-Selective Group-level GLME Models. A) Average burst rate for non-selective group-level data. B) Significance time course for different random effects models. The best model for the post-image onset cluster was the model that included channel ID as a random effect. C) Beta weight time courses from the model with channel ID as a random effect.

**Supplementary Figure 11:**
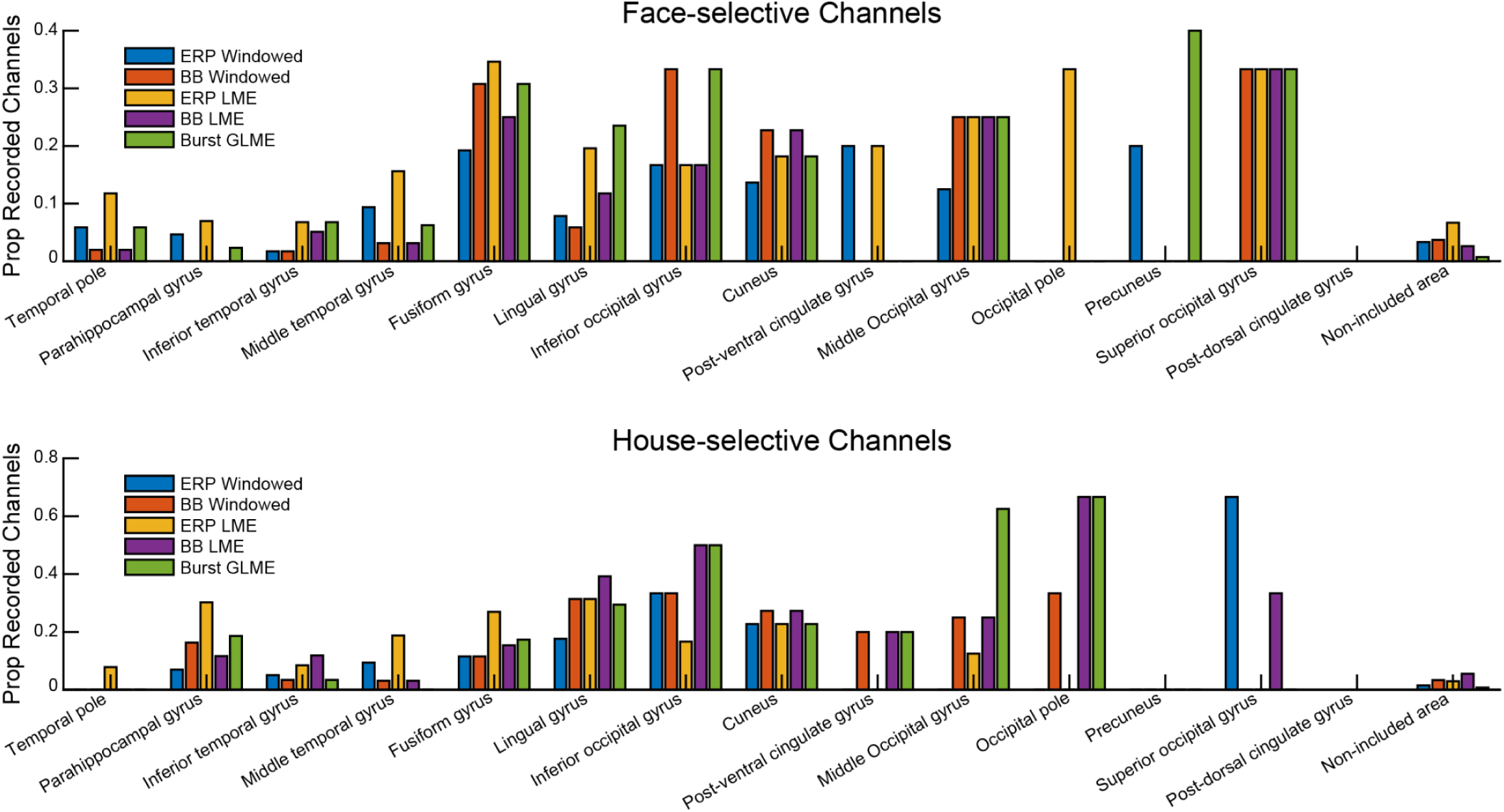
Anatomical Distribution of Category-Selectivity Across Different Neural Signals and Statistical Methods. Proportion of recorded channels significant for face-selectivity or house-selectivity across different types of neural signals and statistical methods.

## Supplementary Materials: Simulation Methods and Results

**Simulation Diagram 1:**
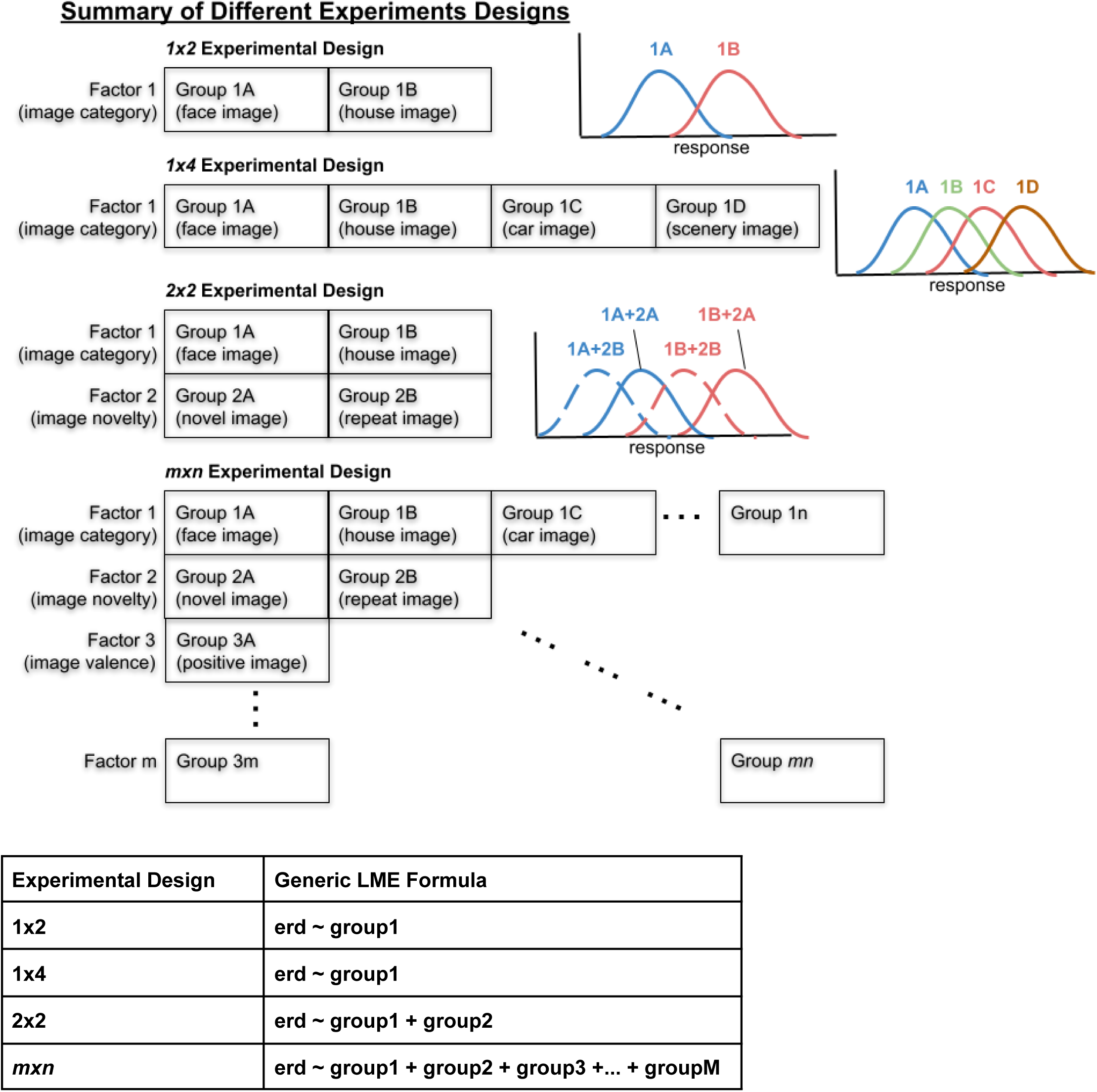
Different Experimental Designs. Illustration of different experimental designs, the associated distributions, and associated LME formulas. Note for 2×2 experimental designs, this diagram is usually shown differently such that Factor 1 is on the top and Factor 2 is on the side, but this design does not extrapolate well to *mxn* experiments; for 2×2 designs you get the following 4 groupings/levels: novel face, repeated face, novel house, and repeated house. Note, for *mxn* experimental designs the number of levels (n) does not have to be the same across factors (m).

## Simulation Methods

### Simulation Introduction

We conducted many simulations to validate the proposed CBPT method across a variety of experimental designs including from *1×2, 2×2*, and *1×4* factorial designs (**Simulation Diagram 1**). The simulations were used to determine if LMEs and GLMEs produce similar results to other commonly used statistical models over a wide parameter space. In particular, we wanted to ensure that we properly controlled for type I errors, type II errors, and FWER. Additionally, we wanted to determine how well commonly used simple models (e.g. t-tests) performed in the presence of random effects. Finally, we wanted to verify that CBPT combined with LMEs and GLMEs properly controlled for type I errors, type II errors, and FWER as well.

### Simple Linear Simulations

We used simulations to determine if LMEs produced at least as good statistical results as other commonly used linear statistical models. We ran three distinct sets of simulations reflecting different experimental designs: 1×2 simulations reflecting very basic two sample experiments, 2×2 simulations reflecting a 2×2 factorial design, and 1×4 simulations to determine how random effects influence results. We could not reliably model independent random slopes with anything less than a 1×4 design. Conceptually, these simulations were meant to reflect the distribution of data observed at individual time points in CBPT (see Step#1 in section 2.2.1).

For each simulation we generated 1000 distributions using the *normrnd* function over a large parameter space (**Simulation Table 1 & Simulation Table 2**). Simulation amplitudes (i.e. mean) and noise levels (i.e. standard deviation) were initially chosen somewhat arbitrarily for 1×2 simulations to produce a range of significance results, but subsequent simulations used parameters from previous simulations that produced sufficient proportions of significant and non-significant results. Simulation trial biases and trial counts were chosen to reflect a range of parameters commonly seen in many experimental designs. Trial bias is the proportion of trials associated with one type of condition; for example, a trial bias of 0.5 in a 1×2 simulation means that 50% of trials are associated with condition 1A and 50% of trials are associated with condition 1B.

We selected a subset of commonly used linear statistical models to compare to LMEs. To determine an LMEs significance, we used the p-value directly from the t-statistic(s). For 1×2 simulations we used two-sample t-tests, 1-way ANOVA, and a permutation t-tests (perm-t). For 2×2 simulations we used separate two sample t-tests, 2-way ANOVA, and separate two sample permutation t-tests. For the 1×4 random effect simulations we used a 1-way ANOVA, 1-way random effects ANOVA (RE-ANOVA), linear model (LM i.e. FE Only), permuted F-statistic (perm-F), and permuted sum-squared t-statistic (sum(*t-statistic*^2^)) from the LME. For all permutation tests, we used 1000 shuffles.

**Simulation Table 1:** Simulation Parameters for Simple Linear Distributions.

| **<u>Simple 1x2 Simulations</u>** | Amplitudes | Noise Levels | Trial Biases | Trial Counts |
| --- | --- | --- | --- | --- |
| Varying Amplitude | A: 10<br>B: 10:0.25:15 | 5.1 | 0.5 | 500 |
| Varying Trial Counts | A: 10<br>B: 11.25:0.25:12.25 | 5.1 | 0.5 | 100:100:1000 |
| Varying Noise Levels | A:10<br>B: 12.25 | 0.1:1:10.1 | 0.5 | 100:100:500 |
| Varying Trial Biases | A:10<br>B: 12.25 | 5.1 | 0.5:0.1:0.9 | 100:100:500 |
| **<u>Simple 2x2 Simulations</u>** |  |  |  |  |
| Varying Amplitude | A: 10 & 11.75<br>B: 5 & 0:1:10 | A: 5.1<br>B: 3.5 | A: 0.5<br>B: 0.7 | 500 |
| Varying Trial Counts | A: 10 & 11.75<br>B: 5 & 0:2:10 | A: 5.1<br>B: 3.5 | A: 0.5<br>B: 0.7 | 100:100:500 |
| Varying Noise Levels | A: 10 & 11.75<br>B: 5 & 0:2:10 | A: 5.1<br>B: 0.5:0.5:7 | A: 0.5<br>B: 0.7 | 500 |
| Varying Trial Biases | A: 10 & 11.75<br>B: 5 & 0:2:10 | A: 5.1<br>B: 3.5 | A: 0.5<br>B: 0.5:0.1:0.9 | 500 |

**Simulation Table 2:** Simulation Parameters for Simple Linear Distributions with Random Effects.

| **<u>Random Effects 1x4 Simulations</u>** | Random Intercept Amplitude | Random Slope Amplitude | Amplitudes | Noise Levels | Trial Biases † | Trial Counts | Number of Random Effects Levels |
| --- | --- | --- | --- | --- | --- | --- | --- |
| Varying Random Intercept | 0.1:1:4.1 | 0 | [10,11.75, 13.5, 15.25] | 5.1 | 0.25 | 200:100:500 | 5 |
| Varying Random Slope | 2.1 | 0.2:0.2:1 | [10,11.75, 13.5, 15.25] | 5.1 | 0.25 | 200:100:500 | 5 |

† A trial bias of 0.25 means uniform trial counts across the 4 conditions.

### Simple Nonlinear Simulations

We conducted two sets of simple 1×2 nonlinear simulations using binomial and Poisson distributions to ensure that GLMEs produce similar results to other commonly used nonlinear statistical models. We did not simulate greater than 1×2 experimental designs because separate analyses of fixed effects should still be less powerful than nonlinear statistical models that account for all fixed effects together and the presence of random effects should reduce power in nonlinear models that do not include random effects.

For the binomial data simulations, we varied probability, trial counts, and trial biases (Simulation Table 3). Binomial random distributions were generated with the Matlab function *binornd*. Similarly, for the Poisson simulations, we varied rate (λ), trial counts, and trial biases (Simulation Table 3). Poisson random distributions were generated with the Matlab function *poissrnd*. For the binomial simulations we compared the p-value from the t-statistic of logistic GLME to a χ^2^ test, Fisher’s exact test, and permutation of the sum(*t-statistic*^2^) from the GLME. For the Poisson simulations we compared the p-value from the t-statistic of Poisson GLME to a Kolmogorov-Smirnov test (ks-test), z-test on z-transformed data, and permutation of the sum(*t-statistic*^2^) from the GLME.

**Simulation Table 3:** Simulation Parameters for Simple Nonlinear Distributions.

| **<u>Simple 1x2 Binomial Simulations</u>** | Probability | Trial Biases | Trial Counts |
| --- | --- | --- | --- |
| Varying Probability | A: 0.5<br>B: 0.5:0.1:0.9 | 0.5 | 100:100:500 |
| Varying Trial Biases | A: 0.5<br>B: 0.7 | 0.5:0.1:0.9 | 100:100:500 |
| **<u>Simple 1x2 Poisson Simulations</u>** |  |  |  |
| | Rate ($\lambda$) | Trial Biases | Trial Counts |
| Varying Rate | A: 10<br>B: 10:0.5:12.5 | 0.5 | 100:100:500 |
| Varying Trial Biases | A: 10<br>B: 11 | 0.5:0.1:0.9 | 100:100:500 |

### Validation of Cluster-Level Permutation Methods

#### Validating sum(t-statistic^2^) as a Whole-Model Test Statistic

We were interested in identifying a robust whole-model test statistic for permutation testing that was not influenced by distribution parameters, especially the presence of random effects. Several whole-model permutation methods have been proposed in previous literature (M. Anderson & Braak, 2003; M. J. Anderson, 2001; Winkler et al., 2014). We chose to limit the test statistics and permutation methods to those we thought were easy to implement no matter the complexity of the model including the number of fixed effects and the presence of interaction terms. We calculated 3 test statistics: Sum F from an ANOVA, R^2^, and sum(*t-statistic*^2^). Further, we permuted the data using 3 different methods: full model permutation (Manly method), residual permutation ignoring random effects (ter Braak method), and a Freedman-Lane-style permutation accounting for random effects. We additionally tested the permutation of a fixed effects-only (FE only) model and then used R^2^ as a test statistic.

The Freedman-Lane method was developed to test partial correlation coefficients, but here we conduct a similar permutation test but do not account for either fixed effect in the calculation of the residuals. The sum of the F-statistics from an a 2-way ANOVA without interaction between conditions (MSA/MSE + MSB/MSE) is equivalent to the (MSA + MSB)/MSE, which is ratio of the total variance explained divided by the total unexplained variance; MSA, MSB, and MSE are the Mean Sum of Squares for fixed effect A, the Mean Sum of Squares for fixed effect B, and the Error Mean Sum of Squares, respectively Additionally, the *t*-*statistics* from a GLME is the ß/S.E.(ß) which is the mean estimated beta weight (ß) divided by the standard deviation of the estimate of ß. We squared the *t-statistic* to account for negative *t-statistics* and to create a one-tailed permutation test.

For these whole-model permutation test simulations, parameters were uniformly sampled from a subset of previously used parameters (**Simulation Table 4**). In total we generated 5000 simulations with 1000 simulations per random effect intercept amplitude.

**Simulation Table 4:** Simulation Parameters Whole Model Permutation Testing.

|  | Random Intercept Amplitude | Amplitudes Across Conditions | Noise Levels | Trial Biases | Trial Counts |
| --- | --- | --- | --- | --- | --- |
| **Whole-Model LME 2x2 Permutation Tests** | 0.1:1:4.1 | A:10:0.25:12.5<br>B: 3:7 | A: 5.1<br>B: 3.5 | 0.7 | 300 |

#### Validating Cluster-Level Model vs Cluster-Mass Statistic

We simulated 1×2 cluster-level permutation tests to verify the proposed cluster-level statistics were robust. Specifically, the most common method for testing cluster-level significance uses cluster-mass statistics using the sum of the t-statistics from a t-test across the whole cluster. However, this method is inherently biased towards larger clusters. Instead, for normally distributed data we propose taking the mean of the data across time points within each cluster then fitting the mean data with a new LME; there is little difference between the mean and the sum except for the normalization by the number of samples. Furthermore, by taking the mean response for the cluster we combine two steps into one since we ultimately want a cluster-level model to describe the data. For binomial distributions we take the sum of the binary responses (0’s and 1’s) across time within a cluster, create a burst count distribution, and fit the burst count distribution with a Poisson GLME; we then compared the sum(*t-statistic*^2^) from the Poisson cluster-level model to a cluster-mass statistic using the sum of χ^2^ statistic across all time points in a cluster.

For these cluster-level permutation tests we generated monophasic response curves with variable half widths and fixed effect sizes. Like the previous simulations, we compared the congruency of the cluster-level permutation results using the cluster-mass statistics to the permutation results using the sum(*t-statistic*^2^) from the cluster-level models. For these simulations we used a wide range of parameters inherited from previous simulations (**Simulation Table 5**). Minimum cluster size was determined as the half-width of the simulated curve. Clusters were initially identified using the LME’s or GLME’s p-values from the t-statistic and then we computed cluster-mass statistics for those clusters using the other statistical model; this methodology is potentially biased but is meant to ensure the cluster-level LMEs or GLMEs are not producing false positives. We shuffled the data 1000 times for each permutation test.

For normal distributed data we simulated gaussian response curves with normal statistical properties at each time point. We also added a small amount of random trial noise to the response curves. We did not include random effects in these simulations as the traditional cluster-mass statistics would likely be inaccurate in the presence of random effects. For these simulations we generated a total of 55,000 simulated response curves with 1000 simulations for 11 different fixed effect sizes and 5 different trial counts. We compared permutation results using the sum(*t-statistic*^2^) from the cluster-level LME to the permutation results from the cluster-mass t-statistics from a t-test.

For binomially distributed data we first generated gaussian probability response curves with maximum responses between 0.5 and 1 and then generated random binomially distributed data around those mean probabilities. For the GLME simulations we generated a total 25,000 simulated response curves with 1000 simulations for 5 different fixed effect sizes and 5 different trial counts (**Simulation Table 5**). For the binomial simulations we compared permutation results using the sum(*t-statistic*^2^) from the cluster-level GLME to the permutation results from the cluster-mass χ^2^ statistic from a χ^2^ test.

**Simulation Table 5:** Simulation Parameters for Cluster-Level Permutation Testing.

| | Random Intercept Amplitude | Amplitudes Across Conditions | Noise Levels | Trial Biases | Trial Counts | Halfwidth (Mean ± std) | Random Trial Noise |
| --- | --- | --- | --- | --- | --- | --- | --- |
| **LME 1x2 Cluster Permutation Tests** | 0 | A: 10<br>B: 10:0.25:12.5 | 5.1 | 0.5 | 100:100:500 | 50±25 | 5.1/2 |
| **GLME 1x2 Cluster Permutation Tests** | 0 | A: 0.5<br>B: 0.5:0.1:0.9 | N/A | 0.5 | 100:100:500 | 50±25 | 0.02 |

#### Validation of Multiple Comparisons Correction Across Fixed Effects

We wanted to determine the best method for controlling for multiple comparisons across numerous fixed effects. We started with a Bonferroni correction across fixed effects because it was easy and straightforward to implement, but we were concerned that a Bonferroni correction was too conservative and that a Bonferroni-Holm correction would be substantially more powerful even though it would be more difficult to implement.

In traditional CBPT (Maris & Oostenveld, 2007), there is only one fixed effect; therefore, multiple comparisons testing corrects across potential clusters within the same fixed effect. However, in the presence of multiple fixed effects, multiple comparisons testing can correct across potential clusters within as well as across fixed effects. Note, these simulations were not designed to test within fixed effects corrections where noise and responses can covary across potential clusters. In the proposed methodology, the significance of multiple fixed effects are being considered simultaneously, and not correcting for multiple fixed effects could lead to an increase FWER as potentially seen in (Bianchi et al., 2019) where each fixed effect was analyzed separately.

To understand how various multiple comparison methods perform with CBPT, we carried out a simple 4x(1×2) simulation in which we generated 4 independent 1×2 normal distributions simultaneously with statistics drawn from a limited range of parameters used in previous simulations (**Simulation Table 6**). We chose to simulate 4 independent fixed effects because this was the number of fixed effects in (Bianchi et al., 2019), and a Bonferroni correction becomes more conservative with a larger number of comparisons. In total we generated 10,000 sets of 4x(1×2) distributions with ∼50% of simulations forced to have the same amplitude across distributions in the set and the other ∼50% having uniformly random amplitudes. We then employed various techniques for multiple comparisons corrections: no correction, Bonferroni, Bonferroni-Holm, Max-T, and Max-T with Bonferroni correction; Max-T is meant to simulate the maximum t-statistic from cluster-based permutation tests but here we did not square the t-statistic. All analyses were done using t-tests for simplicity with α = 0.05.

**Simulation Table 6:** Simulation Parameters for Multiple Comparisons Correction.

| Simple 4x(1x2)<br>Simulations | Amplitudes | Noise Levels | Trial Biases | Trial Counts |
| --- | --- | --- | --- | --- |
|  | A: 10<br>B: [10 11 12 13] | 5.1 | 0.5 | 300 |

## Simulation Results

### Simple Linear 1×2 Simulation Results

**Simulation Figure 1:**
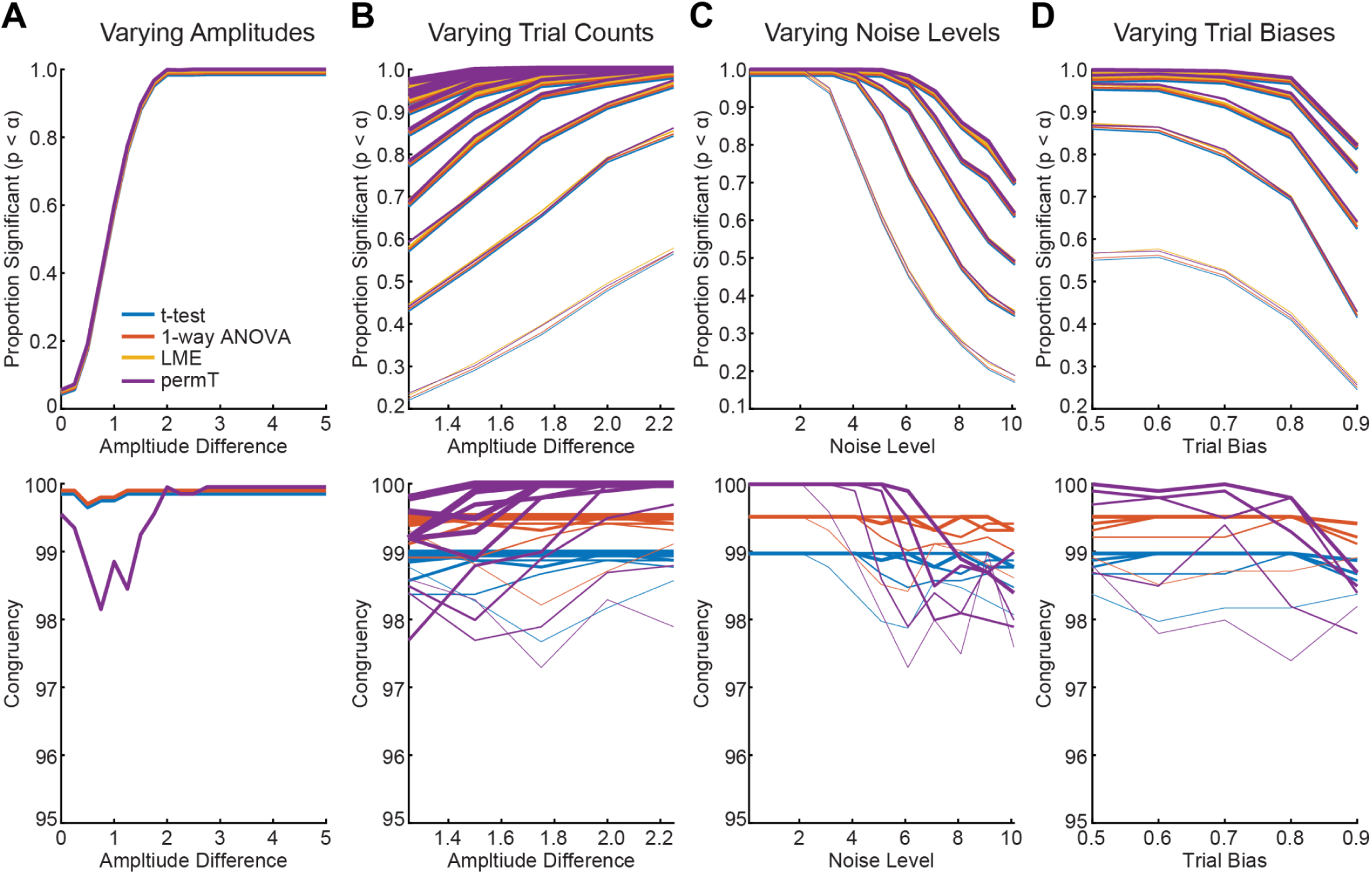
Simple Linear 1×2 Simulations. The proportion of simulations with significant results (p < α, α = 0.05) for various linear statistical models as well as the congruency of the result with the LME results. In general, results were very similar across statistical models for varying amplitudes (A), varying trial counts (B), varying noise levels (C), and varying trial biases (D). Note, lines were offset across statistical models since results were so similar; for the congruency panels, usually the maximum congruence was 100% or nearly 100%. For panels B-D thinner lines represent smaller trial counts (e.g. 100) and thicker lines represent larger trial counts (e.g. 500).

We started by simulating several simple distributions with a 1×2 factorial design. We tested the hypothesis that two group means (i.e. µ_1A_ vs µ_1B_ or 1 fixed effect) are different within a population of normally distributed observations. We wanted to compare the congruency (i.e. consistency) of results across commonly used linear statistical models to the results from LMEs. We varied several different parameters to create different types of normally distributed data including varying amplitude (i.e. mean), trial counts, noise (i.e. standard deviation), and trial bias (i.e. proportion of trials associated with a trial condition 1A). The different parameters were meant to simulate the various types of distributions of data observed at individual time samples encountered in the first step of CBPT with LMEs. In all simulations we generated 1000 random distributions and used 1000 permutations for the permutation t-test.

In general the 1×2 simulations showed the expected results, and results were highly congruent (97.3% to 100.0%) across different statistical models (**Simulation Figure 1**), with the permutation t-test (perm-T) showing slightly lower congruence (< 1%) with the LME results than ANOVA or t-test in certain simulations. Across all models, the probability of finding significant results (p < α, α = 0.05) increased with the amplitude difference across conditions (**Simulation Figure 1A**). Note the false positive rate for the 0 amplitude difference simulation was 5.6% to 5.8% which was not significantly different than 5% (χ^2^ proportion test, p’s ≥ 0.429). Similarly, the probability of finding a significant result increased with trial count (**Simulation Figure 1C**). Conversely, as noise levels increased the probability of finding a significant result decreased (**Simulation Figure 1E**), and as trial bias increased the probability of finding a significant result also decreased. While no strong pattern of congruency was observed across these parameters for any particular statistical model, there was some evidence that results were less congruent at borderline parameters (e.g. amplitude difference 1, **Simulation Figure 1B**) where there may be subtle differences in p-value estimates; results were also slightly less congruent across models in some simulations with high noise or trial biases especially at low trial counts (**Simulation Figure 1D and 1F**).

### Simple Linear 2×2 Simulation Results

We next simulated a similar set of simple distributions with a 2×2 factorial design. We tested the hypothesis that four independent group means (i.e., µ_1A_ vs µ_1B_ & µ_2A_ vs µ_2B_ or 2 fixed effects) were different within a population of normally distributed observations. Like the 1×2 design, we manipulated multiple parameter values to determine how different linear statistical models performed in comparison to LMEs. While results for the 2×2 simulations were often similar to the results to the 1×2 simulations, the separate t-tests showed detrimental significance biases not observed in LMEs or ANOVAs (**Simulation Figure 2**) consistent with the idea that running separate single analyses of multiple fixed effects in 2×2 factorial designs or greater leads to reduced statistical power. Across varying amplitude differences of the 2nd fixed effect (i.e. µ_2A_ vs µ_2B_), ANOVA results were highly congruent with LME results (≥ 99.7%), while separate t-tests and permuted t-tests showed lower congruence in some conditions (≥ 91.7%). For example, for the 2^nd^ fixed effect amplitude difference of 5, LMEs found significantly more simulations (88.3% vs. 83.6%, χ^2^ proportion test, p = 0.00249) with significant results than the separate t-tests (**Simulation Figure 2B**). Results were similar in other simulations with varying trial counts, noise levels, and trial biases (**Simulation Figure 3**). Separate t-tests and permutation t-tests were consistently underpowered compared to LMEs and ANOVAs, especially for low trial counts and high noise levels, while a 2-way ANOVAs performed similarly LMEs under all simulation conditions (congruence ≥ 98.3%).

Overall, the 2×2 factorial design simulation results suggest that separate analysis of multiple fixed effects is inferior to linear models that simultaneously account for multiple fixed effects together such as n-way ANOVAs and LMEs. Specifically, separate t-tests were significantly underpowered and were less likely to identify significant effects than LMEs or ANOVAs even though in 1×2 simulations there were no power differences between linear statistical models.

**Simulation Figure 2:**
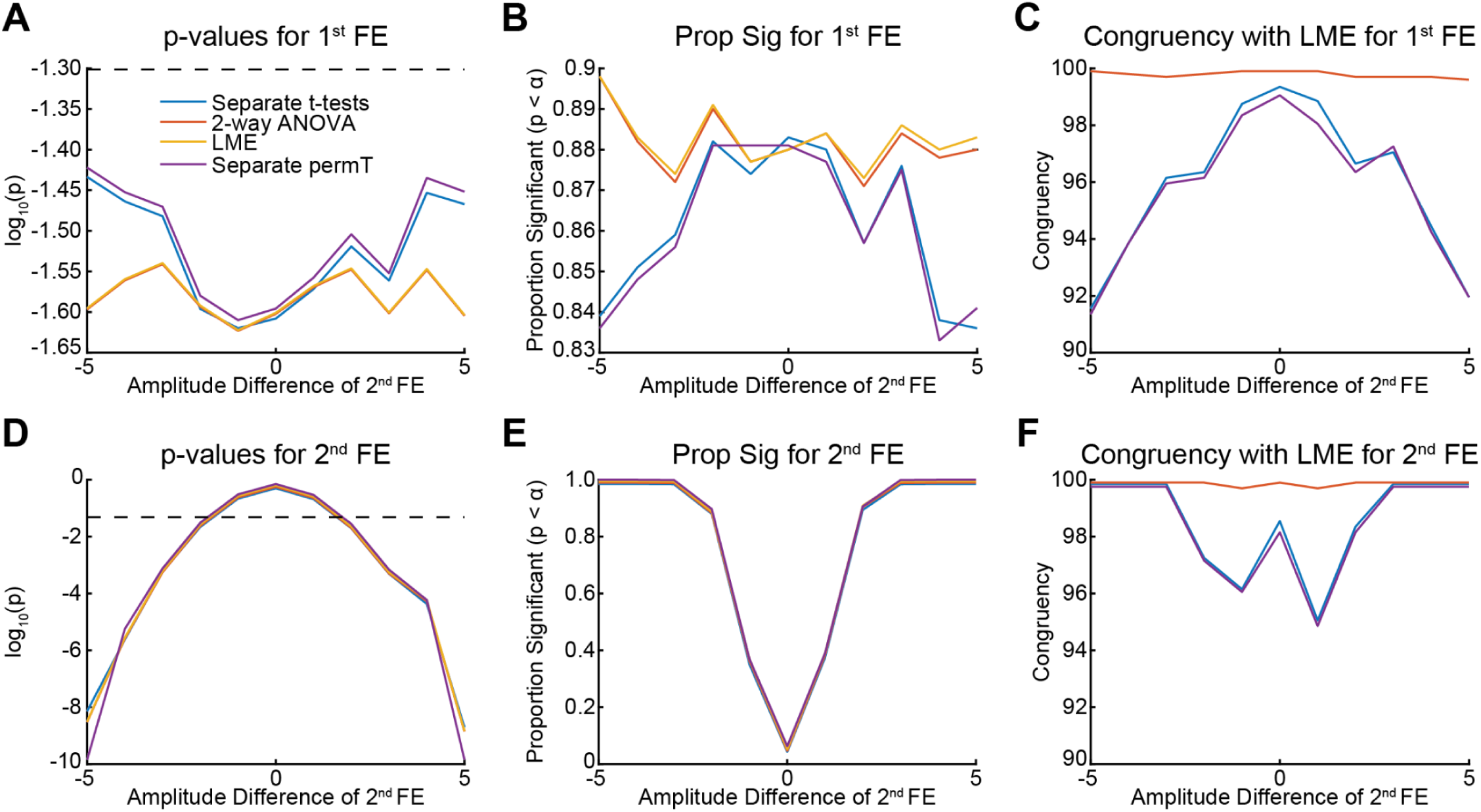
Simple Linear 2×2 Simulations for Varying Amplitude. The proportion of simulations with significant results (p < α, α = 0.05) for various linear statistical models as well as the congruency of the result compared to the LME results (top row 1st fixed effect, µ_1A_ vs µ_1B_; bottom row 2^nd^ fixed effect, µ_2A_ vs µ_2B_). A/D) Average log_10_(p-values) as function of the amplitude difference for the 2^nd^ fixed effect (FE); the amplitude difference was held constant for the 1^st^ fixed effect. B/E) Proportion of simulations identified with a significant result as a function of amplitude difference of the 2^nd^ fixed effect. C/F) Congruence of statistical models with LME as a function of the 2^nd^ fixed effect amplitude. *Note, lines were offset across statistical models since results were so similar.

**Simulation Figure 3:**
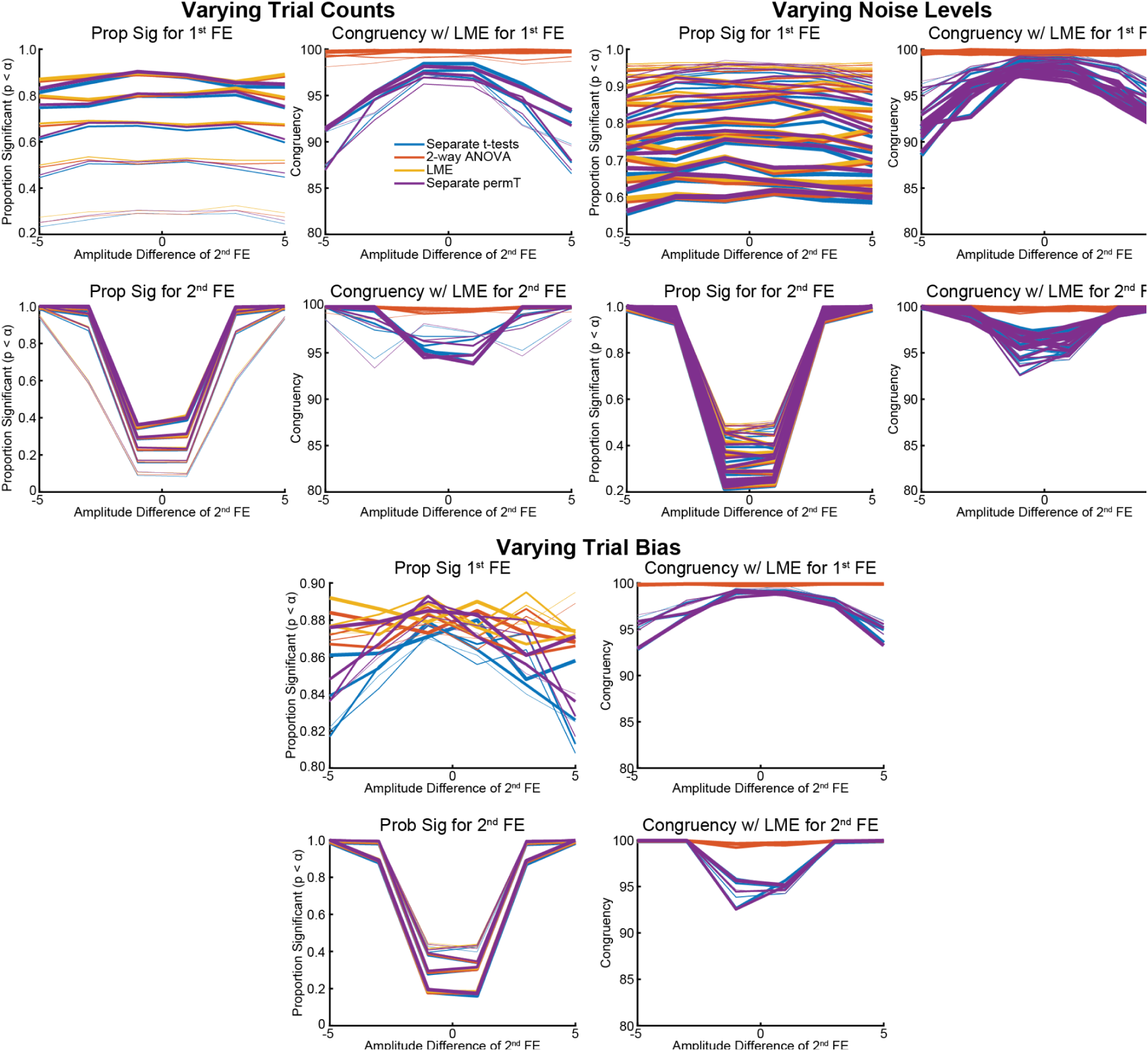
Simple Linear 2×2 Simulations for Varying Trial Counts, Noise, and Trial Bias. The proportion of simulations with significant results (p < α, α = 0.05) for various linear statistical model parameters as well as the congruency of the result compared to the LME results. We varied trial counts from 100 (thin lines) to 500 (thick lines) and found that the overall proportion of significant simulations varied drastically across trial counts; specifically, results tended to be less congruent for lower trial counts than higher trial counts. Similarly, noise levels from 0.5 (thin lines) to 7 (thick lines) for condition 2 drastically affected the proportion of significant results, and there was an apparent effect of noise levels on congruence as well. Trial biases from 0.5 (thin lines) to 0.9 (thick lines) affected the probability of finding significant results for the 2^nd^ fixed effect but not 1^st^, especially at lower amplitude differences; there was some effect of trial biases on the congruence of the results. The congruency between LMEs and 2-way ANOVAs was always between 98.2% and 100%, while the congruency between LMEs and separate t-tests varied from 86.7% to 100% depending on the simulation parameters.

### Simple Linear 1×4 Simulation with Random Effects Results

**Simulation Figure 4:**
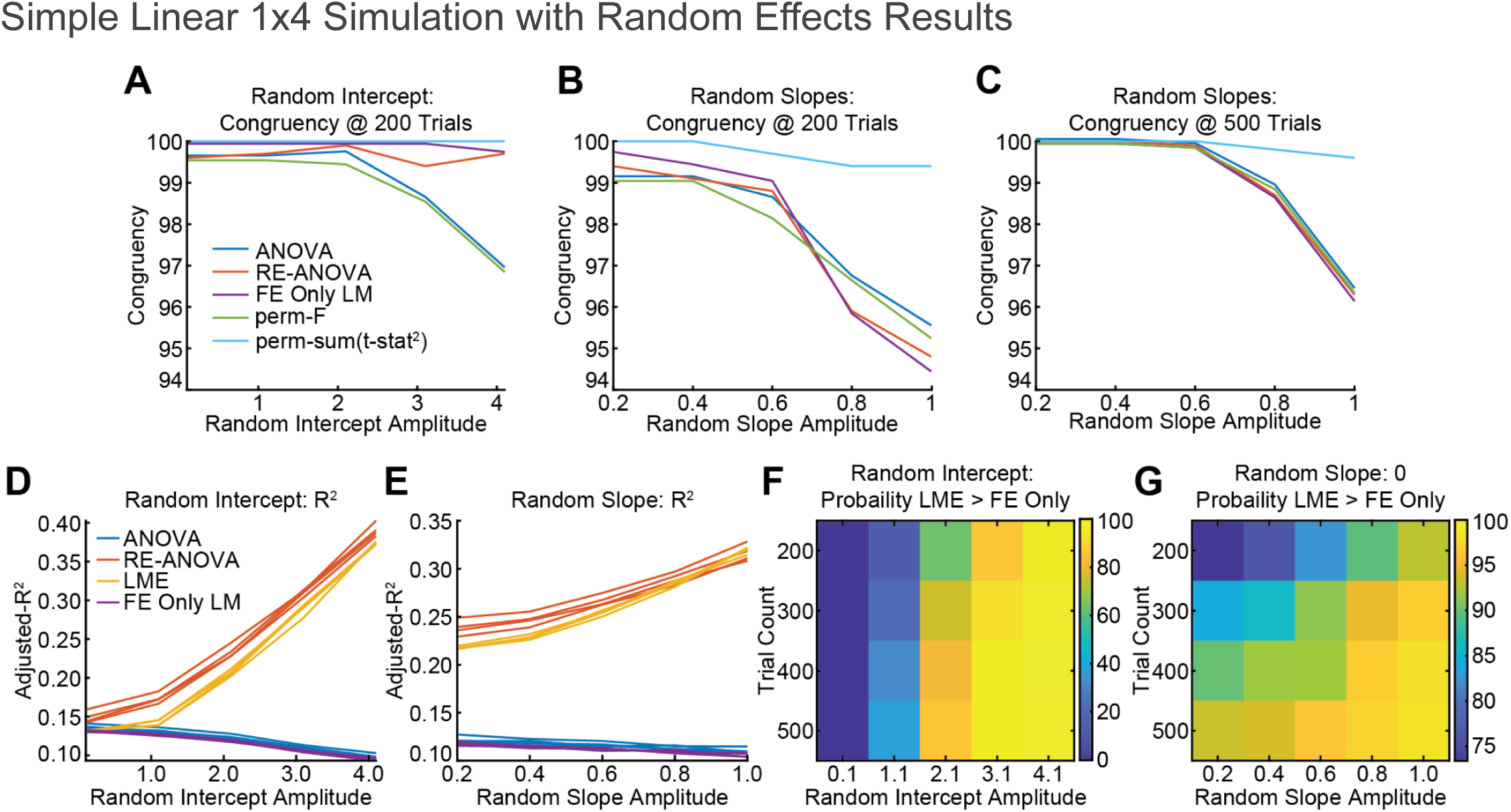
Simple Linear 1×4 Simulations with Random Effects. A) Congruence between various linear statistical models and LMEs in the presence of random intercept amplitudes for 200 trials. At 300 trials or more results were ≥ 99.9% congruent with LMEs for all statistical models. **B & C)** Congruence between various linear statistical models with LMEs in the presence of random slopes for 200 and 500 trials. **D)** Adjusted-R^2^ for all statistical models in simulations with different random intercept amplitudes. There was little evidence of trial counts affecting Adjusted-R^2^ (different lines). **E)** Adjusted-R^2^ for all statistical models for simulations with different random slopes and intercept amplitudes. There was little evidence of trial counts affecting Adjusted-R^2^ (different lines). **F)** The probability that LMEs were a better fit (LLR test, α < 0.05) for the data than fixed effects-only models for different random intercept amplitudes and trial counts. **G)** The probability that LMEs were a better fit (LLR test, α < 0.05) for the data than fixed effects-only models for different random slope amplitudes and trial counts. *Note subplots A-C had a small offset applied to the lines to show trajectories of overlapping results.

One major advantage of LMEs is their ability to account for random effects–other statistical models like random effects (RE) ANOVAs can also do this, but it’s unclear how well other statistical models that do not include random effects will perform in the presence of random effects. The presence of random effects is ubiquitous in neuroscience as many experiments are conducted across subjects, channels, trials, etc. and thus properly accounting for the influence of random effects is critical. Furthermore, the amplitude of random effects can change over time within an analysis window (e.g. **Supplementary Figure 4H**). Therefore, we simulated a set of simple distributions with a 1×4 factorial design with varying random intercepts or random intercepts as with random slopes.

For simulations with varying random intercepts amplitudes, statistical models that included random effects outperformed statistical models without random effects (**Simulation Figure 4**). For random intercepts simulations, congruence across statistical models was generally very good and increased with trial count. For simulations of 200 trials (**Simulation Figure 4A)**, congruence with the LME results was at least 96.9% for the ANOVA and permutation of the F-statistics, and for simulations with 300 trials and above congruence across all statistical models was essentially perfectly congruent (≥ 99.9%). Additionally, the permutation of the sum(*t-statistic*^2^) from the LME produced perfectly congruent (100.0%) results under all simulation parameters.

Simulations of random effect slopes and intercepts produced less congruent results (**Simulation Figure 4B/C)**. Specifically, for low trial counts all statistical models produced less congruent results with an LME as a function of random slope amplitude except the permutation of the sum(*t-statistic*^2^) which produced highly congruent results (≥ 99.2%) to the LME. All other statistical models produced less congruent (≥ 94.5%) results at 200 trials. Further, even at 500 trials the other statistical models could not fully account for higher random slope amplitudes (congruence ≥ 96.2%).

We also compared adjusted-R^2^ and model fitness as a function of random effect amplitude. While adjusted-R^2^ increased for the LME and RE-ANOVA as random intercept amplitude increased, adjusted-R^2^ decreased slightly for the ANOVA and FE Only models (**Simulation Figure 4D)**. Results were similar for the random slope simulations (**Simulation Figure 4E)**. Additionally, trial count had very little effect on adjusted-R^2^ values. Next, we compared model fitness between the FE Only models and LMEs using an LLR test (α < 0.05). As random intercept amplitude increased, the probability of LMEs being a better fit for the data than a FE Only model increased (**Simulation Figure 4F)**. Similarly, as random slope and trial count increased, the probability of LMEs being a better fit for the data than a FE Only model increased (**Simulation Figure 4G)**.

Overall, these simulations show that models that account for random effects generally perform better than models that do not though higher trial counts can make up for some of the difference. However, in the real-world collecting more data to maintain power is difficult and potentially not possible or ethical *post hoc*.

### Simple Nonlinear 1×2 Simulations

We simulated 1×2 factorial experimental designs with binomial and Poisson distributions. We did not extend our simulations to greater than 1×2 factorial designs because the downsides of certain linear statistical models (e.g. multiple separate t-tests) identified in the previous sections should also apply to the nonlinear statistical models in a similar manner.

As expected, our binomial 1×2 simulations showed that Logistic GLMEs performed similarly (congruence ≥ 91.7%) to other nonlinear statistical models (**Simulation Figure 5**). Importantly, results from the permutation of the sum(*t-statistic*^2^) were highly congruent with the results from GLMEs (≥ 96.3%), and results from the GLMEs and results from χ^2^ tests were highly congruent (≥ 95.8%) except in one simulation (87.1%) with the largest trial trial bias (0.9) and lowest trial count (100 trials). Additionally, GLMEs and χ^2^ tests were slightly more likely to produce significant results compared to Fisher’s exact tests or the permutation of the sum(*t-statistic*^2^). For the same probability between conditions across various trial counts there was a average difference in nominal significance rates: 0.0546 for the χ^2^ test, 0.0518 for the GLME, 0.0384 for Fisher’s exact test, and 0.0472 for permutation of the sum(*t-statistic*^2^). Furthermore, congruence was generally lower for borderline conditions (e.g. probabilities of 0.6 to 0.7) especially for lower trial counts while congruency was generally higher (> 99.0%) at higher trial counts (300 to 500) and larger probabilities (e.g. 0.8 and 0.9). Overall, 1×2 binomial simulation results across statistical models were slightly less congruent than the 1×2 linear simulation results. However, Fisher’s exact test was generally the least congruent and even though the nominal rate for Fisher’s exact test was not significantly different than 5% (χ^2^ proportion test), it has been argued that Fisher’s exact test is overly conservative (Crans & Shuster, 2008). Furthermore, a χ^2^ test produced highly congruent results with a Logistic GLME except at low trial counts and high trial biases where a GLME is expected to outperform a a χ^2^ test.

Similar results were obtained from 1×2 simulations of Poisson distributions (**Supplementary Figure 5**). There was high congruence (≥ 97.3%) between Poisson GLMEs and the permutation of the sum(*t-statistic*^2^) for varying trial counts and rate differences, but there was less congruence between the GLME results and a z-test on the z-transformed data (≥ 85.9%) or a ks-test (≥ 63.9%). However, ks-test results appeared substantially under powered and z-tests on the z-transformed data produced more significant results at borderline conditions (e.g. rate differences 0.5 and 1.0). For the ks-test, the false positive rate for the zero-rate-difference conditions across different trial counts was only 1.64% (χ^2^ test vs 5%, p = 4.12e-5). Congruence trends were similar for the simulations in which we varied trial biases, except the ks-test performed even worse overall. Importantly, the congruency of the GLME results were still highly congruent (≥ 97.5%) with results from the permutation of the sum(*t-statistic*^2^). Overall, these simulation results are expected. Unfortunately, Poisson GLM[E]s are the standard statistical model for Poisson distributions and finding a good comparative alternative is difficult; while z-tests on z-transformed data and ks-tests can work in certain scenarios they may produced more type I or type II errors than expected.

**Simulation Figure 5:**
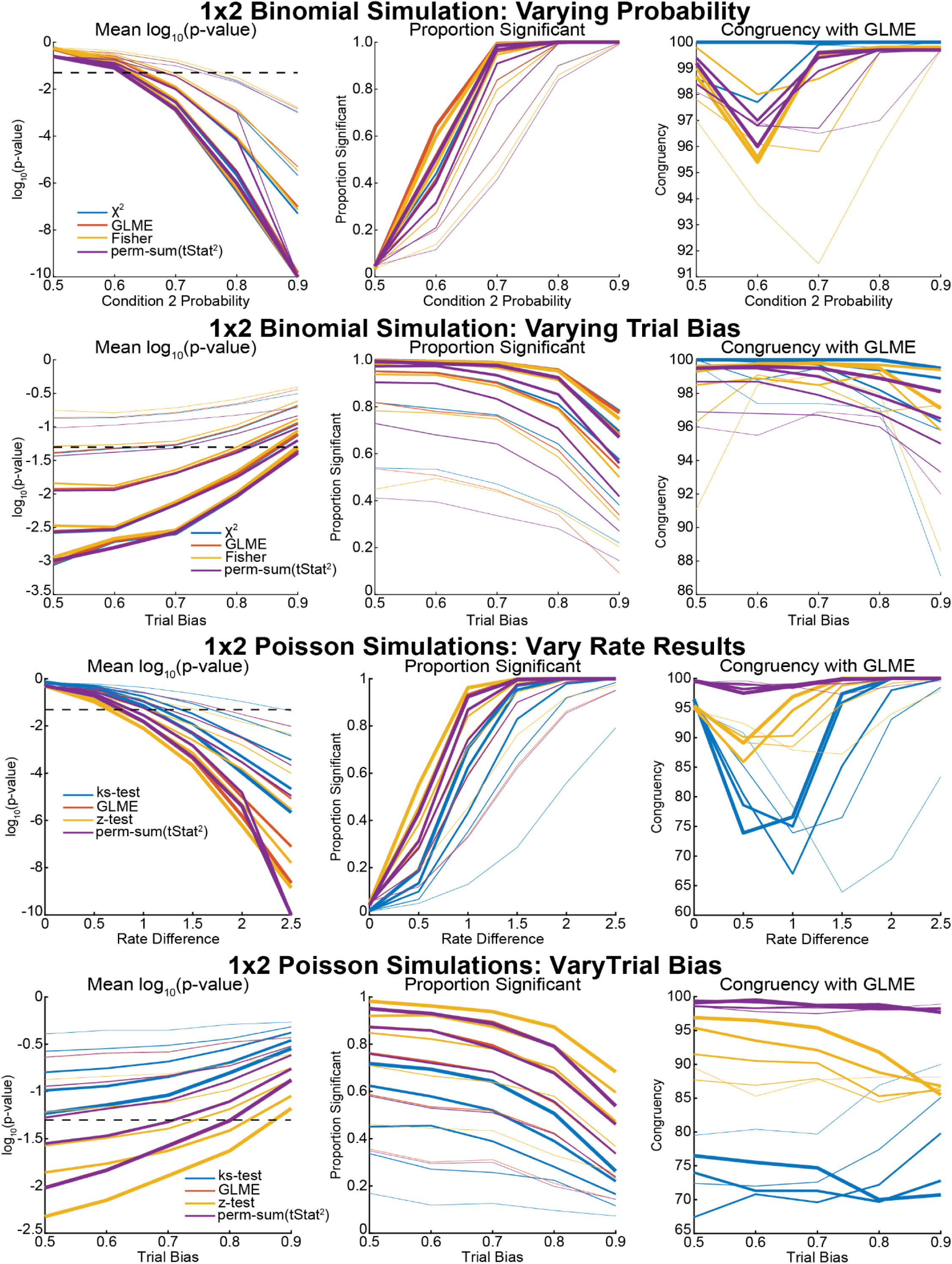
Simple Non-Linear 1×2 Simulations. Simulation results for 1×2 binomial and 1×2 Poisson distributions. For 1×2 binomial simulations, we varied trial counts from 100 to 500 (line thickness), the probability of the 2^nd^ condition, and trial biases. We compared the result of a Logistic GLME to a χ^2^ test, Fisher’s exact test, and the permutation of the sum(*t-statistic*^2^). For the 1×2 Poisson simulations we varied trial counts (line thickness), rate of the 2^nd^ condition, and trial biases. We compared the result of a Poisson GLME to a z-test, ks-test, and the permutation of the sum(*t-statistic*^2^). Note there is some added offset in the plots so that not all lines overlap. Most simulations produced maximum congruence between the GLME results and the other stats at or close to 100%. *Note small offsets were applied to the lines to show trajectories of overlapping results.

### 2×2 Simulations of Whole-Model Cluster-Level Permutation Statistics

We wanted to understand the best permutation method and whole-model test statistic to be used to explain whether a cluster fit the data better than expected by chance while accounting for any biases in the data. Specifically, our goal was to identify a robust permutation method and whole-model test statistic that had good type I and type II error control as well as being invariant to the presence of random effects. We chose several statistics from the literature as well as some alternative statistics we thought might perform well. We generated simulations with a 2×2 factorial design with varying amplitudes across all 4 groups as well as varying random intercept amplitudes.

First, we were interested in finding a permutation method and whole-model test statistic with good type I error control. All permutation methods had good type I error control in a subset of 100 simulations with zero amplitude differences across groups (**Simulation Figure 6A**). However, test statistics using R^2^ produced many false positives, except in the reduced FE Only model. All permutation methods and test statistics had false positive rates of exactly 0.05 except full model permutation tests using R^2^ (0.46) and residual permutation tests using R^2^ (0.46). More simulations would be required to determine if the false positive rate was significantly different than 0.05

We also looked at greater detail at simulations where neither fixed effect was significant (p’s > α/2, α = 0.05) according to a LME but had non-zero fixed effect sizes (|ß| > 0). As a whole, all permutation methods and test statistics resulted in higher “type I” error rates than expected for simulations with 0 significant effects (**Simulation Figure 6D**); however, when we divided simulations into those in which a mixed effects model was better than a random effects only model (LLR test, p < 0.05), the false positive rates were below α, except most R^2^ test statistics (**Simulation Figure 6C**). Further, the magnitude of the fixed effects across groups for simulations benefiting from a mixed effects model was significantly greater (ks-test, p = 1.67e-14) than simulations sufficiently explained by random effects only suggesting that there were some “subthreshold” effects in these models. Note, the prerequisite of our proposed method is that at least 1 fixed effect has a significant p-value so these simulation scenarios are unlikely to influence the results from CBPT with LMEs but nonetheless good to know.

Our simulation results also suggested that test statistics based on R^2^ were not good whole-model test statistics in the presence of random effects because false positive rates correlated positively with random intercept amplitude (**Simulation Figure 6D-E**). Generally, R^2^ is not a good measure of model fitness and therefore it is not surprising that statistics based on R^2^ performed worse than other statistics. However, R^2^ from the FE Only model appeared to perform reasonably well in a manner similar to Sum F, but as the random intercept amplitude increased, the probability of beta weight *t-statistics* for fixed effects-only model being more than 1% different than the beta weight *t-statistics* for the full LME model also increased which may lead to deleterious results in other situations as the use of mixed effects models in the presence of random effects can impact the beta weight estimates and their significance.

Second, we were interested in finding a permutation statistic with good type II error control. Most statistics, except the Freedman-Lane-style permutation tests using R^2^, had good type II error when considering simulations in which both fixed effects were significant (p’s < α/2, α = 0.05) according to a LME (**Simulation Figure 6F**); the Freedman-Lane-style permutation tests using R^2^ lost statistical power as random intercept amplitude increased. There was a slight decrease in finding a significant result with increasing random effect amplitude for Sum F and R^2^ from the FE Only model (< 2% decrease) whereas no other tests, statistics or methods were affected.

We also looked at the simulations in which only 1 fixed effect was significant according to a LME. Our goal was to find a whole-model test statistic that was relatively stable across random intercept amplitudes and could still identify most of the simulations as significant, but it’s not as easy to quantify type I and type II error rates for these simulations. We found similar results for the simulations with 1 significant fixed effect with the sum(*t-statistic*^2^) being the most stable test statistic across random intercept amplitudes and the most likely to find significant effects on average in 94.88% of simulations (**Simulation Figure 6E**); results using the sum(*t-statistic*^2^) test statistics were nearly identical (< 0.6% difference) regardless of permutation method.

**Simulation Figure 6:**
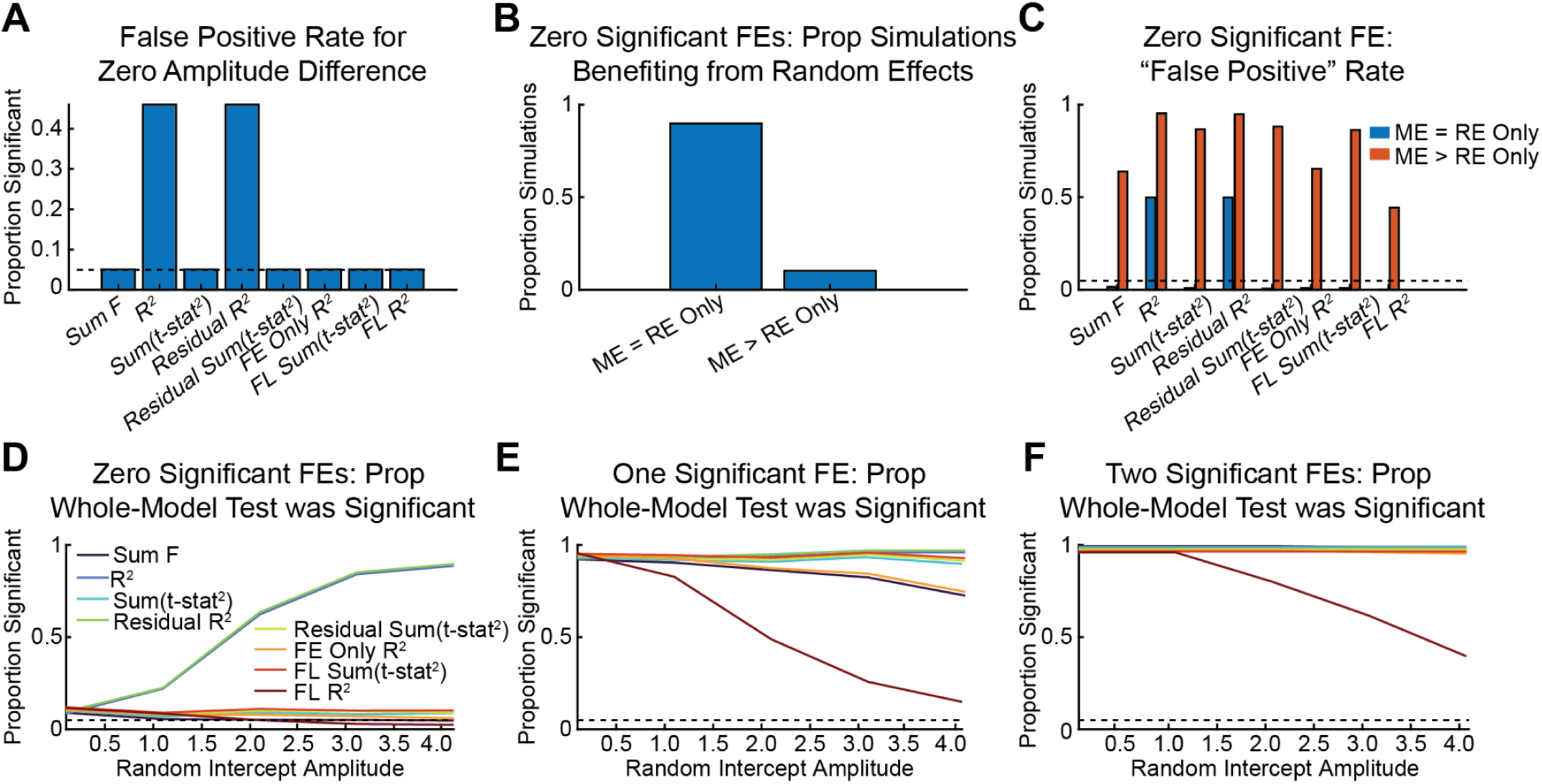
Simple Linear 2×2 Simulations of Whole-Model Test Statistics and Permutation Methods. A) False positive rate for simulations (n = 100) in which there was no amplitude difference across groups. Black dashed line indicates α = 0.05. B) Proportion of simulations with 0 significant fixed effects (FE) for which mixed effects models (ME) were better than random effect only (RE Only) models according to a LLR test. C) The proportion of simulations with significant results for simulations with 0 significant fixed effects for each permutation method and test statistic depending on whether a ME model was a better fit for the data than RE Only model. D-E) The proportion of simulations with significant whole-model results for 0 (D), 1 (E), or 2 (F) significant fixed effects as a function of random intercept amplitude. *\*Sum F is the sum of the F-statistics from a 2-way ANOVA*. *\*R^2^ is the R^2^ from the permutation of the full LME (Manly method)*. *\*Sum(t-stat^2^) is the sum(t-statistic^2^) from permutation of the full LME (Manly method)*. *\*Residual R^2^ is the R^2^ from the permutation of the full LME’s residuals (ter Braak method)*. *\*Residual Sum(t-stat^2^) is the sum(t-statistic^2^) from the permutation of the full LME’s residuals (ter Braak method)*. *\*FE Reduced Model R^2^ is the R^2^ from the permutation Fixed Effect Only Model (Manly method)*. *\*FL Sum(t-stat^2^) is the sum(t-statistic^2^) from the permutation of the full LME using Freedman-Lane-style permutation tests*. *\*FL R^2^ is the R^2^ from the permutation of the full LME using Freedman-Lane-style permutation tests*. *\*\*Lines for subplots D-F were offset slightly since results were so similar*.

### 2×2 Simulation of Proposed Cluster-Level Statistics vs. Cluster-Mass Statistics

In simulations with a 1×2 factorial design, we compared cluster-mass statics using the t-statistic from two-sample t-tests to the proposed sum(*t-statistic*^2^) from a LME modeling the mean cluster response, and cluster-mass statics using *χ^2^*-test to the proposed sum(*t-statistic*^2^) from a GLME modeling the sum of the cluster response. Our goal was to determine if the permutation results of cluster-level statistics were equivalent, because we believe using sum(*t-statistic*^2^) describes each cluster with a cluster-level model, and we believe the cluster-level sum(*t-statistic*^2^) will be less biased towards larger clusters. For these simulations we generated gaussian-like curves (**Simulation Figure 7A/C**) with a wide range of parameters (see Simulation Table 4). For the LME simulations, we found 14,147 simulations with an identifiable cluster. Of these 14,147 simulations, only 4 simulations at low amplitude conditions had different results between the cluster-mass t-tests to the proposed sum(*t-statistic*^2^) of the cluster-level data. For the GLME simulations, we found 10,123 simulations with an identifiable cluster, and of these simulations zero (0) simulations produced different results between a cluster-mass statistic using a *χ^2^*-test and the proposed cluster-level sum(*t-statistic*^2^).

Based on these simulation results, we concluded that the traditional cluster-mass statistic and the proposed cluster-level sum(*t-statistic*^2^) are essentially equivalent. Specifically, the cluster-level sum(*t-statistic*^2^) is unlikely to produce false positive results. The main benefit here is that we combine two steps in one, and we ultimately want a cluster-level model to explain the data.

**Simulation Figure 7:**
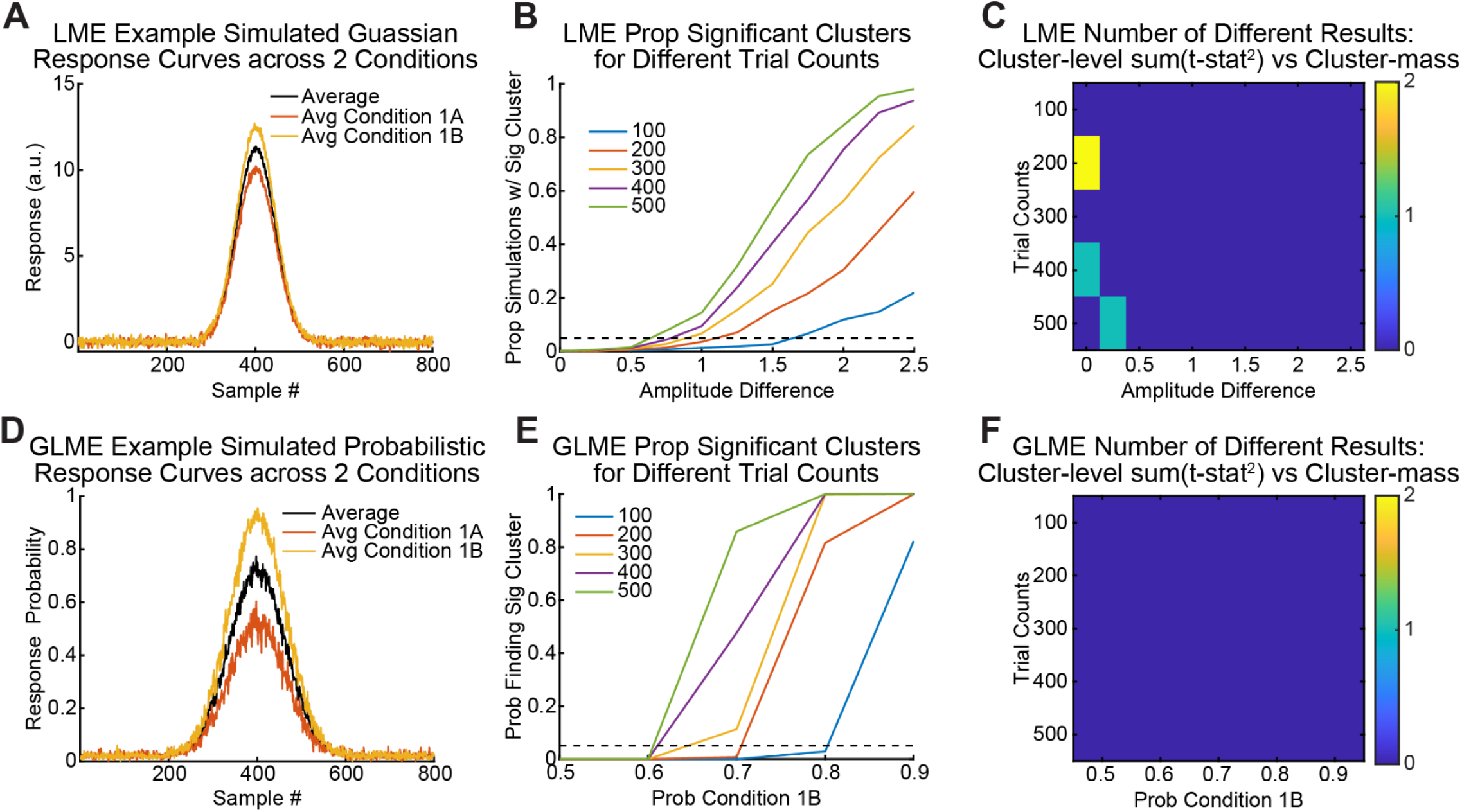
Simple 1×2 Simulations of Cluster-level Models vs Cluster-Mass Statistics. A) Example simulated gaussian response curve across 2 conditions has an apparent cluster around sample number 400. B) Proportion of simulations in which an LME found a significant cluster across different amplitudes and trial counts. C) Cluster-mass statistics from a t-test found similar results to results from proposed cluster-level sum(t-statistic^2^) with only 4 simulations at low amplitude differences having different results out of 14,147 simulations with an identifiable cluster. D-F) Same as A-C except cluster-level models were calculated from a Poisson GLME and cluster-mass statistics calculated from a *χ^2^*-test. Out of 10,123 simulations with an identifiable cluster, all simulations produced the same result.

### 4x(1×2) Simulations of Multiple Comparison Correction Methods for Analyzing Numerous Fixed Effects

**Simulation Figure 8:**
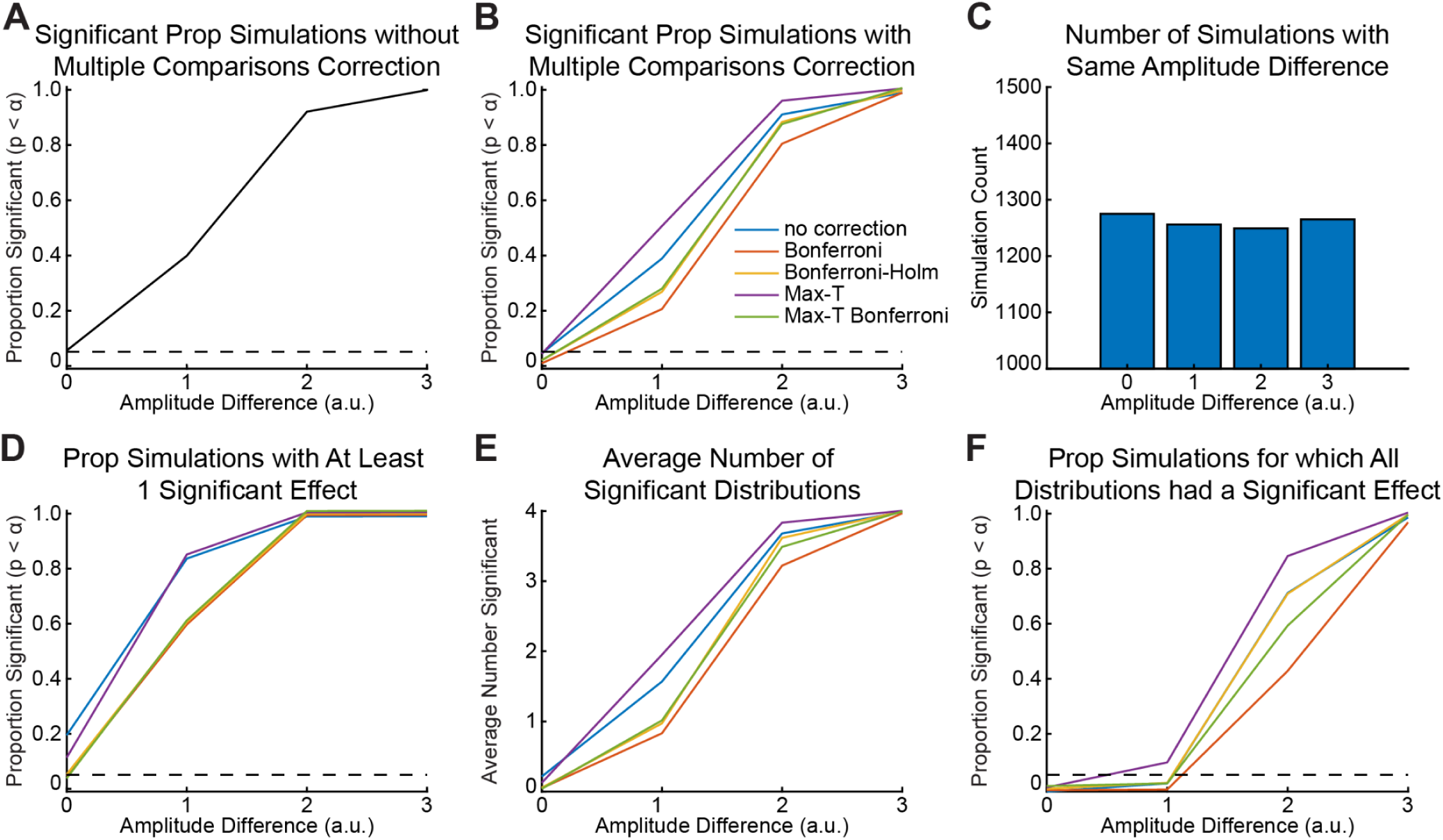
Simple Simulations of Multiple Comparisons Methods for Numerous Fixed Effects. A) The proportion of simulations with a significant result for different amplitudes with *uncorrected* p-values. B) The proportion of simulations with a significant result for different amplitudes using *corrected* p-values. C) Number of simulations in which all distributions were simulated with the same amplitude in a set. D) Proportion of simulations with at least 1 significant result after multiple comparison correction. E) Average number of distributions in a set with significant results after multiple comparison correction. F) Proportion of simulations in which all distributions in a set had significant results after multiple comparison correction.

We simulated 10,000 sets of 4x(1×2) linear distributions to determine how multiple comparisons correction methods affect FWER and statistical power when simultaneously analyzing numerous independent fixed effects.

Replicating previous simulations, the proportion of significant results increased with amplitude difference when not correcting (**Simulation Figure 8A**) or correcting for multiple comparisons (**Simulation Figure 8B).** For uncorrected p-values, 5.57% of simulations were significant at zero amplitude difference and 99.91% of simulations were significant at an amplitude difference of 3. However, for corrected p-values, the results varied across multiple comparisons methods especially at medium amplitude differences (2 & 3); in particular, having no correction or using a Max-T correction produced more significant results than the Bonferroni-Holm method while the Bonferroni method produced the fewest significant results. Note, nearly all methods produced significantly different proportions of results compared to the Bonferroni-Holm method except at zero difference and a difference of 3 because we had 10,000 simulations where even a small difference was significant (χ^2^ proportion test, most p’s < 0.05/16). Overall, Max-T with Bonferroni correction produced the most similar proportions of significant results to the Bonferroni-Holm method.

More importantly, we wanted to know the FWER for each of the multiple comparison correction methods. Since FWER is the probability of making 1 or more type I errors, we looked at the proportion of significant results for simulations in which all amplitude differences in the set were 0 with the expectation that the FWER was approximately equal to α and here α = 0.05 (**Simulation Figure 8D**). The FWER rate was 20.47% for no correction (*χ^2^* proportion test vs 5%, p = 1.29e-31), 5.18% for Bonferroni correction (p = 0.8571), 5.25% for the Bonferroni-Holm correction (p = 0.724), 10.82% for Max-T (p = 5.76e-8), and 2.82% for Max-T with a Bonferroni correction (p = 0.0043). These results suggest that no correction and Max-T correction were insufficient to control for FWER while Max-T with a Bonferroni correction was a little conservative.

We also looked at the probability of finding at least 1 significant result when all amplitude differences in a set were greater than 0 (**Simulation Figure 8D)**. When the amplitude difference in a set was 1, the results were similar for Bonferroni, Bonferroni-Holms, and Max-T with a Bonferroni correction (*χ^2^* proportion test, p’s >= 0.838), but more simulations were significant for no correction and a Max-T correction (*χ^2^* proportion test, p’s = 9.538e-43). At an amplitude difference of 2 and 3, the probability of finding at least one significant result was the same across multiple comparisons methods (*χ^2^* proportion test, p’s = 1).

We also looked at the number of distributions in a set (out of 4) that were significant across simulations with the same amplitude (**Simulation Figure 8E)**. Max-T with a Bonferroni correction appeared to produce the most similar results to a Bonferroni-Holm correction but was slightly more conservative; Max-T with a Bonferroni correction was still more liberal than a Bonferroni correction alone. No correction and a Max-T correction appeared to produce an inflated number of significant results compared to a Bonferroni-Holm correction.

Finally, we looked at the proportion of simulations in which all distributions were considered significant in a set. The Max-T appeared to produce an inflated proportion of sets in which all distributions were significant compared to all other correction methods (including none) while a Bonferroni correction produced the least. Importantly, the Max-T with a Bonferroni correction produced results in between the Bonferroni and a Bonferroni-Hom methods.

Overall, these simulation results indicate that a multiple comparisons correction is necessary to control FWER across simultaneously analyzed independent, multiple fixed effects. Specifically, not correcting for the number of fixed effects will inflate the FWER, and the FWER will grow with the number of fixed effects. Furthermore, Max-T produces an inflated number of significant results; we believe this is most likely to occur when effect sizes for each fixed effect are similar because the associated t-statistics will be similar and thus all of the t-statistic will be larger than any shuffled distribution resulting in a larger than expected proportion of significant fixed effects. As expected a Bonferroni correction is conservative while a Bonferroni-Holm correction appears to balance type I and type II errors the best. Our proposed sum(*t-statistic*^2^) is similar to the Max-T with a Bonferroni correction which was more conservative than expected when amplitude differences in the set were all 0, but otherwise produced results similar to or was only slightly more conservative than a Bonfferoni-Holm correction. Importantly, sum(*t-statistic*^2^) with a Bonferroni correction is straightforward and easily implemented throughout a multi-step hypothesis strategy whereas a Bonferroni-Holm’s correction may not be.

It is important to note these simulations do not make any statements about controlling for FWER within an individual fixed effect only across multiple, independent fixed effects. Correcting for multiple potential clusters for a given fixed effect would require us to consider possible correlations between potential clusters which we did not do here.

## Notes

https://doi.org/10.5281/zenodo.7703148

### Competing Interest Statement

The authors have declared no competing interest.

